# Short-Term Dairy Elimination and Reintroduction Minimally Perturbs the Gut Microbiota in Self-Reported Lactose Intolerant Adults

**DOI:** 10.1101/2021.10.10.463842

**Authors:** Courtney J. Smith, Les Dethlefsen, Christopher Gardner, Linda Nguyen, Marcus Feldman, Elizabeth K. Costello, Oren Kolodny, David A. Relman

## Abstract

One of the outstanding questions regarding the human gut microbiota is how interventions designed to manipulate the microbiota may influence host phenotypic traits. Here, we employed a dietary intervention to probe this question in the context of lactose intolerance. To assess the effects of dairy elimination and reintroduction on the microbiota and host phenotype, we paired fecal 16S rRNA amplicon sequencing with a clinical assay for lactose intolerance, the hydrogen breath test. We studied 12 self-reported mildly lactose intolerant adults, each with tri-weekly collection of fecal samples over a 12-week study period (2 weeks baseline, 4 weeks of dairy elimination, 6 weeks of gradual milk reintroduction) and a hydrogen breath test before and after each phase. We found that although none of the subjects experienced a change in clinically defined status of lactose intolerant or tolerant, most subjects were qualitatively better able to tolerate dairy products with minimal symptoms at the end of the study compared to their baseline. Like clinical status, gut microbiota also resisted modification. Although the mean fraction of the genus *Bifidobacterium*, a group known to metabolize lactose, increased slightly with dairy reintroduction (from 0.0125 to 0.0206; Wilcoxon *P* = 0.068), the overall structure of each subject’s gut microbiota remained highly individualized and largely stable. Our study is small, but it suggests the possibility of qualitatively improved tolerance in the absence of change in clinically defined tolerance nor major change in the gut microbiota.

## Introduction

Lactose is a disaccharide and the main carbohydrate found in mammalian milk. Lactase is an enzyme that allows utilization of lactose from milk by cleaving lactose into glucose and galactose since disaccharides are poorly absorbed. To utilize the lactose in mothers’ milk, mammals produce lactase at the brush border of the jejunum in infancy and normally stop expressing this enzyme shortly after the age of weaning. Undigested lactose passing through the small intestine into the colon can cause lactose intolerance, which manifests as diarrhea, abdominal discomfort, bloating and flatulence following consumption of dairy products. These symptoms are likely the product of the osmotic load of undigested lactose in the colon, as well as gas and other metabolites from bacteria fermenting the lactose.^1,2^

In humans, genetic adaptations occurred following the domestication of cattle, causing approximately 35% of the world’s population to continue expressing lactase into adulthood, referred to as lactase persistence.^3–8^ Lactase persistence confers lactose tolerance, but tolerance can evidently be conferred by other mechanisms as well: many lactase non-persistent individuals consume dairy products regularly without reporting any symptoms.^1,4,9,10^ One potential mechanism of lactose breakdown in lactase nonpersistent individuals is by lactase-producing microbes in the gut. In humans, genetic variation within the locus that includes the gene LCT, which encodes the enzyme lactase, has been shown to have an age-dependent, genome-wide significant association with abundance in the gut microbiota of the genus *Bifidobacterium*, a group known to metabolize lactose, even though host genetics have a minor role in determining microbiota structure relative to other environmental factors such as diet.^11,12^

The lactose hydrogen breath test is the primary diagnostic test for clinical lactose intolerance and measures the concentration of hydrogen and methane in the breath of patients in the hours following the consumption of a standardized dose of lactose (typically 25 grams, or the equivalent of about two cups of milk).^13,14^ The negative effects of dairy product consumption frequently lead to avoidance of dairy products by intolerant individuals. Such avoidance has become a widespread recommendation for these individuals by medical practitioners and is growing in popularity among large segments of Western society. However, the regular consumption of dairy products (and the associated calcium intake) is linked to positive health outcomes, such as a reduced risk of osteoporosis and bone fractures.^15,16^ In addition, previous studies suggest that lactose tolerance can be acquired by lactose intolerant individuals over a few weeks by the inclusion of regular, or regularly increasing amounts of lactose in the diet.^1,17–22^

The importance of the role of the microbiota in health and disease is increasingly evident. Colorectal cancer, inflammatory bowel disease, and obesity are examples of conditions to which the gut microbiota contributes.^17,23^ In the last decade, researchers have begun exploring this relationship more deeply through intervention studies designed to discover whether the structure of the gut microbiota can be manipulated in such a manner as to influence host phenotype.^24,25^ Prior studies have suggested that lactose tolerance can be acquired via gradual introduction of dairy products into the diet, but little is known about the microbiota changes that are presumed to accompany and facilitate acquired tolerance.^5,18,26,27^ Therefore, we examined the changes in gut microbiota structure brought about by a dietary manipulation of dairy product consumption. To date, changes in overall human gut microbiota structure in response to dietary changes regarding the consumption of dairy products have not been well-studied.

Here, we combined gut microbiota data with lactose hydrogen breath test results to assess the responses of the microbiota to dietary manipulation of dairy product consumption. Our results demonstrate a surprising level of resistance to perturbation in response to the dairy product interventions, both in terms of lactose tolerance and gut microbiota structure.

## Results

### Overview of study design and sample collection

We studied 12 self-reported mildly lactose intolerant subjects (see **Supplementary Table 1** for subject information), each with tri-weekly collection of fecal samples over a 12-week study period (2 weeks baseline, 4 weeks of complete dairy elimination, 6 weeks of gradual milk reintroduction) and a hydrogen breath test (HBT) before and after each phase (**Figure 1**). We recruited individuals who self-reported being lactose intolerant to enrich for lactase non-persistent study subjects, but 6 subjects despite self-reporting as intolerant still consumed some dairy products.

**Figure 1.**
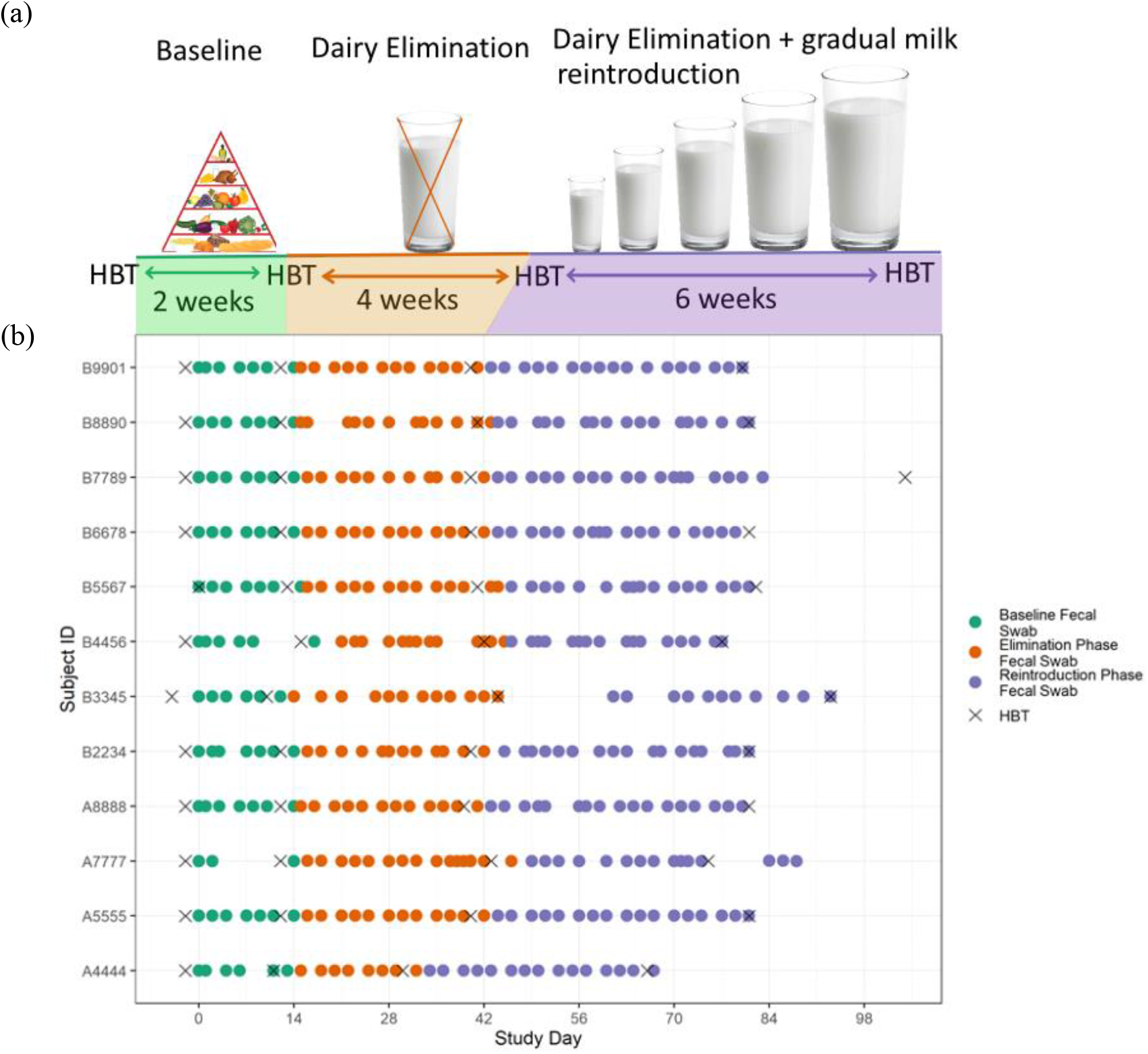
The study consisted of tri-weekly fecal sample collection and three diet phases with a HBT before and after each phase. (a) Overview of 12-week study design. (b) Sample collection (day 0 was defined as the day on which the first fecal sample was collected for that subject). During the two-week baseline phase subjects were instructed to maintain their normal diets. During the following four-week elimination phase subjects continued their normal diets but avoided all dairy products. During the six-week reintroduction phase subjects were instructed to continue their same diet, and in addition, to follow a specific protocol for gradually increasing consumption of milk, working up to two cups of milk a day during the last week. A HBT was performed before and after each phase and fecal swabs were collected three times per week.

Throughout the twelve weeks of the study, the severity of subjects’ lactose intolerance was evaluated with two metrics. The first was based on the HBT: we measured the combined concentration of hydrogen and methane gas in each breath sample, the standard measurement in the clinic, collected for each subject over a six-hour period after drinking two cups of milk. Each subject completed four HBTs at standard timepoints during the study, enabling temporal analysis within subject as well as between subjects. The second metric was based on self-reported symptoms recorded in a daily log (**Supplementary Table 2**) throughout the study and every half-hour during each HBT.

### Clinical status at baseline and in response to intervention

At baseline, all subjects self-identified as mildly lactose intolerant and 6 of 12 reported regularly consuming some amount of dairy products in their diet, while the others abstained from dairy products completely or used lactose-free products. We hypothesized that subjects who regularly consume some dairy products in their diet might lose or see reductions in the relative abundances of lactose-utilizing bacteria during the dairy elimination phase, thus becoming somewhat more lactose intolerant compared to baseline, and that subjects might increase lactose tolerance during the reintroduction phase if the milk consumed during this phase were sufficient to increase the growth of specialized bacteria that metabolize lactose with few ill-effects for their host.

Each HBT (**Figure 1**) included 13 breath samples taken at 30-minute intervals over six hours, where the combined concentration of hydrogen and methane gas of each sample could be plotted against time since milk consumption at time zero (**Supplementary Figure 1**). Subjects are classified by the HBT as clinically intolerant of lactose if the concentration at any of these time points is greater than 20 ppm above the concentration for that individual at time zero. At baseline, the HBT classified 8 of the 12 subjects as lactose intolerant, despite all 12 self-reporting lactose intolerance, and none of the subjects changed their clinical status during the study. Towards the development of a quantitative measure of lactose tolerance rather than relying on a dichotomous clinical classification, we then calculated the change in area under the HBT curve (AUC) over time as this reflects the total hydrogen and methane gas concentration over the six hour collection period for each HBT. We observed a significant increase in AUC between the second baseline HBT and the HBT after the dairy elimination phase (Paired Wilcoxon signed-rank test; p=0.0015) (**Figure 2; Supplementary Figure 2**). Of the 12 subjects, 11 had a higher AUC in the HBT after the dairy elimination phase than in the second baseline HBT, including all 8 subjects with clinically defined lactose intolerance (Paired Wilcoxon signed-rank test; p=0.008). We also observed a significant increase in AUC between the second baseline HBT and the HBT after the reintroduction phase (Paired Wilcoxon signed-rank test; p=0.021). Of the 12 subjects, 9 had a higher AUC during the HBT after the dairy reintroduction phase than during the second baseline HBT, including 7 of the 8 subjects with clinically defined lactose intolerance (Paired Wilcoxon signed-rank test; p=0.023).

**Figure 2.**
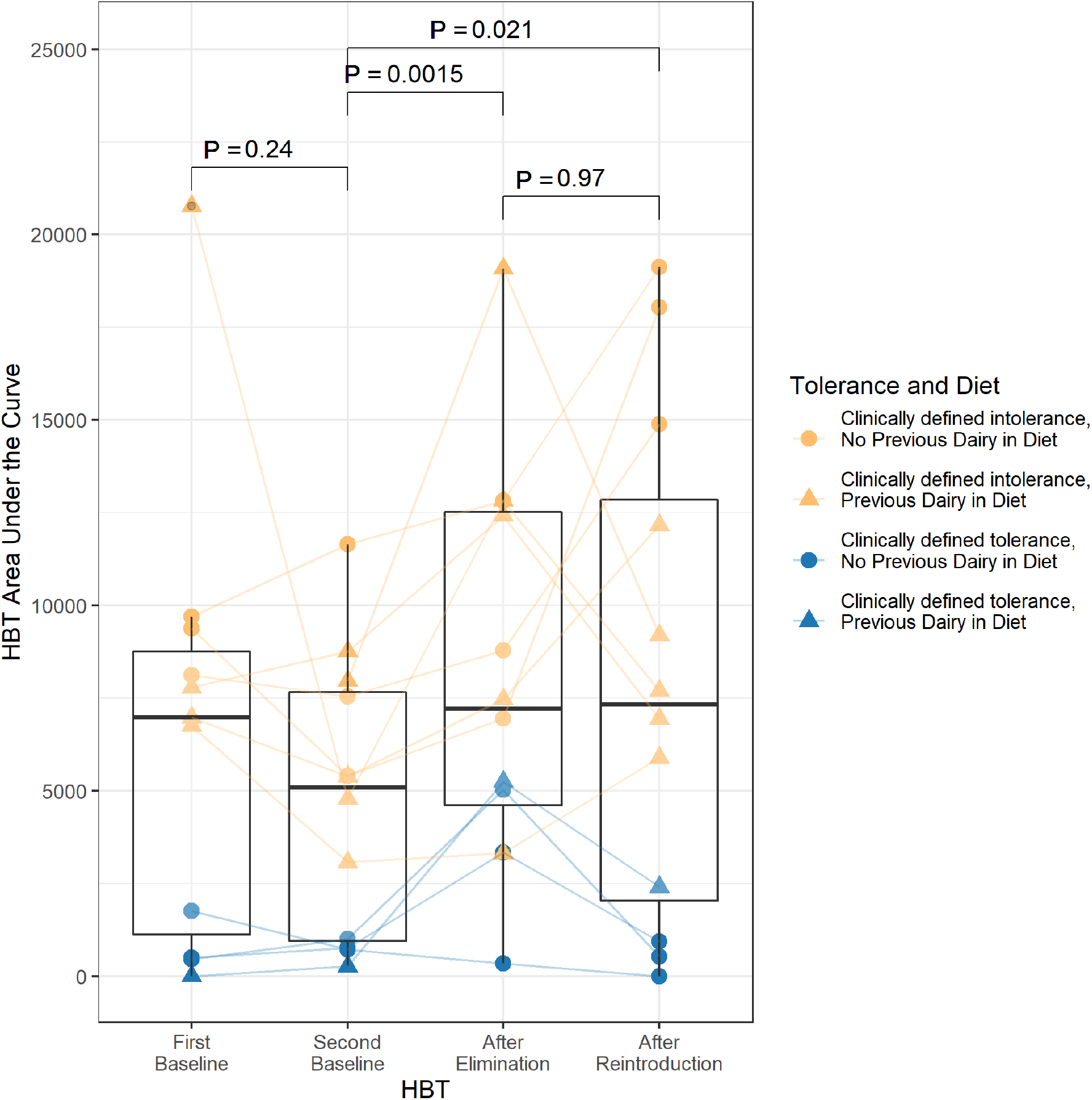
Area under the curve (AUC) of the HBT for each subject across the study. AUC significantly increased after the dairy elimination phase, relative to the second baseline just before this phase, potentially reflecting an increase in intolerance—despite no change in clinical classification. After the dairy reintroduction phase, some subjects increased further while others seemed to recover. Coloring indicates the clinically defined intolerance status of that subject based on HBT results, and shape indicates whether subjects were dairy product abstainers (circles) or consumers (triangles) prior to the study. See **Methods** for further information about calculation of HBT area under the curve. Paired samples Wilcoxon signed-rank test p-values are shown.

However, based on the self-reported symptom data, the majority of subjects reached high qualitative self-reported lactose tolerance by the end of the dairy reintroduction period relative to the baseline period, revealing an unexpected discordance between clinical lactose tolerance as assessed by HBT and self-reported lactose intolerance defined by symptoms (**Supplementary Table 3**). Nine of 12 subjects (including 5 of the 8 subjects defined as clinically lactose intolerant by the HBT) were able to tolerate two cups of milk daily in the final week of the study without any reported symptoms (except mild gassiness), and the symptoms reported by the others over this phase were not severe.

### Subject identity determines microbiota structure over and above temporal factors, including the dietary intervention

We next investigated changes in the microbiota structure of subjects throughout the study and in response to the dietary intervention using 16S rRNA gene sequencing. **Figures 3a and 3b** display the first two axes of the principal coordinates analysis (PCoA) of all samples across the study timeline and all subjects, using binary Jaccard dissimilarity measures. Samples cluster predominantly by subject ID (**Figure 3a, Supplementary Figure 3a**) over and above any clustering by study phase (**Figure 3b, Supplementary Figure 3b**), indicating that stable differences between individuals are much more pronounced than any changes in each subject’s microbiota over the course of the study (Permanova with 1000 permutations on Jaccard distances; by subject ID: R^2^ = 0.758, p-value<0.001; by study phase: R^2^ = 0.004, p-value=0.707). Even when comparing the overall microbiota structure of an individual to themselves throughout the study though, we did not identify any consistent shifts in microbial alpha diversity or beta diversity in response to the elimination or reintroduction of dairy products (**Figure 3c, Supplementary Figure 3c, Supplementary Figure 4**). There was significant clustering of samples by study phase nested within each subject, but only a small amount of variation in the distance matrix was explained by this (Permanova with 1000 permutations on Jaccard distances by study phase nested within subject ID: R^2^ = 0.032, p-value<0.001).

**Figure 3.**
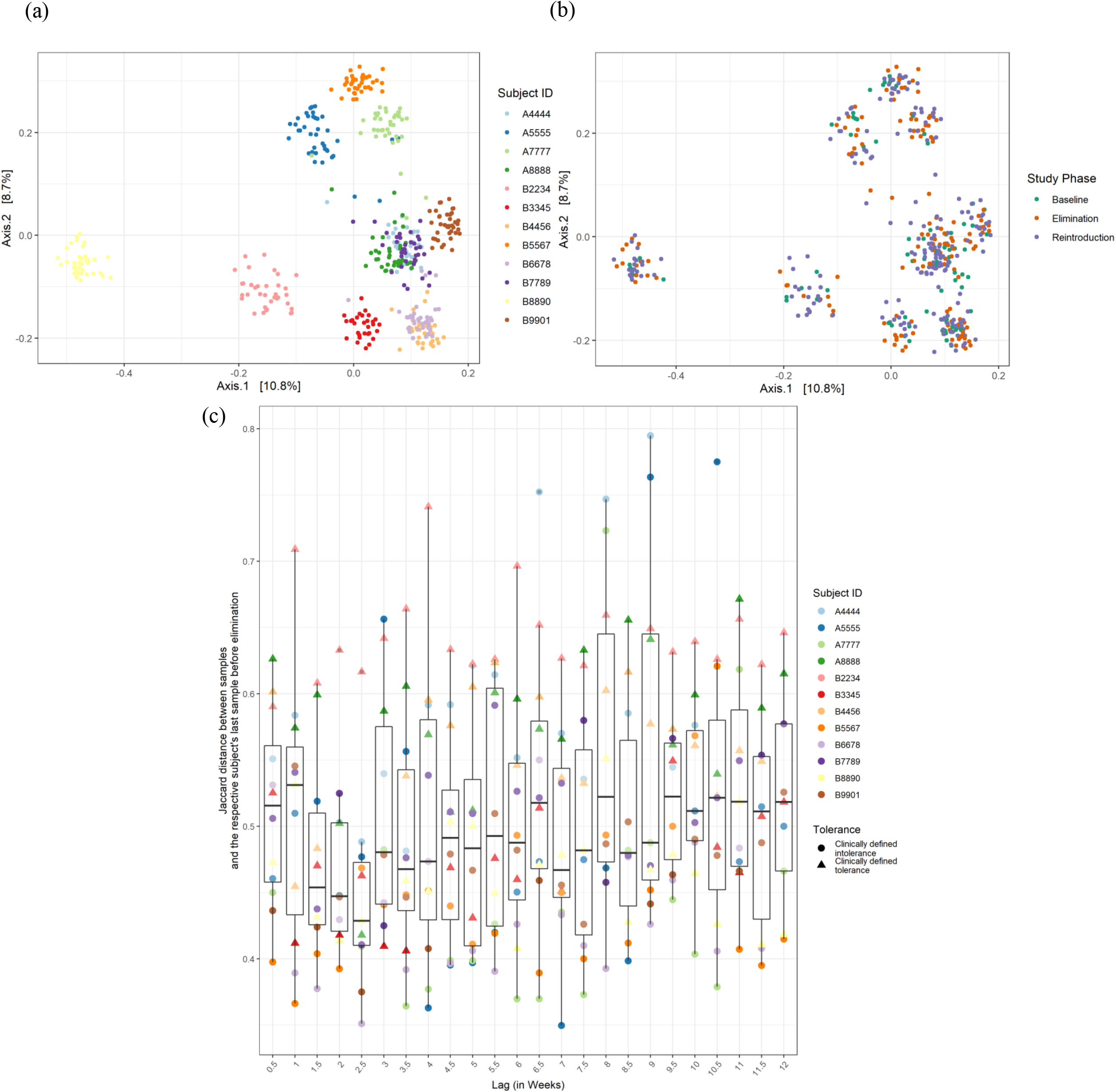
Individual-specific overall microbiota structure was minimally perturbed by the dietary intervention. Visualization of the first two principal coordinates from PCoA across all samples for all subjects based on Jaccard index distance, colored by subject (a) and by study phase (b). Samples throughout the study clustered predominately based on subject ID. (c) The beta-diversity (Jaccard index distance) between each subject’s last sample before dairy elimination and that subject’s samples other time points. The samples are placed in half-week bins with a maximum of one sample from each subject included in each bin. The half week bins correspond to the number of days between when the samples in each sample pair were collected (lag), not the study day. If more than one sample for a given subject was collected within a given half-week bin, one sample was randomly selected to be the one included. Not every bin has every subject due to occasional incidental variations in sampling frequency by some subjects. See **Supplementary Figure 5** for corresponding plots using Bray-Curtis distance metrics.

### Subtle microbiota changes in response to the dietary intervention are consistent across a few individuals

The absence of a strong, consistent intervention-associated shift in overall subject microbiota structure did not rule out the possibility that the interventions had a more subtle effect, e.g., on individual microbial taxa. We next investigated whether there were specific taxa for which a consistent shift in abundance across subjects was observed. To identify such shifts between study phases, we performed a supervised linear discriminant analysis using TreeDA.^28^ This package performs sparse discriminant analysis using a phylogenetic tree structure provided to the algorithm. The necessary inputs are the classes to be discriminated (our elimination vs. reintroduction phase samples), a set of predictors (the taxa abundances identified in our data), and a tree describing the relationship between the predictors (the phylogenetic tree of the microbial taxa). TreeDA identified *Bifidobacterium*, along with *Ruminococcus_2* and *Agathobacter* as key predictor genera that encompass many of the amplicon sequencing variants (ASVs) useful for distinguishing samples in the elimination phase from those in the reintroduction phase. While the specific genera prioritized by TreeDA were sensitive to the input parameters, this result suggests that there may be a consistent and interpretable microbiota structure shift across subjects (see **Methods**). The genus *Bifidobacterium*, for example, has previously been implicated in lactose tolerance and many species within this genus have the ability to break down lactose.^18,27^ However, increased *Bifidobacterium* abundance has also been implicated in lactose intolerance.^29^

Following up on this result, we investigated the change in ASVs assigned to the genus *Bifidobacterium* abundance across subjects’ microbiota over the course of the entire study. **Supplementary Figure 3** shows that there was no obvious decrease in the abundance of *Bifidobacterium* in most subjects during the elimination phase, or an obvious increase during the reintroduction phase as might have been expected. There was a trend towards increased abundance of *Bifidobacterium* between the last three weeks of elimination and the last three weeks of reintroduction in several subjects but the increase in mean fraction of *Bifidobacterium* across all subjects from 0.0125 to 0.0206 was not statistically significant (Wilcoxon signed rank test p-value = 0.068, **Figure 4**). Evaluation of the relationship between *Bifidobacterium* abundance and other lactose intolerance metrics can be found in **Supplementary Figures 6 and 7**; however, no significant correlation was found. For example, when we evaluated the relationship between the area under the curve of the HBT measurements and the abundance of *Bifidobacterium*, the correlation was not significant (r = −0.074, p = 0.625) (**Supplementary Figure 7**).

**Figure 4.**
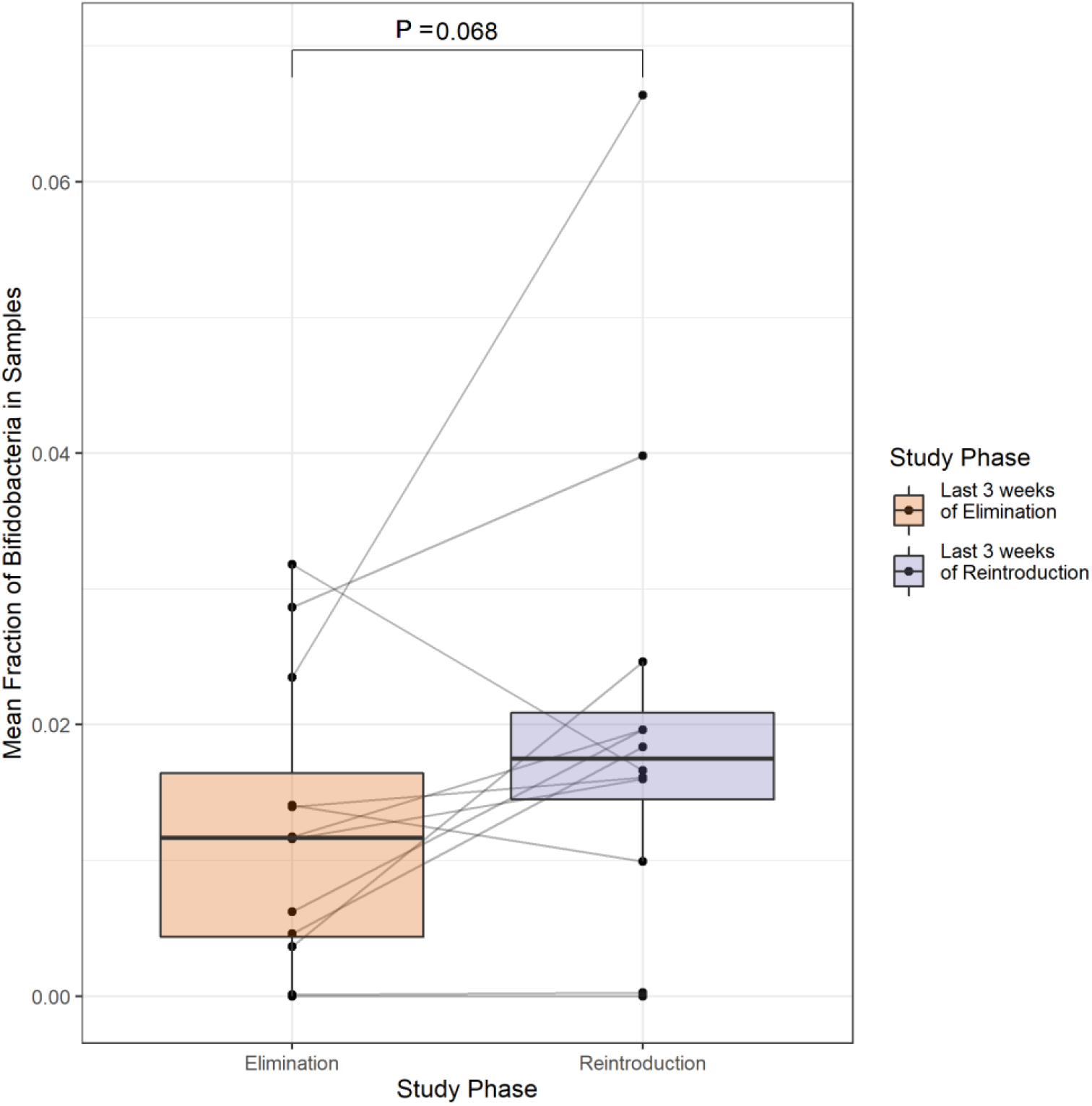
Positive but non-significant trend in relative *Bifidobacterium* abundance after dairy reintroduction. Mean fraction of *Bifidobacterium* in samples from the last three weeks of the elimination phase compared to the mean fraction in samples from the last three weeks of the reintroduction phase across all subjects.

### Microbiota structure shifts within a tightly constrained and highly individualized space even in the face of significant change in metabolic output

**Supplementary Figure 8** shows weak correlation between the first principal coordinates from the PCoAs in **Figure 3a** and time. We then wondered whether individuals with greater gut microbiota variability over time had greater HBT variability among tests. To test this, we computed the correlation between subjects’ microbiota dispersion, as measured by each subject’s average Bray-Curtis distance to the median, and the variance of their HBT areas under the curve; the correlation was not significant (r = 0.343, p-value = 0.275).

We thus did not find evidence that temporal dynamics or dietary variability are major determinants of variation in microbiota structure; on the contrary, there were surprising levels of resistance to perturbation in subjects’ microbiota structure. An individual’s microbiota structure can be viewed as located in a multidimensional space of possible microbiota structures. A conservative proxy for the range of structures within this space that are feasible among healthy individuals is the range spanned by the samples of all subjects in the study. **Figure 3** suggests that each individual’s microbiota is individualized throughout the study and tightly constrained to a small subsection within this range. To quantify the dynamics of change over time in each individual’s microbiota, we investigated the microbiota’s *autocorrelation time*, i.e. the time interval between two samples from a subject across which the difference in microbiota structure approaches the difference between randomly chosen pairs of samples from that individual. We defined autocorrelation time as the typical number of days that it takes until two consecutive samples from the same subject have the same distance between them as 90% of the median distance between pairs of randomly chosen samples from that subject.

We found an autocorrelation time of 5.25 days (SD ± 2.72 days) using the Jaccard distance metric. Our interpretation is that the typical distance in the composition space that the microbiota covers within a week approaches the median distance between any two points in the constrained region that characterizes the individual’s microbiota throughout the experiment (see **Methods** for comparison with same day control swabs). This rapid alteration of microbiota structure *within* each individual’s constrained region contrasts sharply with the restricted and stable nature of these regions. Importantly, these characteristics held both within and across study phases, highlighting the resistance to perturbation of each individual’s microbiota structure, and was consistent across individuals **(Figure 5)**. Sample pairs taken further apart than the autocorrelation time show little to no correlation between the Jaccard distance or the Bray-Curtis distance between them and the number of days apart that they were collected (**Supplementary Figures 9 and 10**). While overall short-term relative resistance to perturbation of microbiota structure is consistent with previous studies,^30,31^ the short autocorrelation period was surprising: even in the absence of the dietary intervention, a sample from a given individual is expected to be somewhat more similar to other samples collected from that individual in temporal proximity than to samples collected at more distant times. In addition, **Figure 2** demonstrates that metabolic output from the gut in the form of hydrogen and methane gas varied widely throughout the study for many subjects.

**Figure 5.**
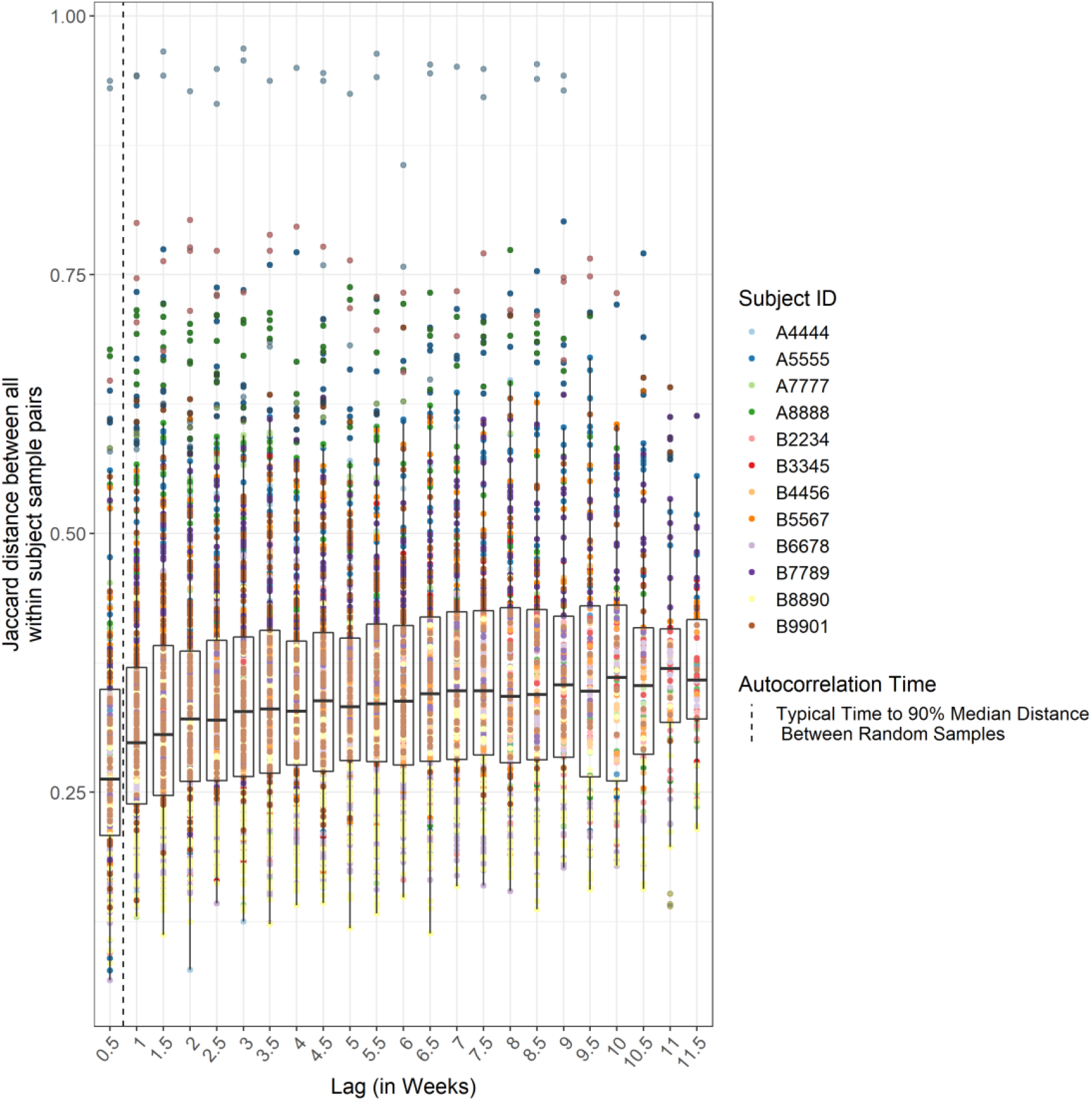
Beta-diversity of microbiota shows rapid change over time. Jaccard index distance between every sample pair from the same subject throughout the study is plotted against time interval in weeks. See **Supplementary Figure 11** for plot with Bray-Curtis index distance.

## Discussion

We have investigated how self-reported lactose intolerance symptoms, lactose intolerance clinical diagnostic test results, and gut microbiota structure vary in response to a complete dairy elimination and subsequent reintroduction. For the lactose intolerance symptoms and test results, we found that 8 of 12 subjects were clinically lactose intolerant and remained that way throughout the study; however, 10 of 12 subjects had only mild to no symptoms while drinking two cups of milk per day by the end of the dairy reintroduction period. This is consistent with prior literature that reported examples of subjects tolerating two cups of milk with only mild symptoms despite being defined as clinically lactose intolerant by the HBT.^32^ The discrepancy between the HBT results and the self-reported symptoms suggests that the clinical test used to diagnose lactose intolerance may tell only part of the story. Therefore, it could be useful to investigate alternative routes of action for patients experiencing lactose intolerance symptoms after dairy product consumption, such as making sure some dairy products are regularly kept in their diet or gradually reintroducing them after an intermission. The increase in HBT area under the curve suggests four weeks of complete dairy elimination may increase intolerance. The HBT area under the curve offers an interesting alternative to classification of lactose intolerance, because it offers more nuanced information than the binary results of the current clinical assessment. However, although we do see a potentially meaningful trend in the results, the lack of correlation with symptoms suggests that future research is still required to better understand the utility of this metric and the nature of its relationship to experienced symptoms.

Each subject’s gut microbiota remained within a tightly constrained region of the composition space (**Figure 3**), and pairs of samples collected as close together as within a week were close in similarity to samples collected months apart (**Figure 5**). The gut microbiota is often described as constantly changing, on a broad range of time scales, both in response to a plethora of external drivers and as a result of random species’ drift.^33,34^ An individual’s microbiota can thus be viewed as engaged in a random or pseudo-random walk in the space of possible microbiota structures. The nature of this walk, however, is an open question. Intuitively one might imagine, for example, that since the microbiota is constantly changing, similarity between two samples should be a function of the time between their collection: samples collected from an individual in the same week would be more similar to one another than samples collected a month apart, which would in turn be more similar than samples collected three months from one another. Alternatively, it could be the case that the microbiota structure is constantly changing, but that forces are at work that prevent it from straying too far from an anchor point which functions as an attractor in the composition space for that individual’s microbiota. Our results support the latter model as a better description of the human gut microbiota, at least on the time scale of our experiment.

Determination of the factors that anchor an individual’s microbiota structure is key to understanding its dynamics and to designing informed individualized microbiota manipulations in the future. It has been highlighted in previous studies that dietary interventions may cause dramatic shifts in microbiota structure.^34–38^ Our results stand in contrast to these previous observations: in our study relatively dramatic dietary changes – complete elimination of dairy and later reintroduction of dairy up to consumption of more than 2 glasses of milk per day – caused surprisingly limited change. In particular, these changes did not draw the subjects’ microbiota structures beyond the regions that they occupied in the composition space prior to the perturbations and did not even limit the compositions to subsections within those regions. One possible reason for this contrast may be related to the nature of the dietary change, possibly including both the subtlety of dietary intervention and the nutrient categories affected.^37^ For example, manipulation of fiber intake, a major influence on the nutrient profile that reaches the gut, might have a greater impact on microbiota structure than dietary changes that focus on other products such as dairy.^39,40^ Some dietary interventions may even indirectly affect nutrient categories other than that being directly manipulated, such as an increase in fiber resulting from elimination of meat products and subsequent increase of plant products in their place.^41^ The large discrepancy in the extent of microbiota changes caused by dietary interventions, even dietary interventions that we perceive as similarly dramatic, highlights the need for mechanistic understanding of the drivers of change in these complex systems and the necessity for a better understanding of the ecological principles that underlie the structure of the microbiota.

The difference between the effects of our dietary intervention and those in other studies also hints at the possibility that there has been some publication bias with respect to reports about drivers of change in microbiota structure: it is conceivable that different interventions have been previously explored and led to little change in microbiota structure, a result that might be less likely to be published. Our findings emphasize that publication of such results, often viewed as “negative results” is crucial to making progress in the understanding of the microbiota’s behavior over time and its response to external perturbations.

This study may have been limited by the insensitivity of fecal samples to microbial activity confined to the small intestines or by the insensitivity of 16S rRNA-based analyses to changes in microbial metabolic activity.^34^ Future studies may be better equipped to detect changes from similar interventions in larger cohorts, other types of microbiota or host physiological activity measurements, longer elimination and reintroduction phases, or a means of sampling the microbiota of the small intestines.

To the best of our knowledge, this is the first study analyzing human microbiota data alongside a noninvasive quantitative biochemical assay to assess the effect of an intervention on the gut microbiota structure and host phenotype. Despite the surprising level of resistance to perturbation in response to the dairy interventions in terms of clinically defined lactose intolerance status and of overall gut microbiota structure, future studies may benefit from coupling of such complementary approaches to probe how interventions designed to manipulate the microbiota may influence phenotypic traits of the host, especially those relevant to human health.

## Materials and Methods

### Ethics Statement

The research was approved by an Administrative Panel for the Protection of Human Subjects (Institutional Review Board) of Stanford University (protocol 42241). All subjects were properly informed of the risks and benefits of this study, and then signed an approved, written consent form.

### Experimental Design

The response of the human gut microbiota to dairy elimination and reintroduction was evaluated by collecting swabs of fecal samples three times per week from twelve healthy subjects for twelve weeks. The twelve weeks were divided into three dietary phases: two weeks of their normal diet, which is referred to as baseline; four weeks of complete dairy elimination, referred to as elimination; and six weeks of controlled gradual reintroduction of whole milk, referred to as reintroduction. Details of the reintroduction protocol can be found in **Figure 1**. All subjects used the same type and brand of milk throughout the study.

The response of the clinical tolerance of lactose was evaluated with a HBT, which is the clinical standard for evaluation of lactose intolerance, before and after each diet phase (**Figure 1**). Twenty-four hours before each HBT, the subjects followed a strict diet and fasting protocol as commonly used in the clinic as HBT preparation to limit foods that may linger and produce delayed gas. On the morning of each HBT, subjects collected a single breath sample using an at-home collection kit from QuinTron Instrument Company, West Milwaukee, WI. They then drank two cups of whole milk. At every half-hour over the next six hours they collected an additional breath sample and recorded the severity and type of symptoms they experienced, when applicable. Throughout the study subjects recorded all dairy product consumption, major deviations in lifestyle, lactose intolerance symptoms and any compliance issues. Two aspects of this HBT protocol differed from the standard procedure in the clinic. First, the duration of the sample collection period was extended to six hours from the standard of up to three hours, motivated by our interest in quantifying lactose intolerance beyond a binary classification of intolerant or not. Second, two cups of milk were used in place of the standard procedure of adding a 25 gram lactose powder (approximately equivalent to the amount of lactose in 2 cups of milk) to water to evaluate lactose intolerance in the context of dairy products.

DNA was extracted from the stool samples and used for amplicon sequencing of the V4-V5 region of the 16S rRNA gene. The data were analyzed to reveal community structure, in an attempt to characterize the gut microbiota’s response to dairy elimination and reintroduction, and to assess the correlation of these microbiota structure changes with changes in lactose intolerance as reflected by the HBT and reported symptoms. The HBT and symptoms data were also analyzed to assess their response to this dairy elimination and reintroduction protocol in these subjects. Self-reported symptoms were quantified on a scale of 0-4, with 0 corresponding to no symptoms, 1 corresponding to very mild symptoms such as gassiness, 2 corresponding to experiencing mostly mild symptoms but occasionally moderate symptoms such as cramping or other abdominal pain, 3 corresponding to moderate symptoms, and 4 corresponding to severe symptoms such as diarrhea.

### Subjects and Sampling Protocol

Healthy nonpregnant adults with self-reported mild lactose intolerance were recruited from the Stanford community and nearby area, excluding individuals with chronic disease, hospitalization or antibiotic use in the previous 6 months, immunizations, or international travel in the previous 4 weeks, or routine use of any prescription medication except birth control or hormone replacement therapy. Characteristics of the twelve subjects who completed the sampling protocol are summarized in **Table 1**. Subjects collected two swabs of each stool sample at home, which were frozen immediately in *RNAlater* in home freezers. Samples were transferred without thawing to −80°C storage in the laboratory approximately within a week of when subjects completed the study.

A total of 1008 stool swab samples were collected; the timing of samples throughout the study for each subject is shown in **Figure 1**. Some intended daily samples were not collected because subjects did not produce stool that day, in which case samples were collected at the next stool sample opportunity. 16 subjects enrolled in and began the study, but four subjects did not complete the full dietary interventions due to reasons unrelated to the study and were thus excluded from analysis.

### Sample Processing and DNA Extraction

All chemicals, solvents, and reagents were purchased from Sigma-Aldrich unless otherwise noted. Half of the collected swab samples, one of the two swab samples collected each collection day, were processed for sequencing and the other half remained in the −80°C for storage and used in case of contamination during the sample preparation process. Swab samples were thawed to room temperature during a 10 min centrifugation at 10000 rpm then transferred to bead tubes from the MP Biomedicals (Irvine, CA) Lysing Matrix E kit. Extraction was then performed with the Qiagen AllPrep DNA/RNA 96 Kit following the manufacturer’s protocol after homogenization for 1 minute at speed 6.5 using MP Biomedicals FastPrep-24 5G Instrument followed by 5 min at 4 °C and 10 minutes of centrifugation at 15000 rpm. Five extraction control blanks were included per extraction plate.

### 16S rRNA Gene Sequencing

Primers were purchased from Integrated DNA Technologies (Coralville, IA). The V4-V5 region of the 16S rRNA gene was amplified for sequencing using 515F and barcoded 926R primers (515F Forward Primer: GTGYCAGCMGCCGCGGTAA, 926R Reverse Primer: CCGYCAATTYMTTTRAGTTT).^42^ Triplicate 25 μL PCR reactions using Hot MasterMix (5 Prime) with 2 μL extracted DNA as template and 10μg/μL BSA were cycled as follows: denaturation at 94°C for 3 minutes, 25 cycles of 94°C for 45 seconds, 52°C for 60 seconds, 72°C for 120 seconds, final extension at 72°C for 10 minutes. PCR amplicon libraries were purified using the UltraClean-htp 96 Well PCR Cleanup Kit (Qiagen). Amplicon libraries were quantified by fluorometry (Quant-iT dsDNA High Sensitivity Kit; Invitrogen, Waltham, MA) on a SynergyHT plate reader (BioTek) and combined in equimolar ratios into one pool. The pooled library was concentrated by ethanol precipitation and gel purified (QIAquick Gel Extraction Kit, Qiagen). Each pool of V4-V5 16S rRNA amplicons was sequenced (2×300nt paired end) on one lane of a MiSeq V2 sequencer (Illumina, San Diego, CA) at the Carver Biotechnology Center of the University of Illinois, producing an average of 42673 reads per sample, with a total of 24,579,392 reads produced for this study. Raw reads were demultiplexed using QIIME 1 (version 1.9.1),^43^ trimmed of non-biological sequence using cutadapt (version 1.14),^44^ and resolved into amplicon sequence variants (ASVs) using DADA2 (version 1.1).^45^ Taxonomy was assigned using a Silva reference database (version 132)^46^ and DADA2’s implementation of the RDP naive Bayesian classifier.^47^ A phylogeny was built using a Silva backbone tree (version 132)^46^ and QIIME 2’s fragment insertion plugin,^48^ which runs SEPP.^49^

### Hydrogen Breath Test Processing

At-home hydrogen breath tests were conducted before and after each of the three study phases for four tests in total for each subject (**Figure 1**). Thirteen breath samples were collected for each HBT using the at-home QuinTron EasySampler Breath Collection Kit. Breath samples were processed within fourteen days of sample collection using the QuinTron BreathTracker analyzer and AlveoVac Extraction system at the Stanford Digestive Health Center in Redwood City to measure the concentrations of hydrogen, methane and carbon dioxide in each sample. Each test was first analyzed using the clinical definition of lactose intolerance which is an increase greater than 20ppm combined concentration of hydrogen and methane gas in any breath sample above the concentration in the baseline sample. The area under the curve for each test was calculated by first subtracting each sample’s combined concentration of hydrogen and methane gas from the combined concentration of hydrogen and methane gas in the baseline sample, then plotting this adjusted concentration for each of the thirteen breath samples for each test versus the minutes elapsed from the first sample and taking the integral of the line connecting the datapoints. For some tests, the baseline sample had a concentration above that of breath samples later in the study, producing negative adjusted values. A negative concentration is not biologically interpretable so for tests that had a later breath sample greater than the baseline sample concentration, we set all concentrations that would have been negative to zero. The raw area under the curve results including the negative values are shown **Supplementary Figure 2**. Two subjects in the first baseline HBT completed the full HBT procedure but did not properly collect breath samples such that the concentrations of hydrogen and methane in the breath sample were not able to be determined. Thus, no HBT data for these two tests were included in any plots or analyses.

### Principal coordinate analysis

Throughout our analysis, both binary Jaccard and Bray-Curtis were used because Bray-Curtis considers both structure and abundance, whereas binary Jaccard only takes into account the presence or absence of a given taxon. This allowed us to both detect changes in abundance of taxa overall as well as the appearance or disappearance of taxa, particularly low-frequency species. Both measures demonstrated similar results so only plots using binary Jaccard were included in the main text and the corresponding Bray-Curtis plots were included in the Supplementary Figures.

### Linear Discriminant Analysis

We performed a supervised linear discriminant analysis using TreeDA.^23^ This package performs sparse discriminant analysis using a phylogenetic tree structure provided to the algorithm. Incorporating the phylogenetic tree structure into the sparse discriminant analysis expands the feature space to include higher-order taxonomic units as predictors alongside the specific ASVs. For example, it is possible that at the ASV level a signal cannot be detected, due to high strain-level variation between participants, but a consistent pattern can be seen across participants when considering a higher taxonomic level such as the family Bifidobacteriaceae. The necessary inputs are the classes to be discriminated (our elimination vs. reintroduction phase samples), a set of predictors (the taxa abundances identified in our data), and a tree describing the relationship between the predictors (the phylogenetic tree of the microbial taxa). We compared samples from the last three weeks of the elimination phase to those in the last three weeks of the reintroduction phase so that the intervention had time to take effect, especially during the reintroduction which was a more gradual intervention. This resulted in a confusion matrix that correctly labeled 91 of the 105 elimination samples as elimination, and 85 of the 106 reintroduction samples as reintroduction. We also compared samples from the entire elimination phase versus those from the entire reintroduction phase, as well as those from just the last two weeks of the elimination and reintroduction phase. Of note, the resulting top taxa prioritized by this analysis were highly sensitive to parameter input, such as transformation and filtering of sample counts, and were not all supported by some of the further investigations.

We evaluated the performance of TreeDA on a negative control of discriminating between samples across the study and subjects based on whether the sample was processed on an even or odd plate, which should have no correlation with microbiota structure. This negative control test was to make sure the test would fail if there were truly no signal. This resulted in a confusion matrix that incorrectly classified the plate as even for 150 of the 172 samples on an odd numbered plate. It classified the plate as even for 210 of 232 samples on even numbered plates indicating that the model was, as expected, unable to discriminate between these arbitrary classifications relative to microbiota structure and was instead simply classifying almost all samples as even.

We also repeated the test to discriminate between samples in the last three weeks of the elimination phase versus the last three weeks of reintroduction phase on each subject individually to see even if we intentionally overpowered the test if it could find any signal, but this did not result in any interpretable consistent patterns.

### Autocorrelation time

We defined autocorrelation time as the typical number of days that it takes until two consecutive samples from the same subject have at least the same distance between them as 90% of the median distance between any two randomly chosen samples where again both samples are from the same subject. To calculate this, we first found the median distance across all sample pairs where both pairs were from the same subject individually for each subject regardless of the days between, giving us the median distance between any two randomly picked samples for a given subject. Then, we binned each sample pair where both samples were from the same subject into half week bins (based on the days between when the samples in the pair were collected) and calculated the median distance for each half week bin for each subject. We then found, for each subject, the minimum half week bin whose median distance between samples reached at least 90% of the distance between the median distance between any two randomly picked samples for that subject. We took the median of this number across subjects giving us an autocorrelation time of 5.25 days using the binary Jaccard measure. In addition, we calculated the autocorrelation time using the Bray-Curtis distance metric instead of binary Jaccard, which resulted in an autocorrelation time of 7.00 days.

The choice of defining autocorrelation as a fraction of the median distance between any two randomly chosen samples was to account for the expected asymptotic nature of the distance between microbiota samples over time. The autocorrelation time defined using the fraction of 90% was reported but because this fraction choice was slightly arbitrary, we also calculated the autocorrelation time based on definitions using 80% and 95% respectively. The former gave an autocorrelation time of 3.50 days, and the latter gave an autocorrelation time of 10.5 days.

The median value across subjects of 90% of the median distance between any two randomly chosen samples (for each subject) was 0.394 for binary Jaccard and 0.310 for Bray-Curtis distance metrics. This is higher than the average median distance between our technical controls (five pairs of two swabs from the same stool) of 0.267 for binary Jaccard and 0.152 for Bray-Curtis.

## Supporting information

blank_participant_log.xlsx

ReportedSymptoms_FinalWeek.xlsx

HBTsymptoms.xlsx

## Acknowledgements

We thank all the study subjects, without whom this research and paper would not have been possible. We thank Arati Patankar for guidance on the experimental methods and Anna Robaczewska for the barcoded primers. We thank Ruth Hicks and Rafael Ruiz De Luzuriaga and other personnel of the Redwood City Medical Clinic for training and permission for use of the equipment for processing the HBT samples. We thank the members of the Relman and Feldman labs, Avihu Yona, and Yoav Ram, for helpful comments and discussions on the study design and analysis approach. This study was supported by the Stanford TRAM (Translational Research and Applied Medicine) pilot grant program. C.J.S. is supported by a National Science Foundation Graduate Research Fellowship and Stanford’s Knight-Hennessy Scholars Program. O.K. is supported by the US–Israel Binational Science Foundation (BSF), the Israel Science Foundation (ISF), and the Gordon and Betty Moore Foundation. D.A.R. is supported by the Thomas C. and Joan M. Merigan Endowment at Stanford University.

