## Supplementary Information and Figures for "Short-Term Dairy Elimination and Reintroduction Minimally Perturbs the Gut Microbiota in Self-Reported Lactose Intolerant Adults"

| **Demographic variable** | **Summary Statistics** |
| --- | --- |
| Age | Median: 32 years old; Range: 23-60 years |
| Sex | Female: n=9; Male: n=3 |
| Prior amount of dairy products in diet | No dairy: n=2; some dairy: n=4; lactose-free dairy: n=4; regular dairy: n=2 |
| Clinical status from HBT from first baseline | Intolerant: n=8; tolerant: n=4 |

**Supplementary Table 1.** Aggregate background information on subjects that participated in the study.

[Attached document titled “blank_participant_log.xlsx”]

**Supplementary Table 2.** Example empty log given to subjects to track symptoms and notable diet or lifestyle changes.

[Attached documents titled “HBTsymptoms.xlsx” and “ReportedSymptoms_FinalWeek.xlsx”]

**Supplementary Table 3.** Summary of subjects’ self-reported symptoms during the (a) HBT and (b) final week. 0 corresponds to no symptoms, 1 corresponds to very mild symptoms such as gassiness, 2 corresponds to experiencing mostly mild symptoms but occasionally moderate symptoms such as cramping or other abdominal pain, 3 corresponds to moderate symptoms, and 4 corresponds to severe symptoms such as diarrhea.

**Supplementary Note 1.** One lesson learned from this pilot study is that because the gut microbiota structure is incredibly resistant to this perturbation, a longer or more aggressive intervention is likely needed to see a more obvious shift in microbiota structure for a study of this size. The aggressiveness of the dairy reintroduction intervention was limited by feasibility and ethical considerations because the subjects were self-identified lactose intolerant human subjects and would understandably be deterred by too aggressive of a dairy reintroduction. Another lesson is that a longer baseline period allowing for additional baseline measurements of both microbiota structure and lactose intolerance symptoms is needed to better identify what constitutes normal baseline fluctuations in the absence of an intervention. This is especially important in the case of the HBT where the variation between the two baseline samples were often the same, if not more, than the variation from the post-elimination and post-reintroduction samples. A possible explanation for this is that the HBT necessitated a major dietary restriction in the preceding 24-hour period to each test and also required subjects to drink two cups of milk (see **Methods**). Therefore, because many of the subjects had not consumed significant amounts of dairy products in a long time, the first Hydrogen Breath test at the beginning of the study might have been an unintentionally significant dietary intervention. A future way to identify if this could have played a role in shifting the microbiota at the beginning of the study would be to start fecal swab sample collection prior to the first HBT.

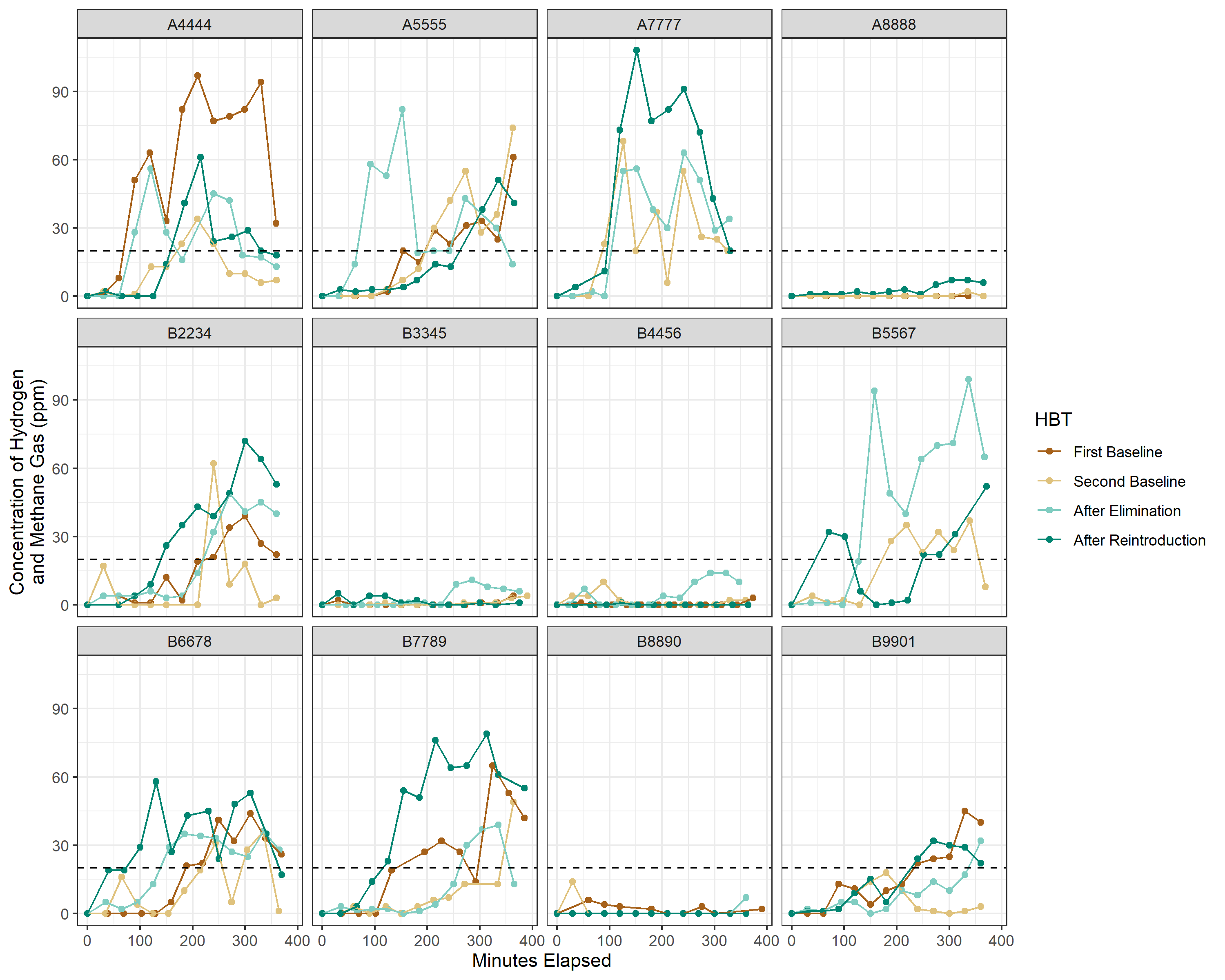
**Supplementary Figure 1.** HBT results for each subject showing the combined concentration of hydrogen and methane gas in their breath (with the baseline sample value subtracted from each, or set to 0 if adjusted value was negative) as time elapsed after drinking two cups of milk. Individuals are classified by the HBT as lactose intolerant in the clinic if the sum of hydrogen and methane concentrations in their breath sample reaches greater than 20 ppm above their first sample concentration at any timepoint during the HBT. At baseline, the HBT classified 8 of the 12 subjects as lactose intolerant and none of the subjects changed their initial status of lactose tolerance during the study tests. HBT baseline results for subjects A7777 and B5567 are not included because the breath sample collection was not completed properly.

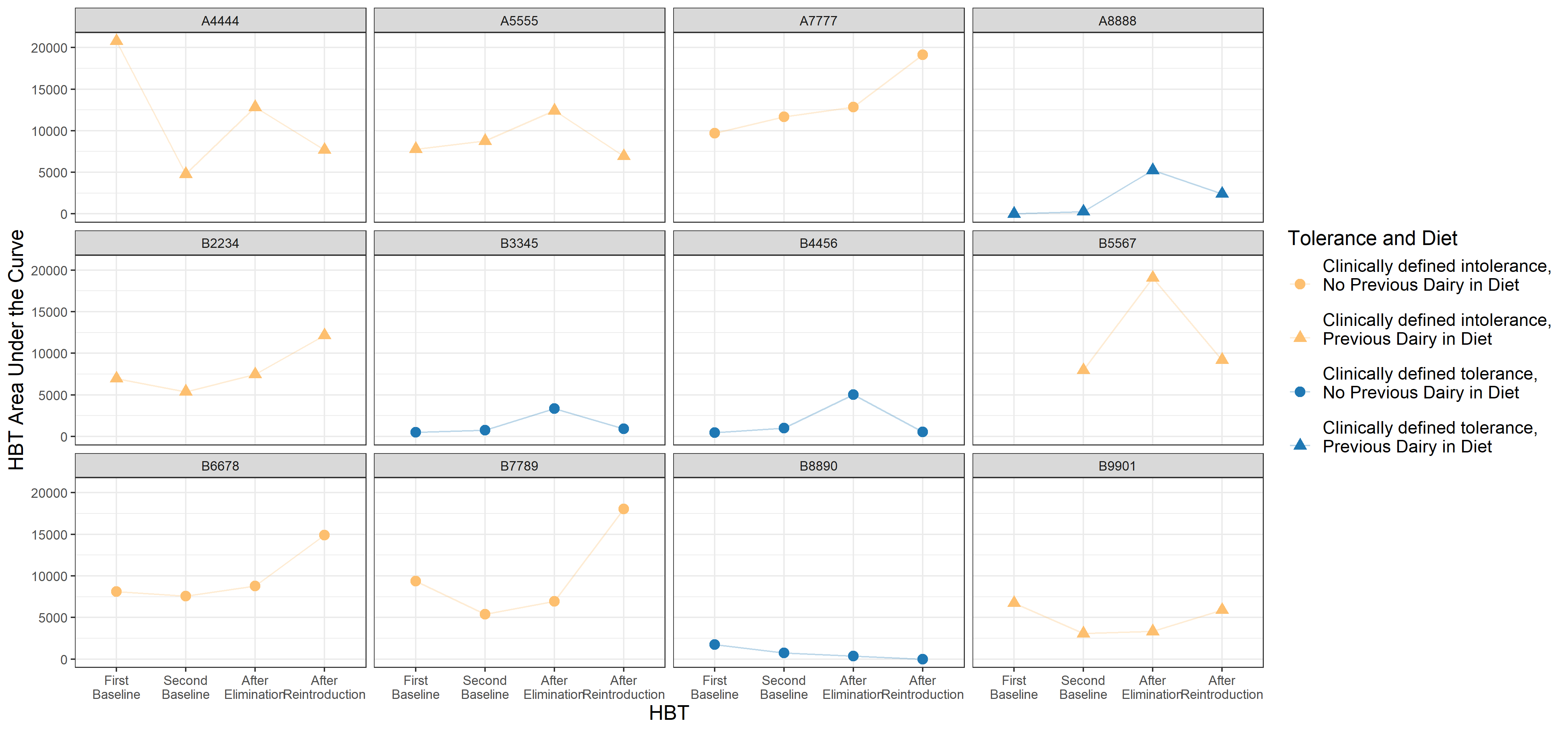

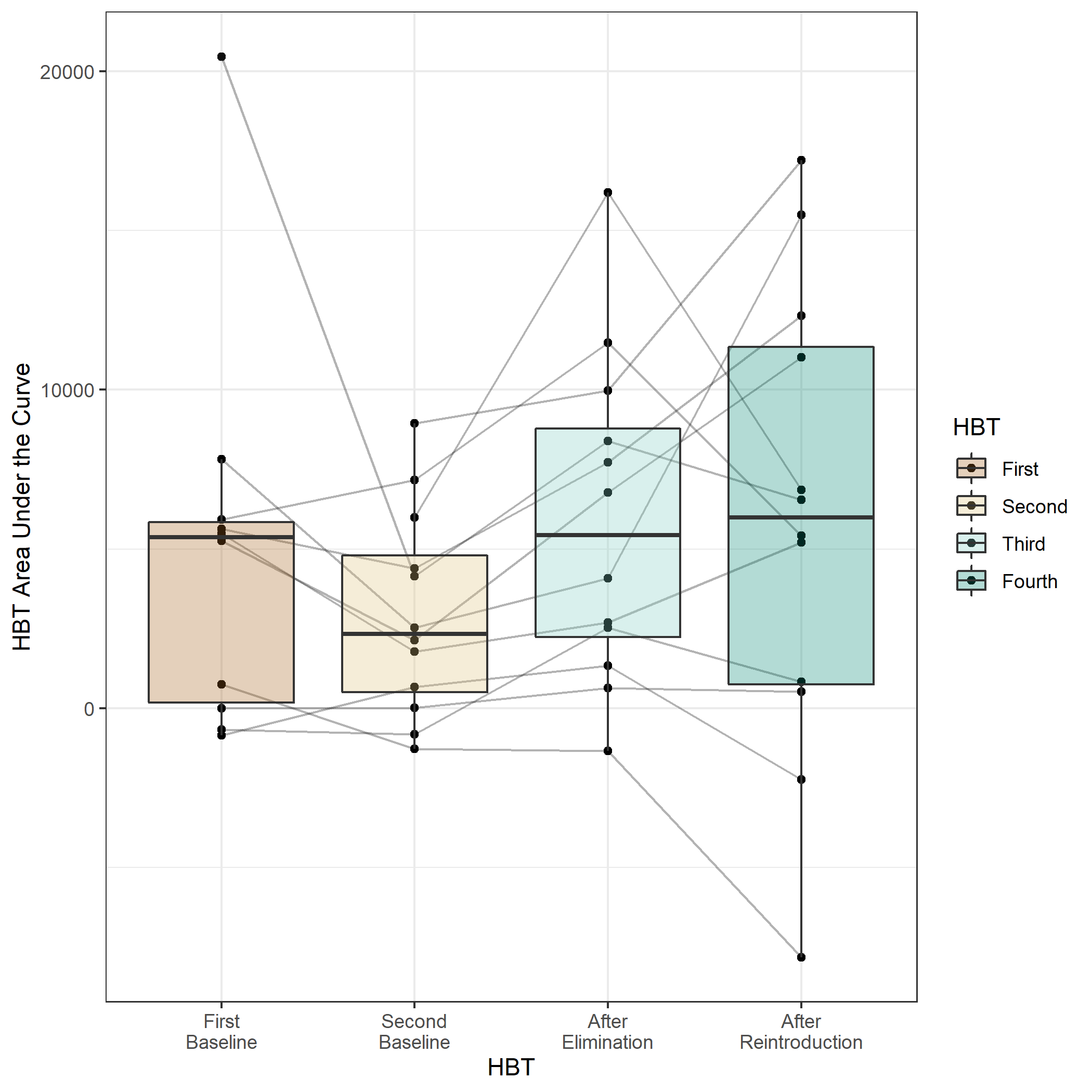

(a)

(b)

**Supplementary Figure 2.** (a) Individual results for area under the curve of the HBT for each subject across the study. (b) Aggregate area under the curve of the HBT for each subject across the study, showing raw values instead of those shown in **Figure 2** where any negative calculated breath sample (after subtracting the concentration measured in a given breath sample from the baseline sample concentration for that test) was set to 0. See Methods for further information. As shown in **Figure 2**, subjects’ HBT area under the curve significantly increased after the dairy elimination phase, potentially reflecting an increase in intolerance.

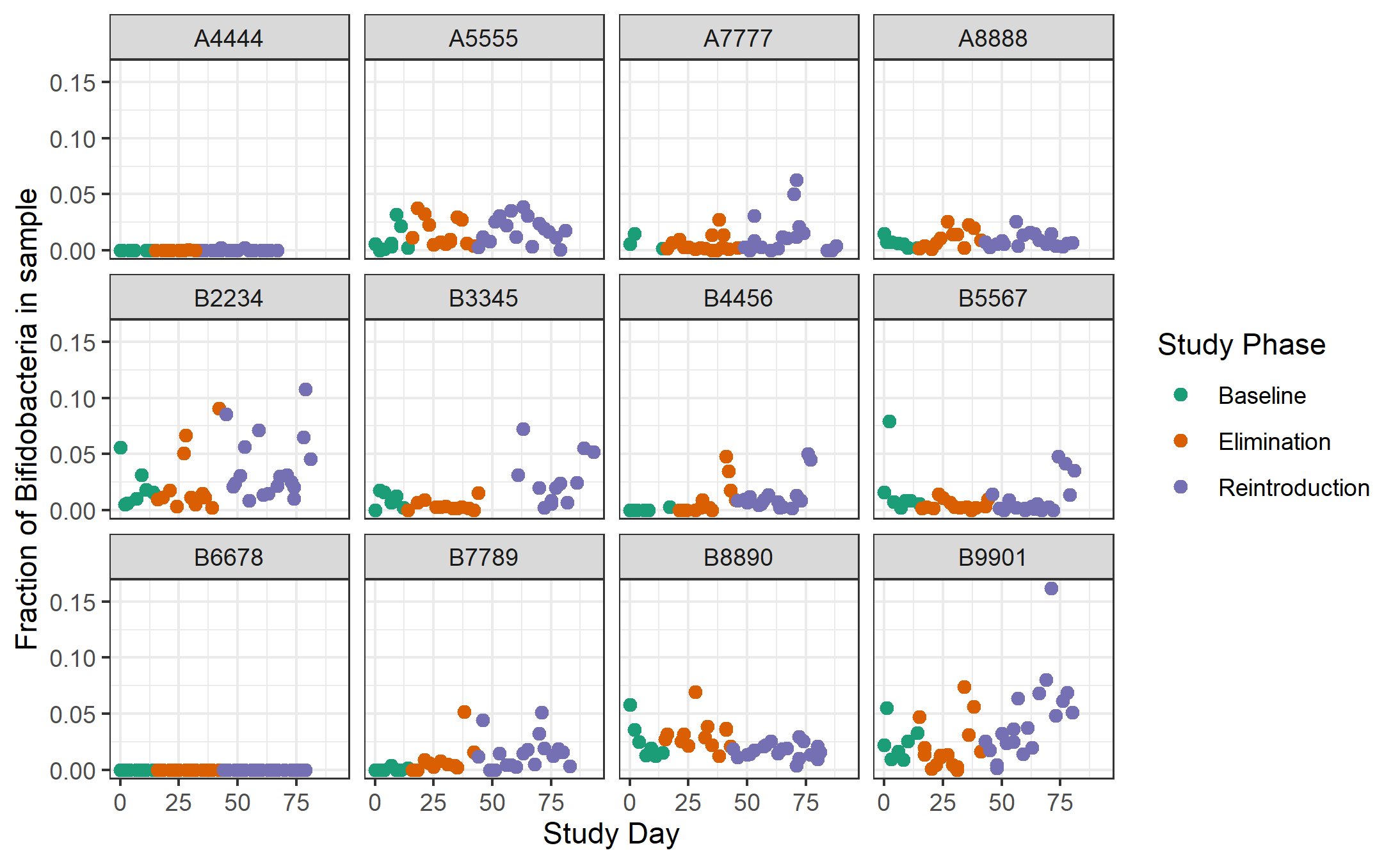
**Supplementary Figure 3.** Abundance of *Bifidobacterium* in each sample (normalized based on the total bacterial abundance in that sample) for all subjects over the course of the study, colored by study phase. *Bifidobacterium* was one of the taxa prioritized by the supervised linear discriminant analysis with TreeDA and was also one of our hypothesized candidate taxa. There was not an obvious decrease in the abundance of *Bifidobacterium* in most subjects during the elimination phase, or an obvious increase during the reintroduction phase as might be expected.

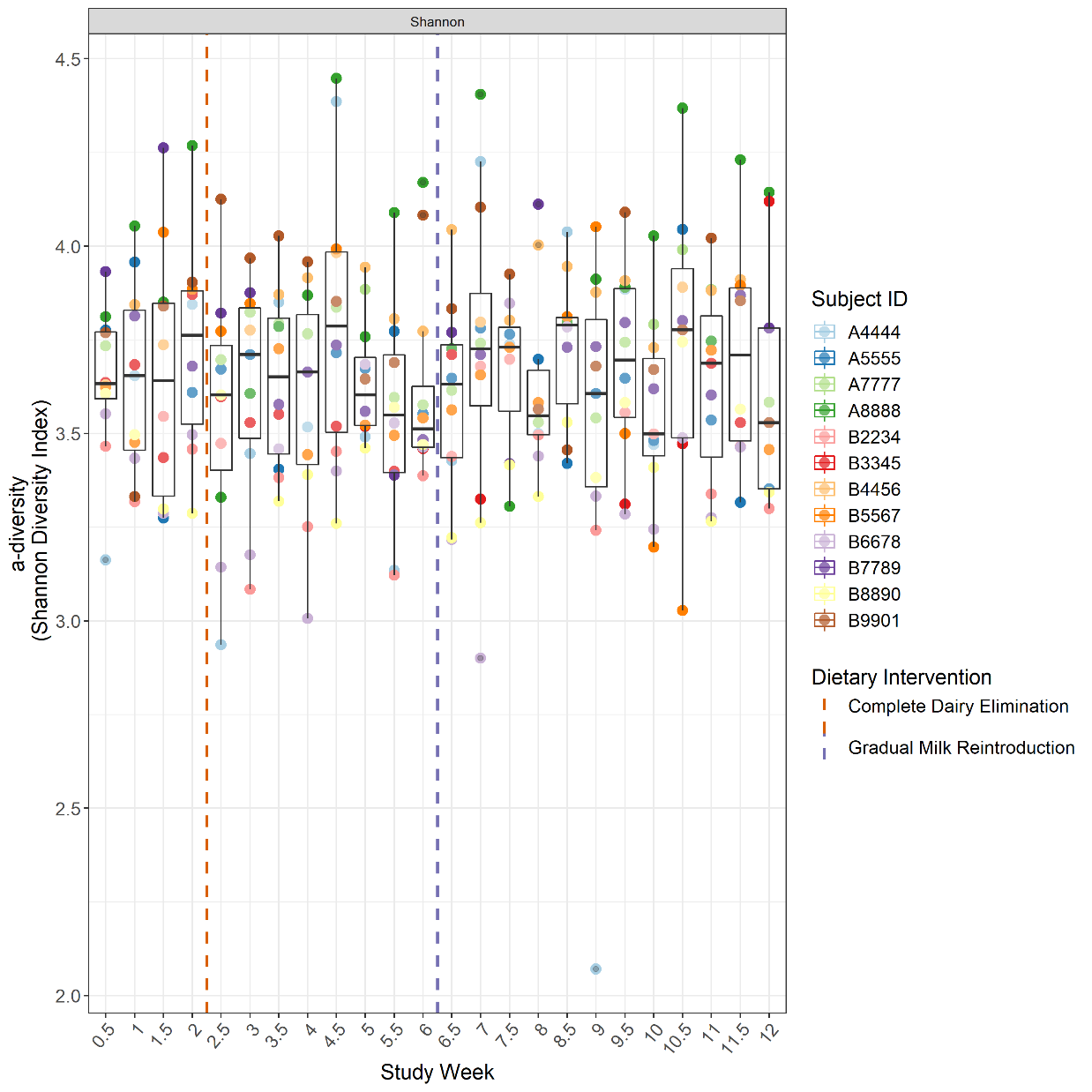
**Supplementary Figure 4.** Alpha-diversity (Shannon Diversity index) in half week temporal bins with a maximum of one sample from each subject included in each bin. If more than one sample for a given subject was collected within a given half-week bin, one sample was randomly selected to be the one included. Not every bin has every subject due to occasional incidental variations in sampling frequency by some subjects. No consistent patterns of alpha-diversity changes throughout the study across subjects were found.

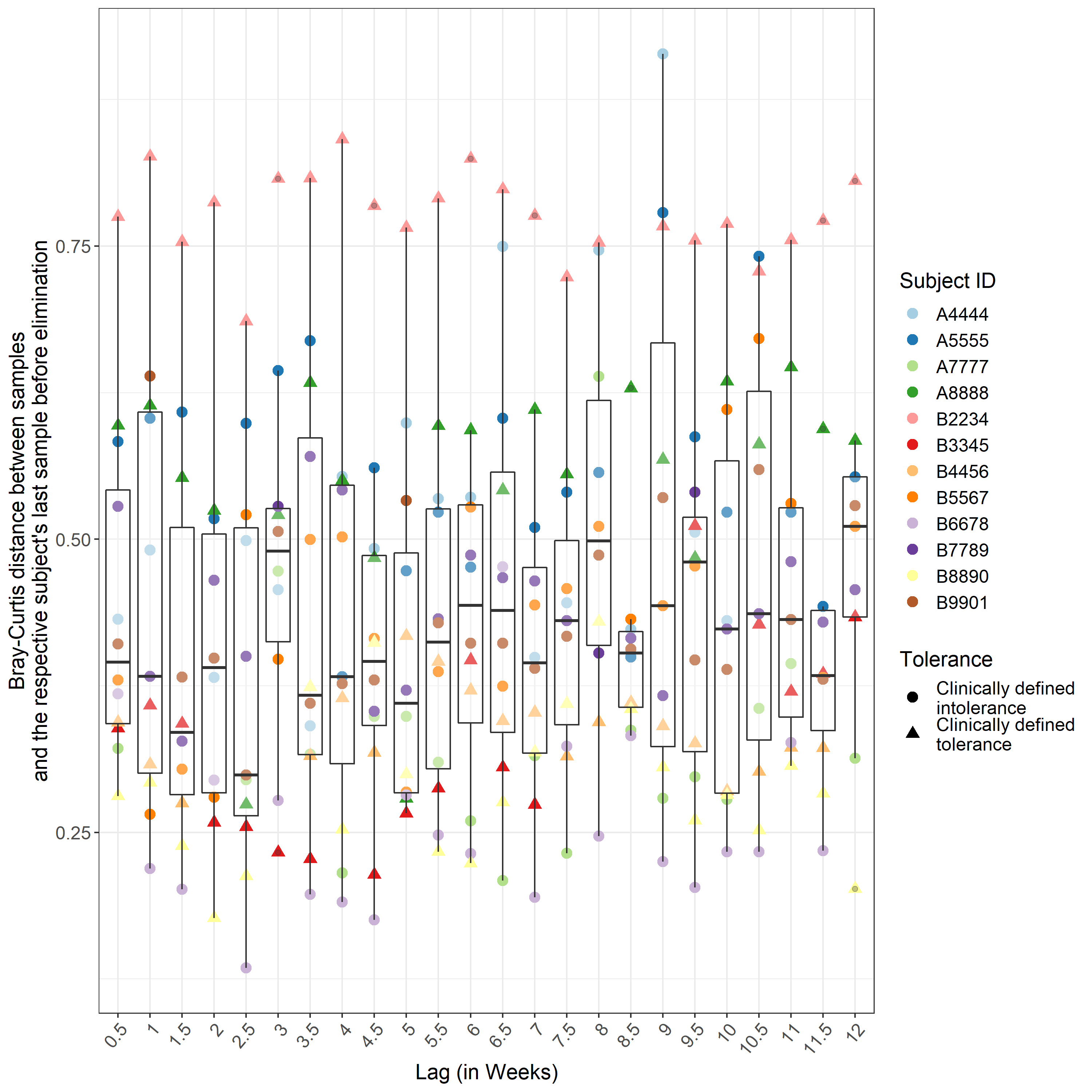

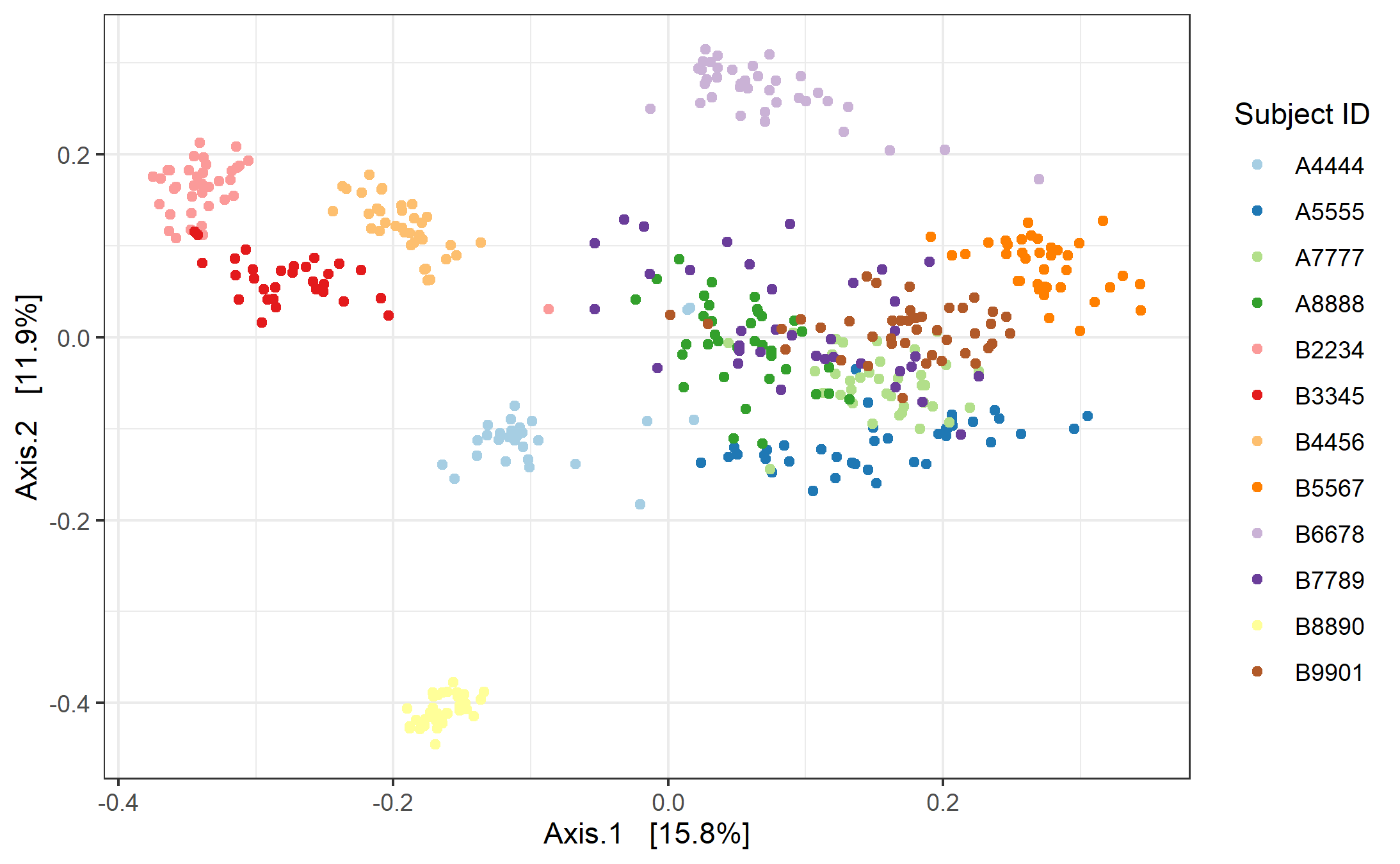

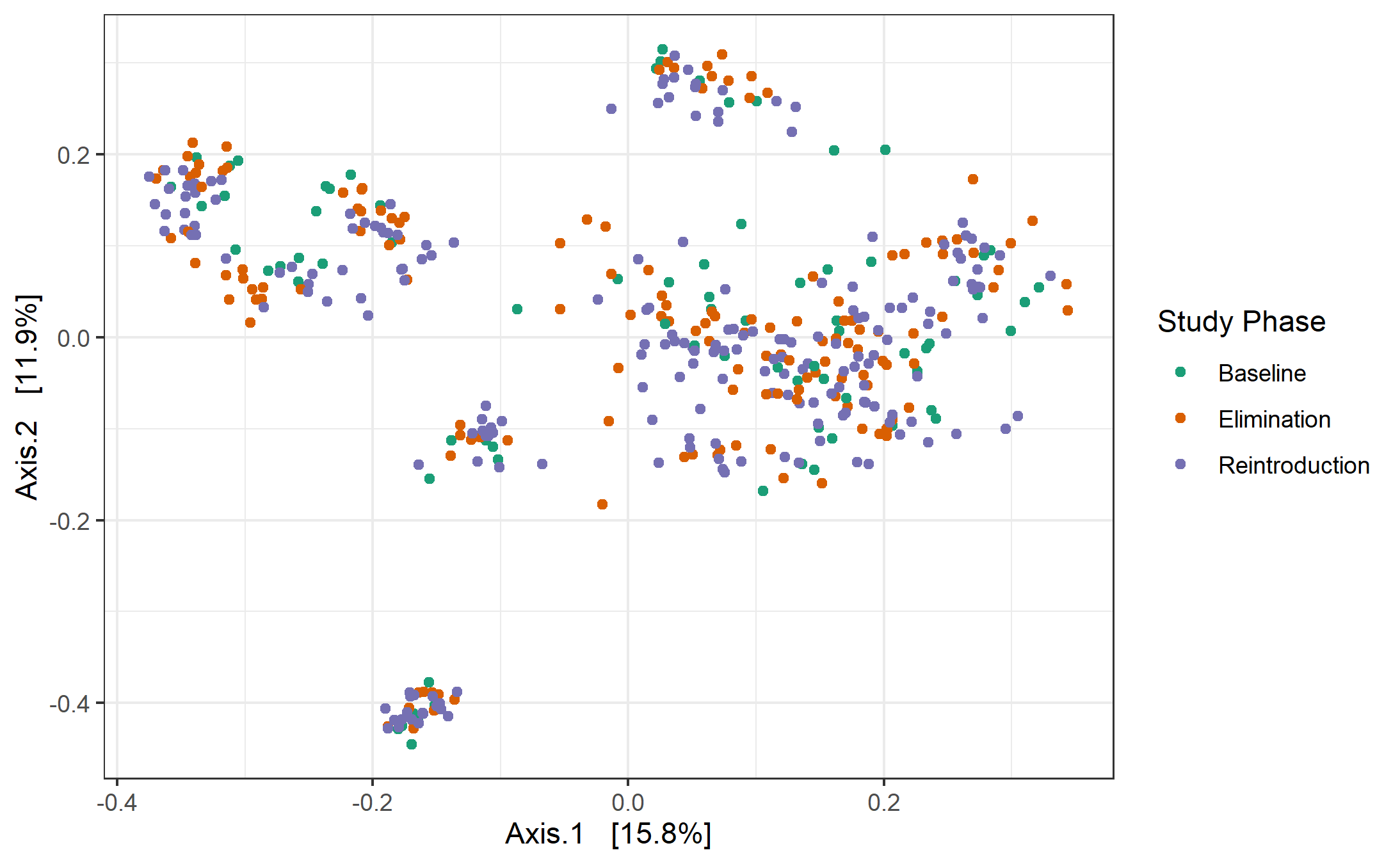
**Supplementary Figure 5.** Visualization of the first two principal coordinates from PCoA across all samples for all subjects using distance metrics of Bray-Curtis index, colored by subject (a) and by study phase (b). Samples throughout the study clustered predominately based on subject ID over study phase Permanova with 1000 permutations on Bray-Curtis distances; by subject ID: R^2^ = 0.728, p-value=0.001; by study phase: R^2^ = 0.004, p-value=0.665. Study phase nested within subject ID has R^2^ = 0.036, p-value=0.001. (c) The beta-diversity (Bray-Curtis distance) between each subject’s last sample before dairy elimination and that subject’s samples other time points. The samples are placed in half-week bins with a maximum of one sample from each subject included in each bin. The half week bins correspond to the number of days between when the samples in each sample pair were collected (lag), not the study day. If more than one sample for a given subject was collected within a given half-week bin, one sample was randomly selected to be the one included. Not every bin has every subject due to occasional incidental variations in sampling frequency by some subjects.

(c)

(a)

(b)

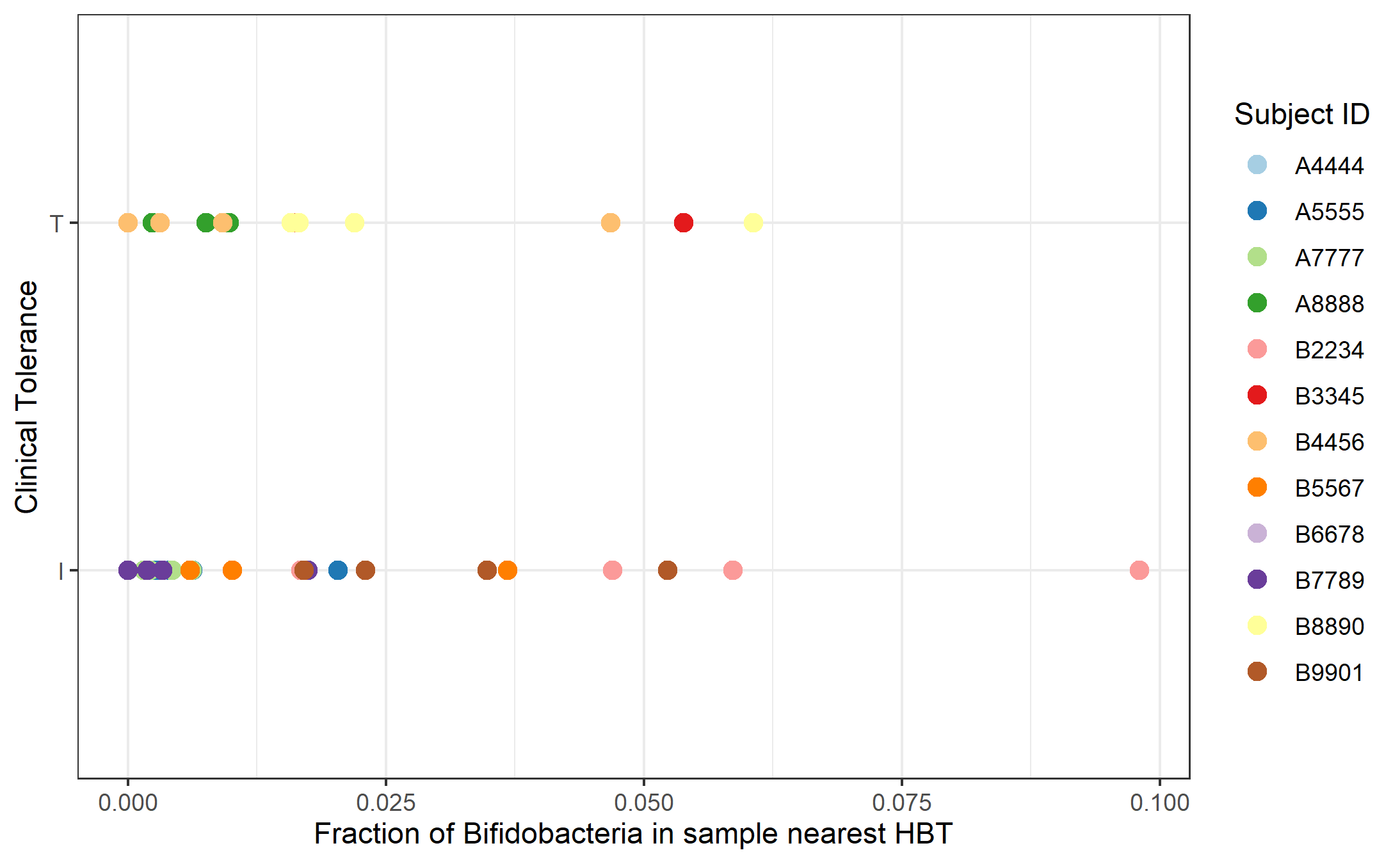

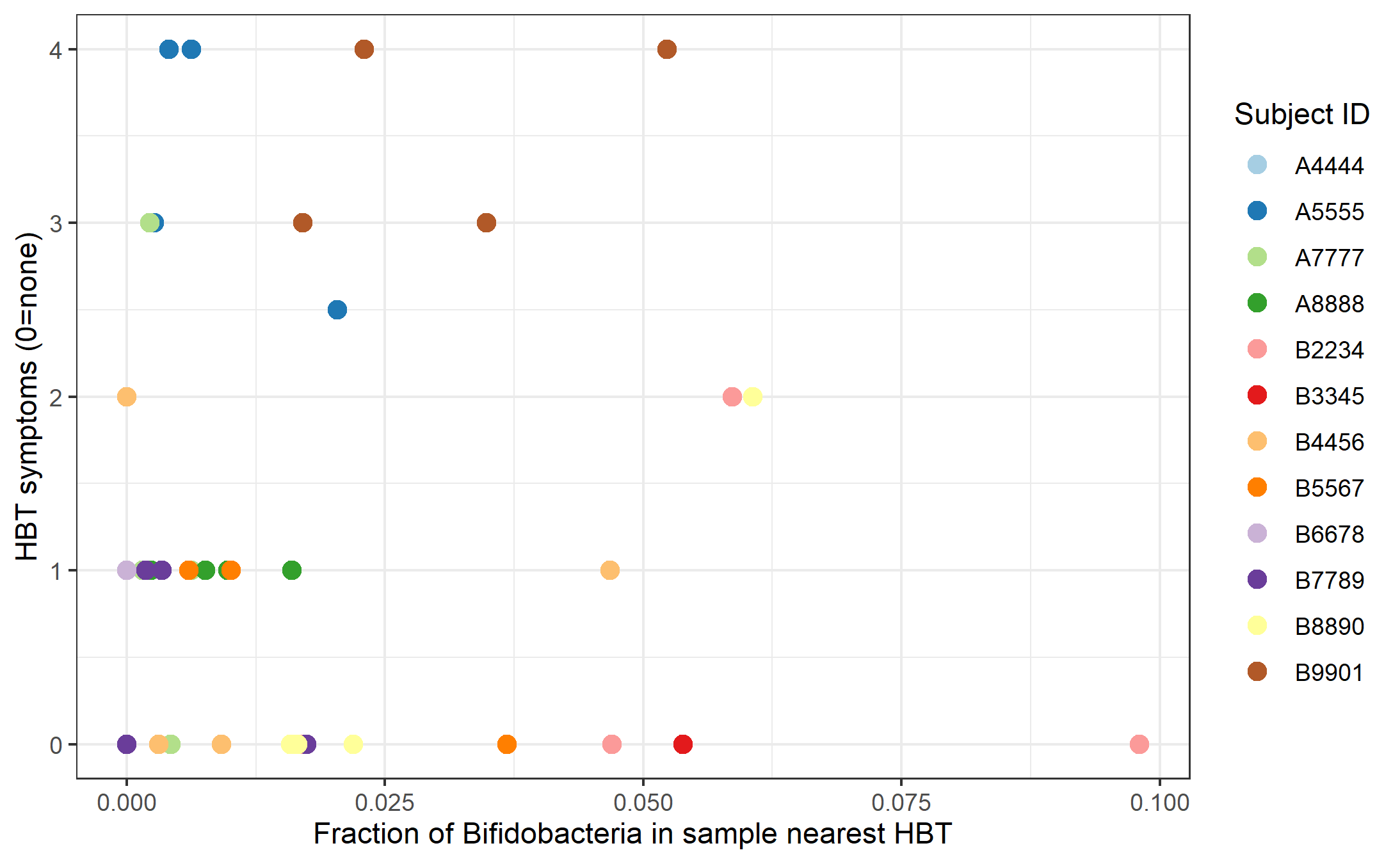

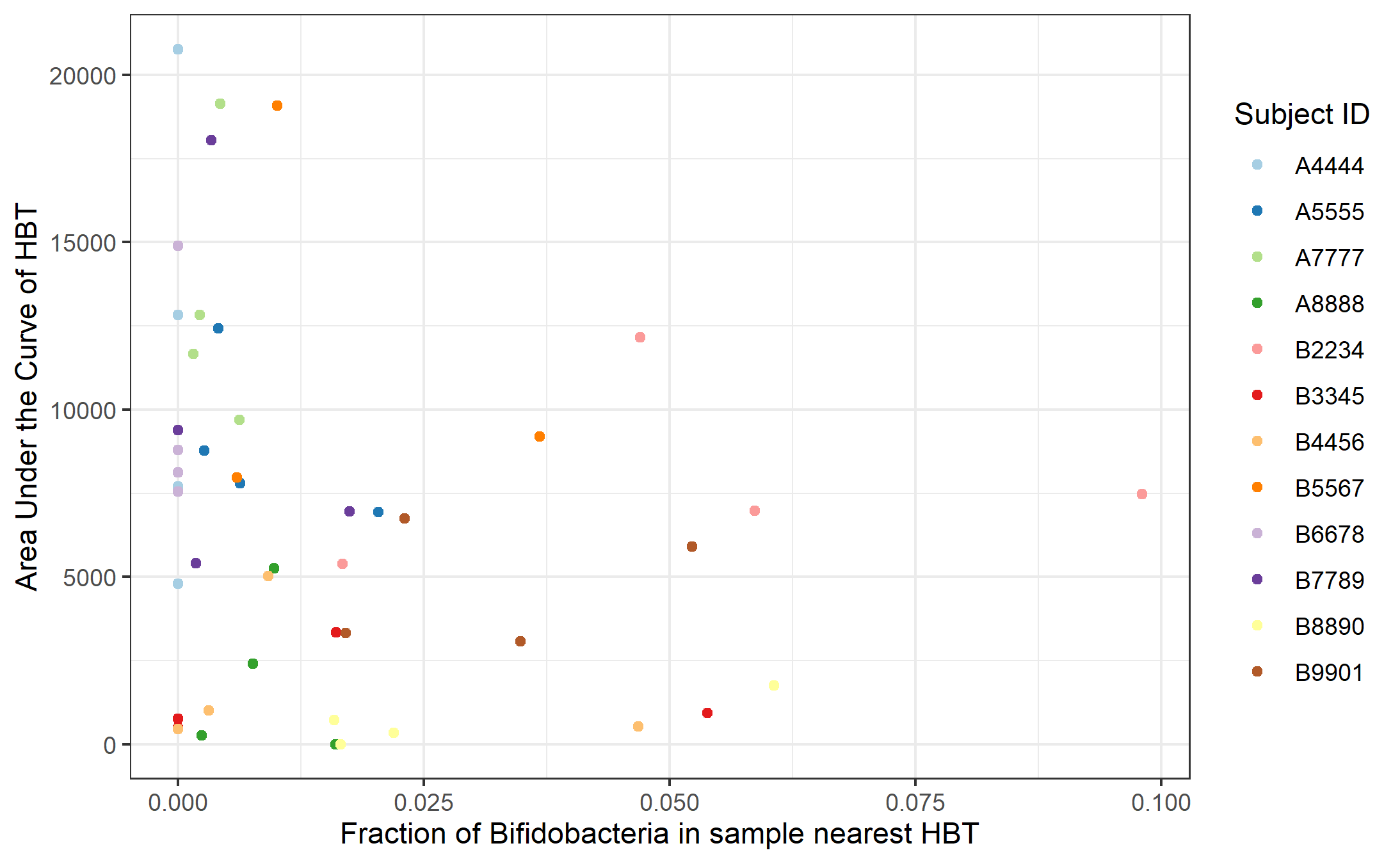

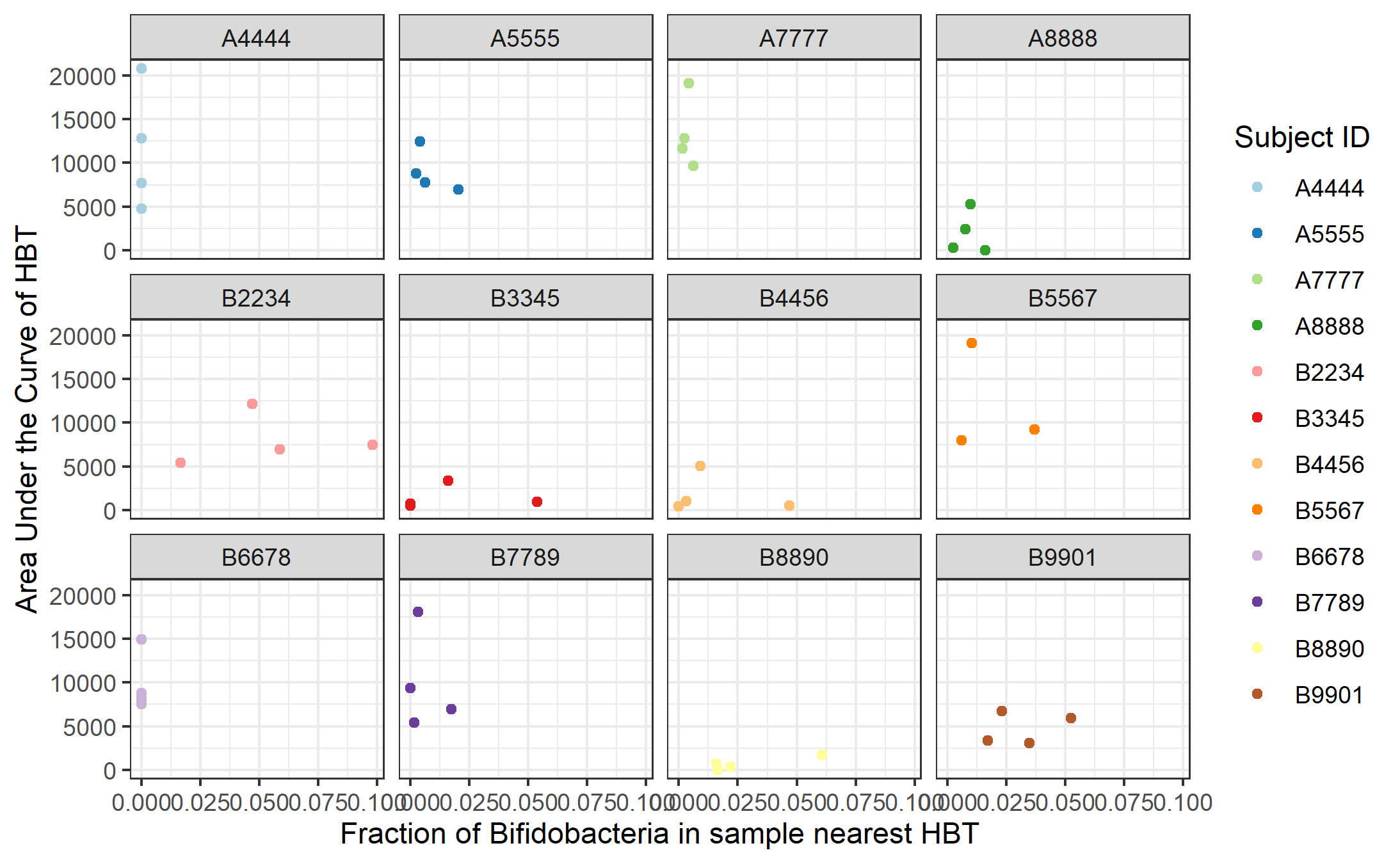
**Supplementary Figure 6.** The HBT area under the curve of combined concentration for all subjects (a) together and (b) individually versus the abundance of *Bifidobacterium* in the sample (normalized based on the total bacterial abundance in that sample) that was collected nearest in time to when the HBT was performed. No significant relationship was found (r(44) = -0.212, p=0.152).

(a)

(b)

(b)

(a)

**Supplementary Figure 7.** (a) Clinical lactose tolerance status as defined by the HBT versus the abundance of *Bifidobacterium* in the sample (normalized based on the total bacterial abundance in that sample). b) Rating of self-reported symptoms by subjects (0=no symptoms, 4=severe symptoms such as diarrhea) during each HBT versus the fraction of *Bifidobacterium* in the sample nearest that HBT. Results did not show a significant relationship between abundance of *Bifidobacterium* (t(34) = -0.362, p=0.719) and Clinical Tolerance or symptoms (r(44) = 0.035, p=0.818).

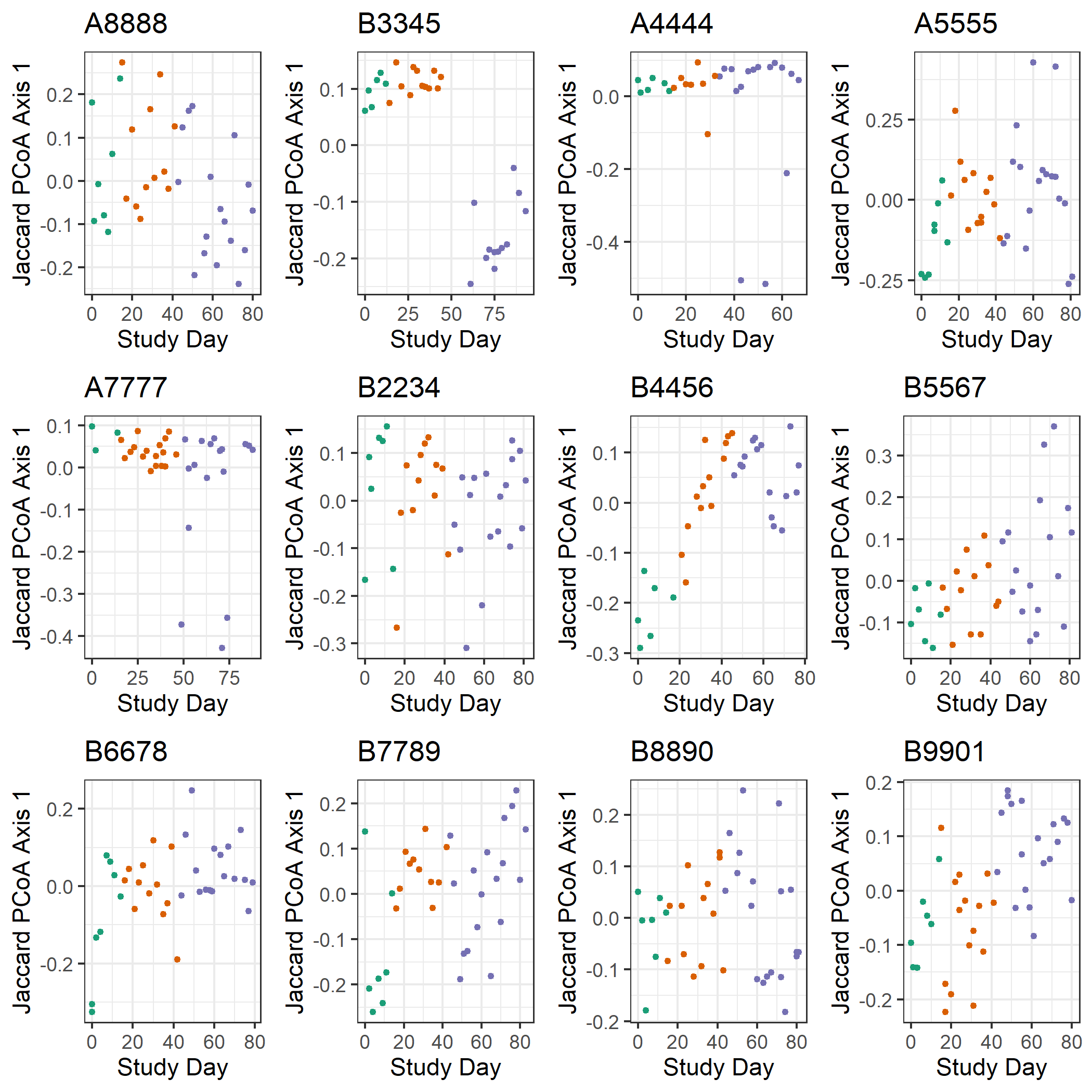

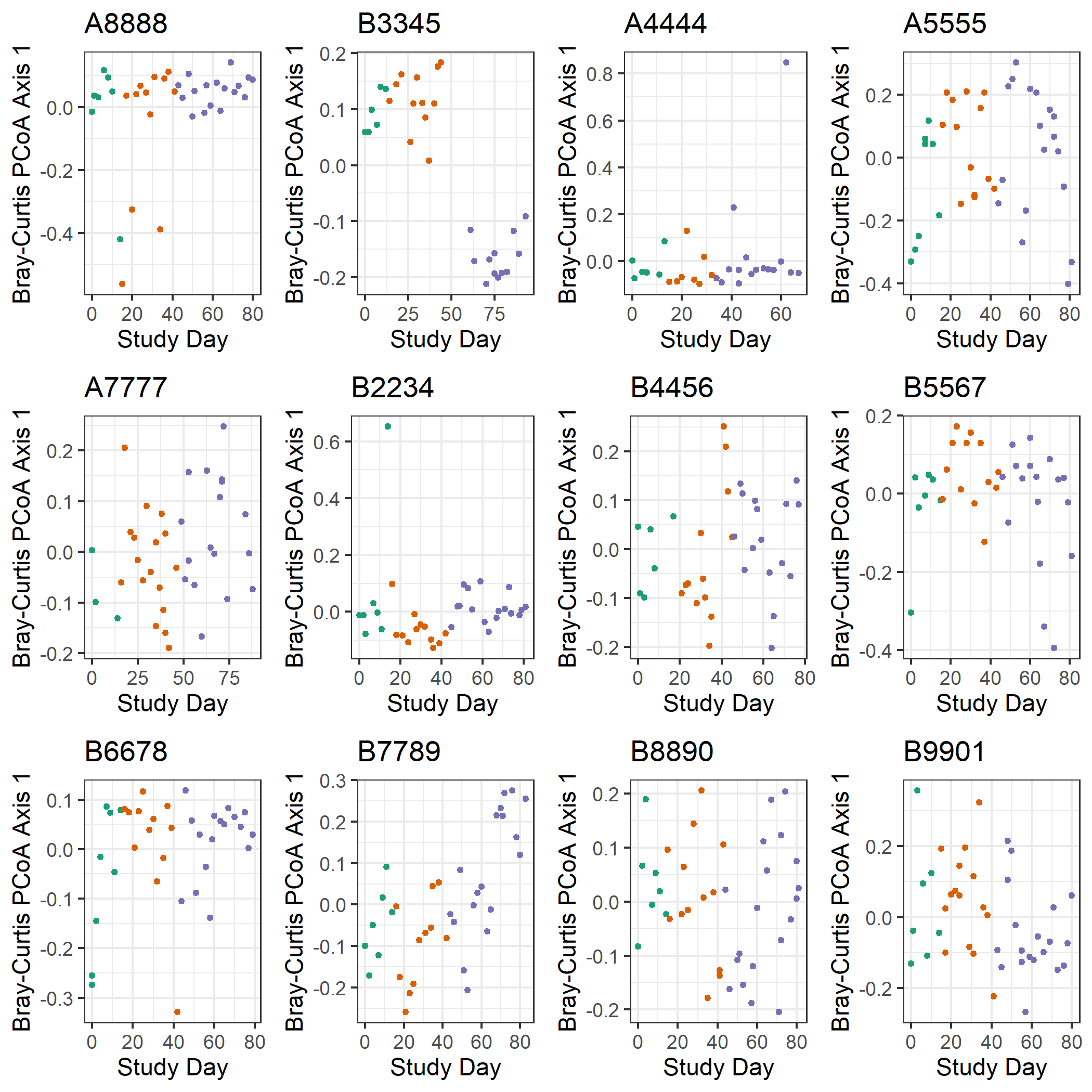

(b)

(a)

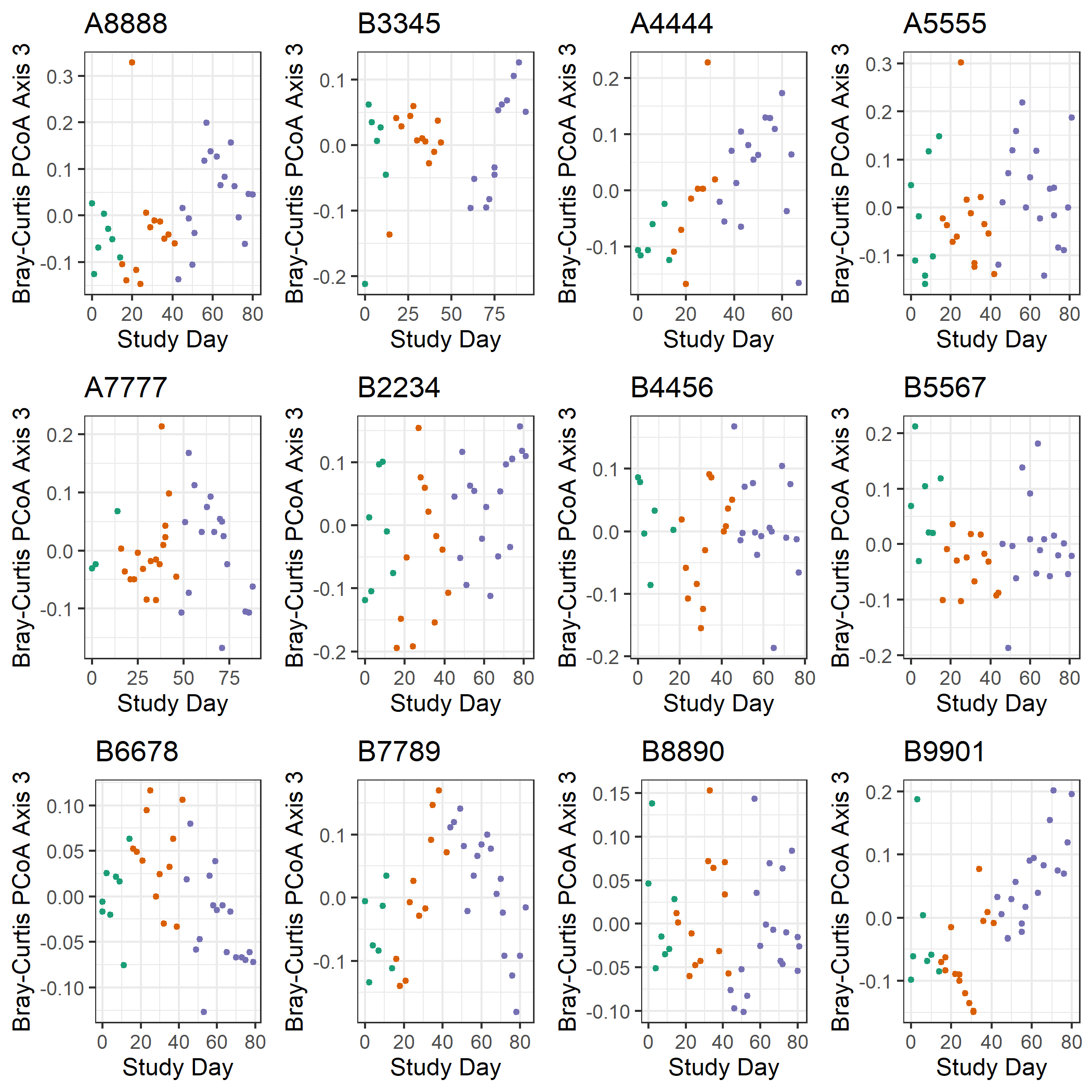

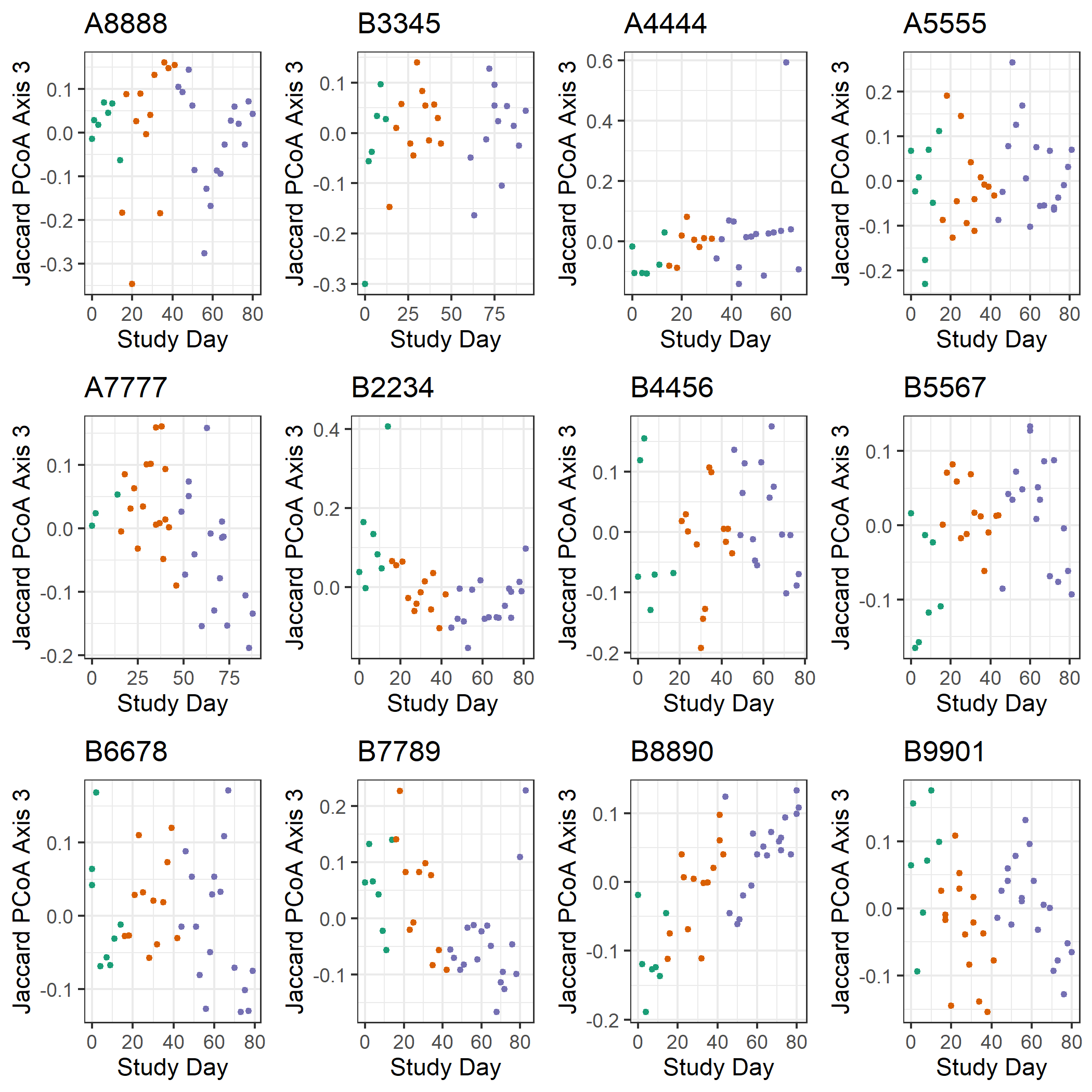

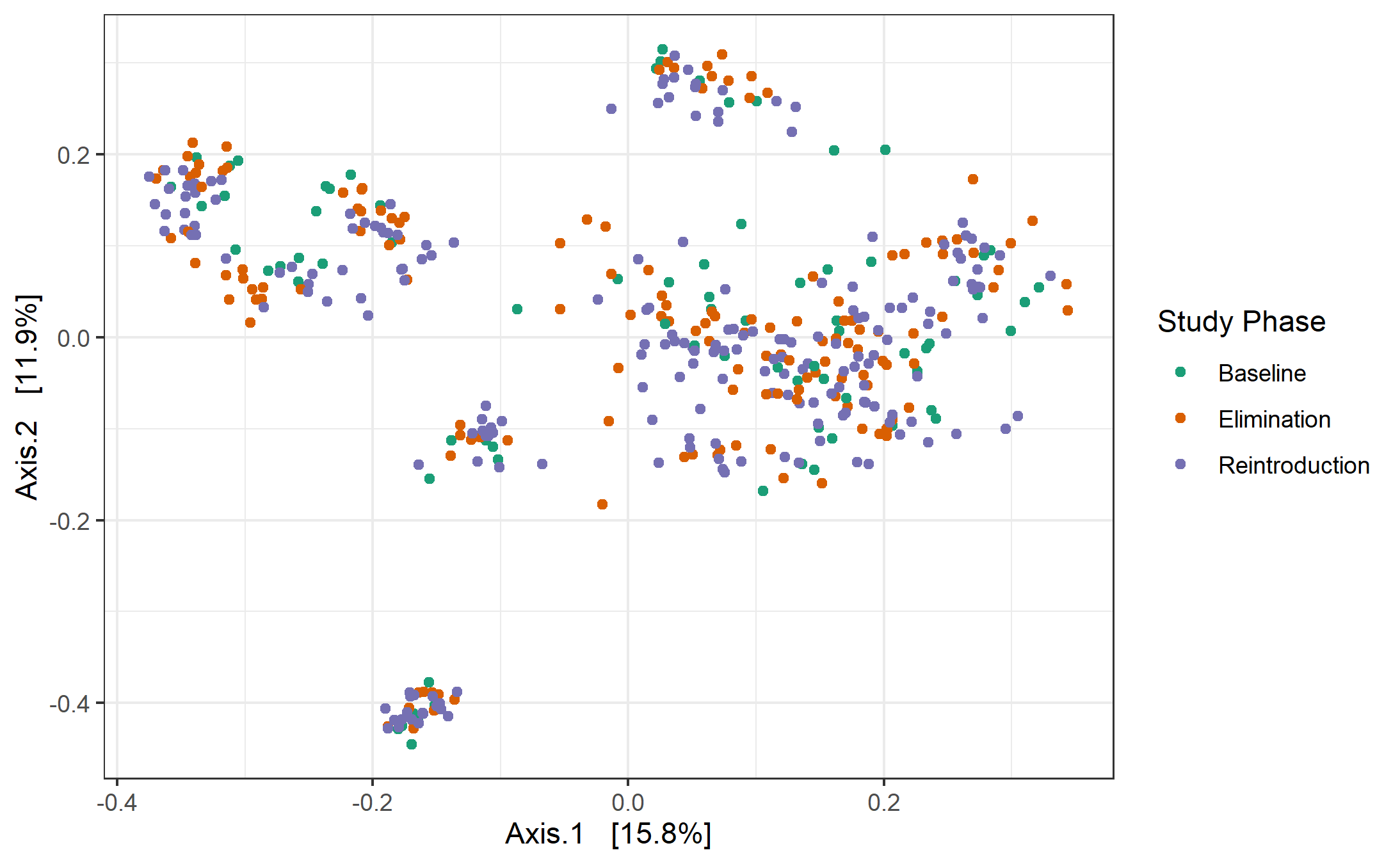

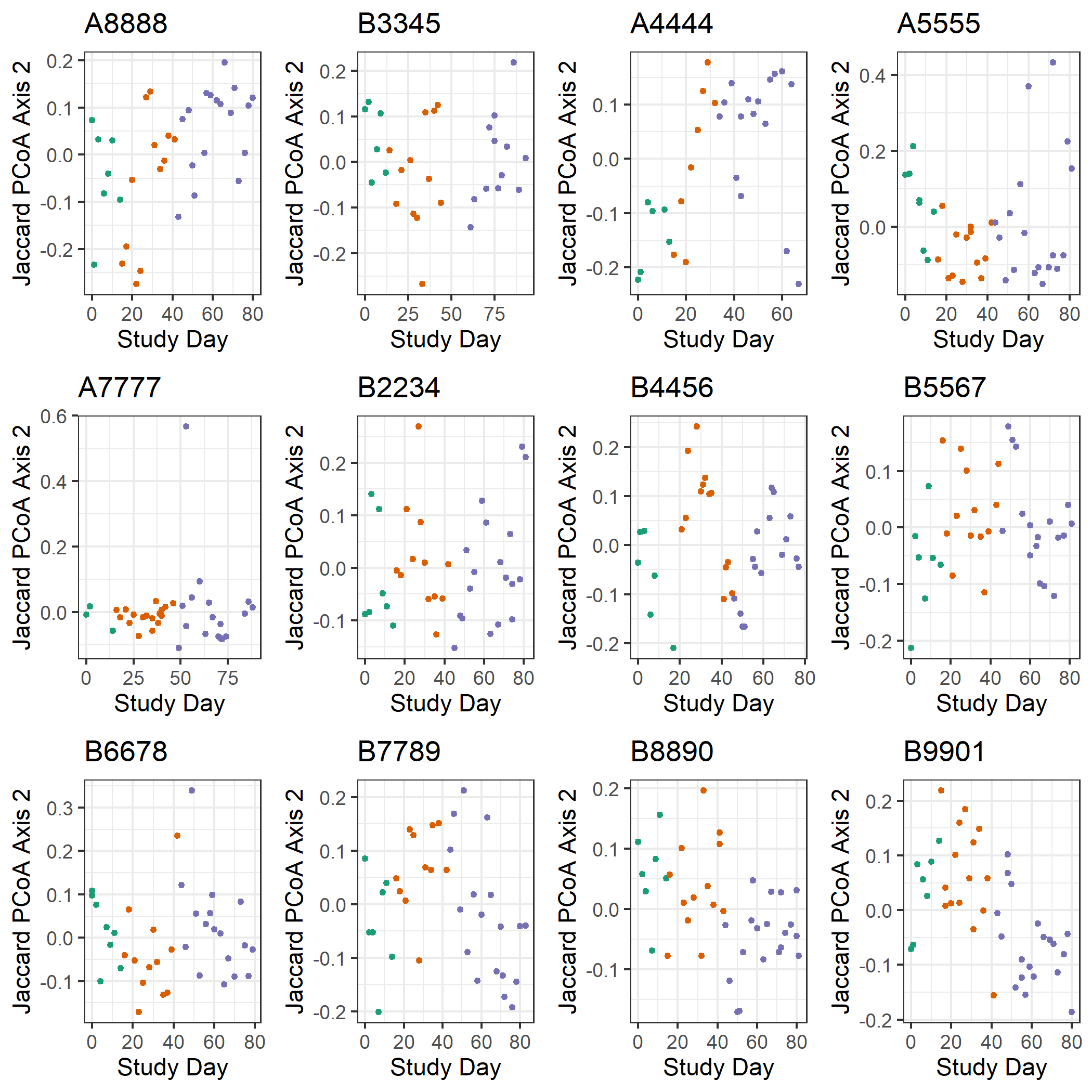

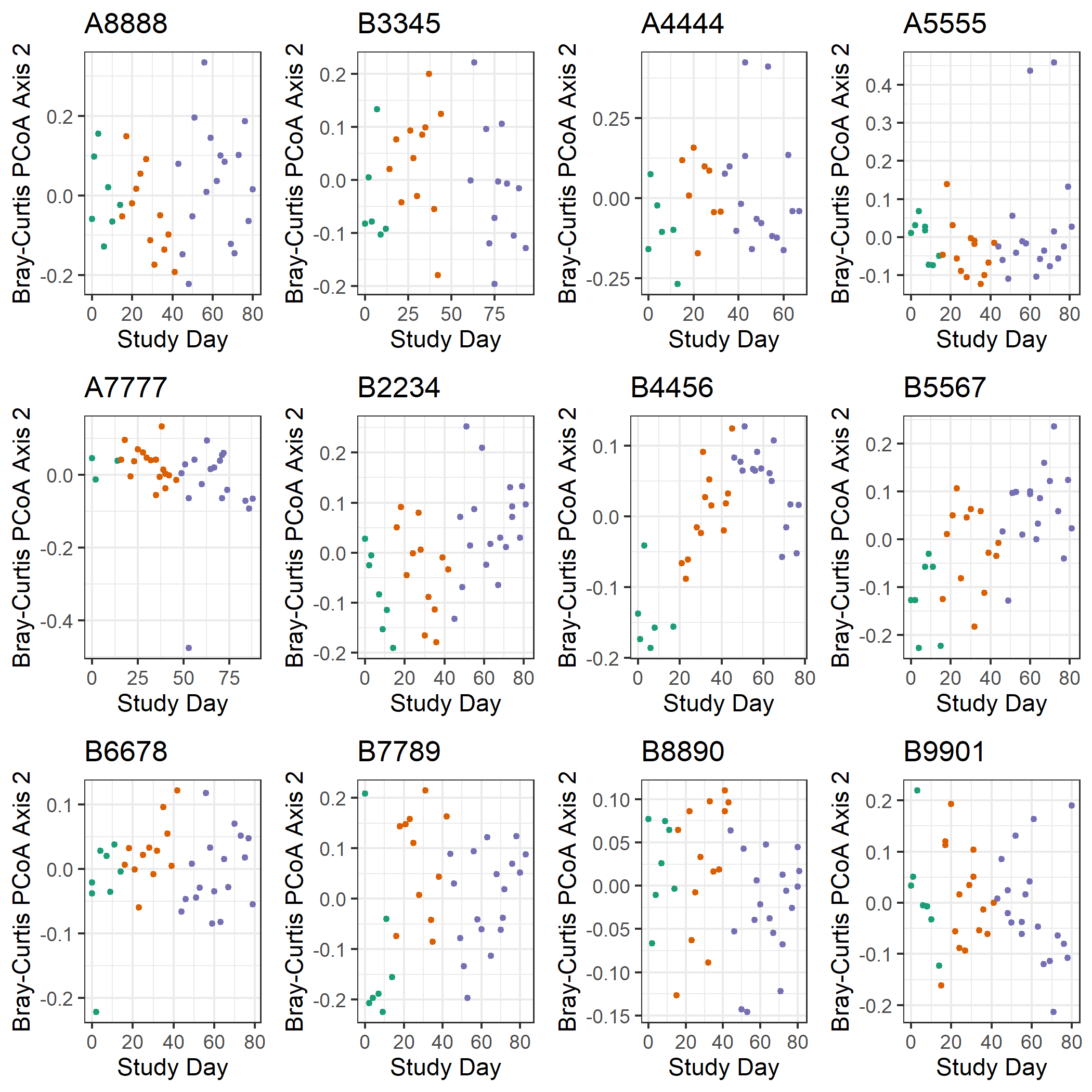

(e)

(f)

(c)

(d)

**Supplementary Figure 8.** Visualization of the first principal coordinates from PCoA across all samples for each subject individually using (a,c,e) Bray-Curtis and (b,d,f) Jaccard index distance metrics versus the day in the study for that subject that the sample was collected, colored by study phase. These results suggest that there is little correlation between the first principal coordinates from the PCoAs in **Figure 2** and the time course of the study. It is surprising that even for non-weighted distance measures such as Jaccard index, there seems to be limited overall microbiota changes which means that few low abundance species, such as the candidate taxa expected to respond to the intervention, dropped in or out of presence in the microbiota. A caveat to this, however, is that the interventions might not have been dramatic or long enough and that if they were, a shift would have been found.

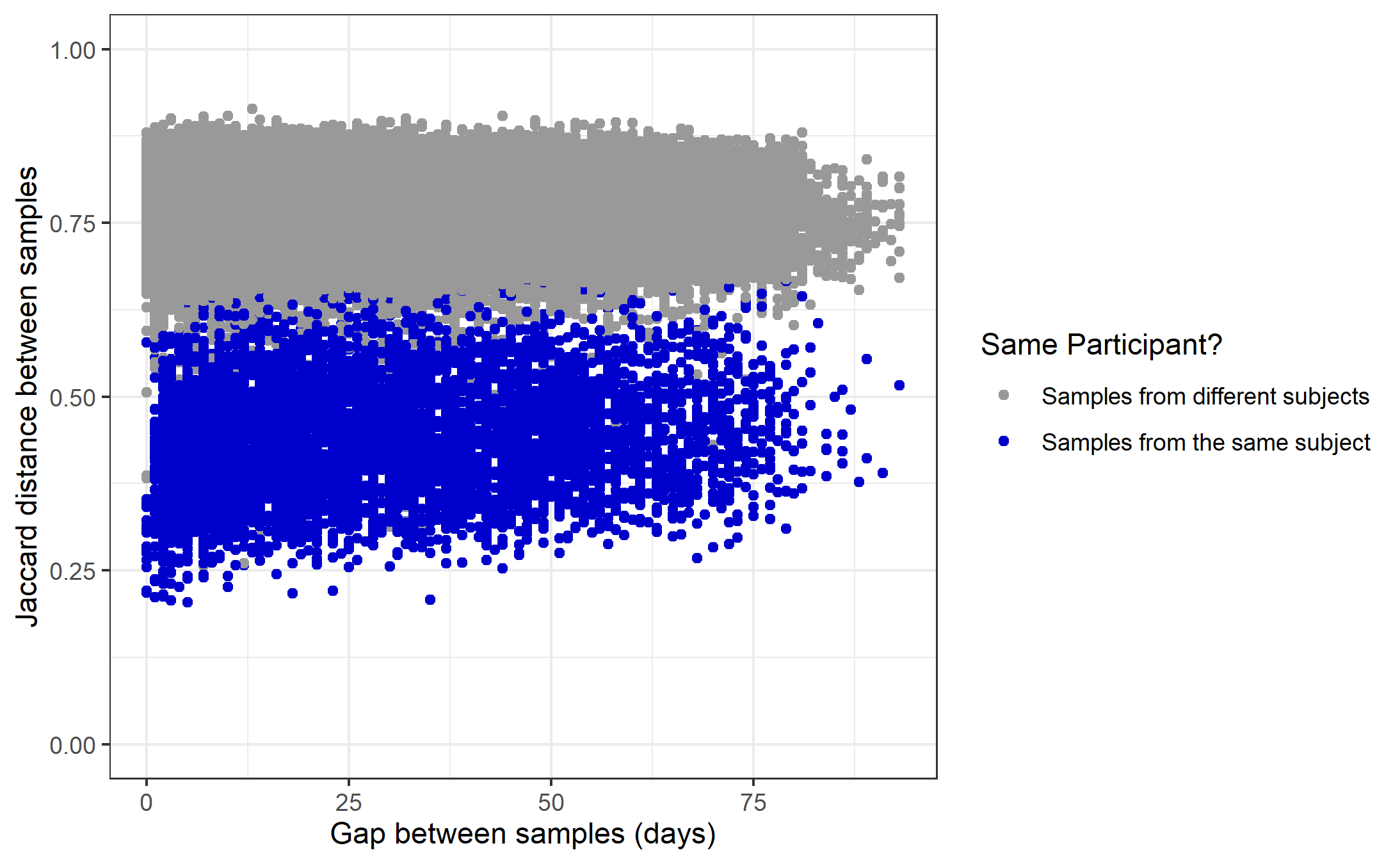

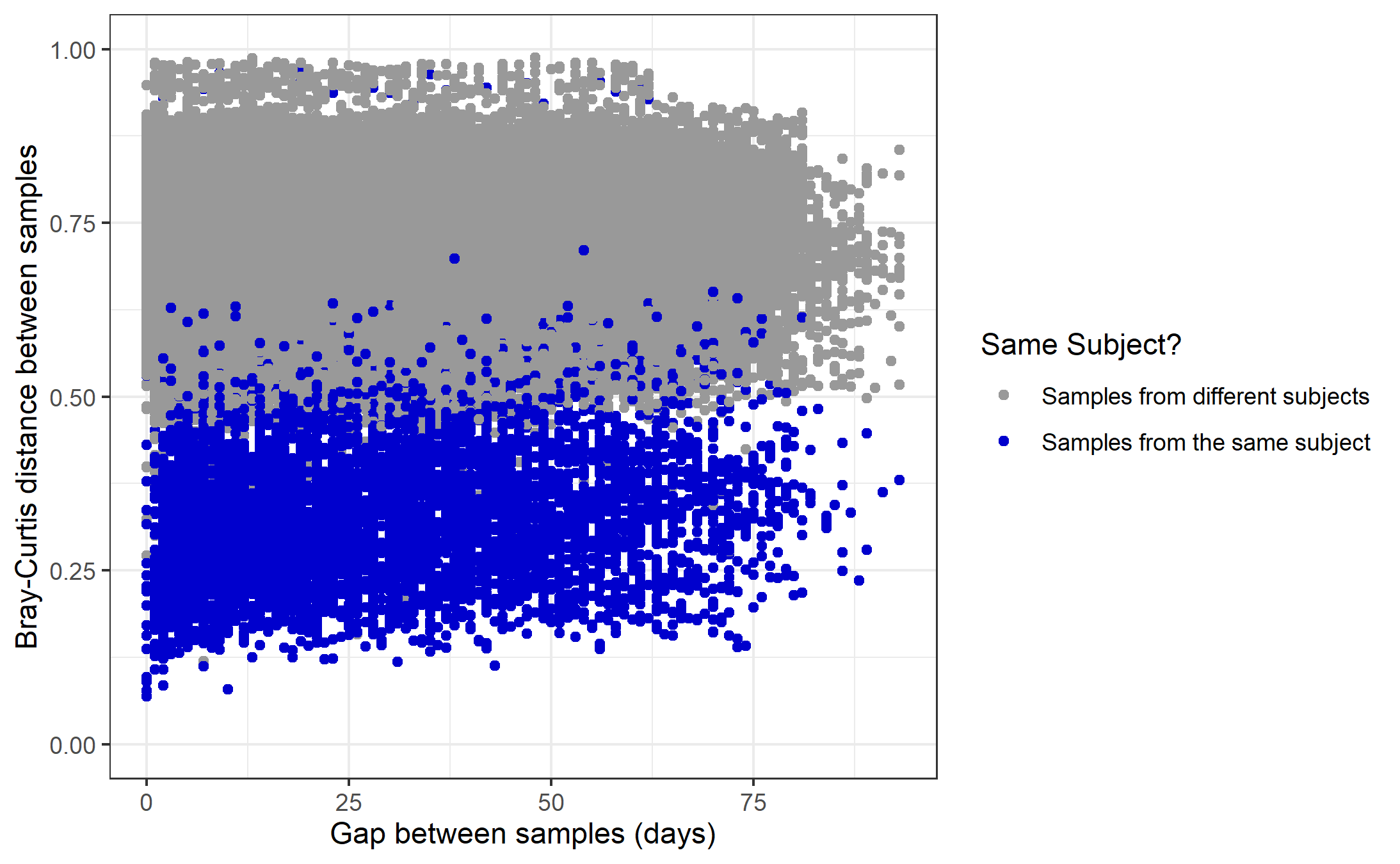

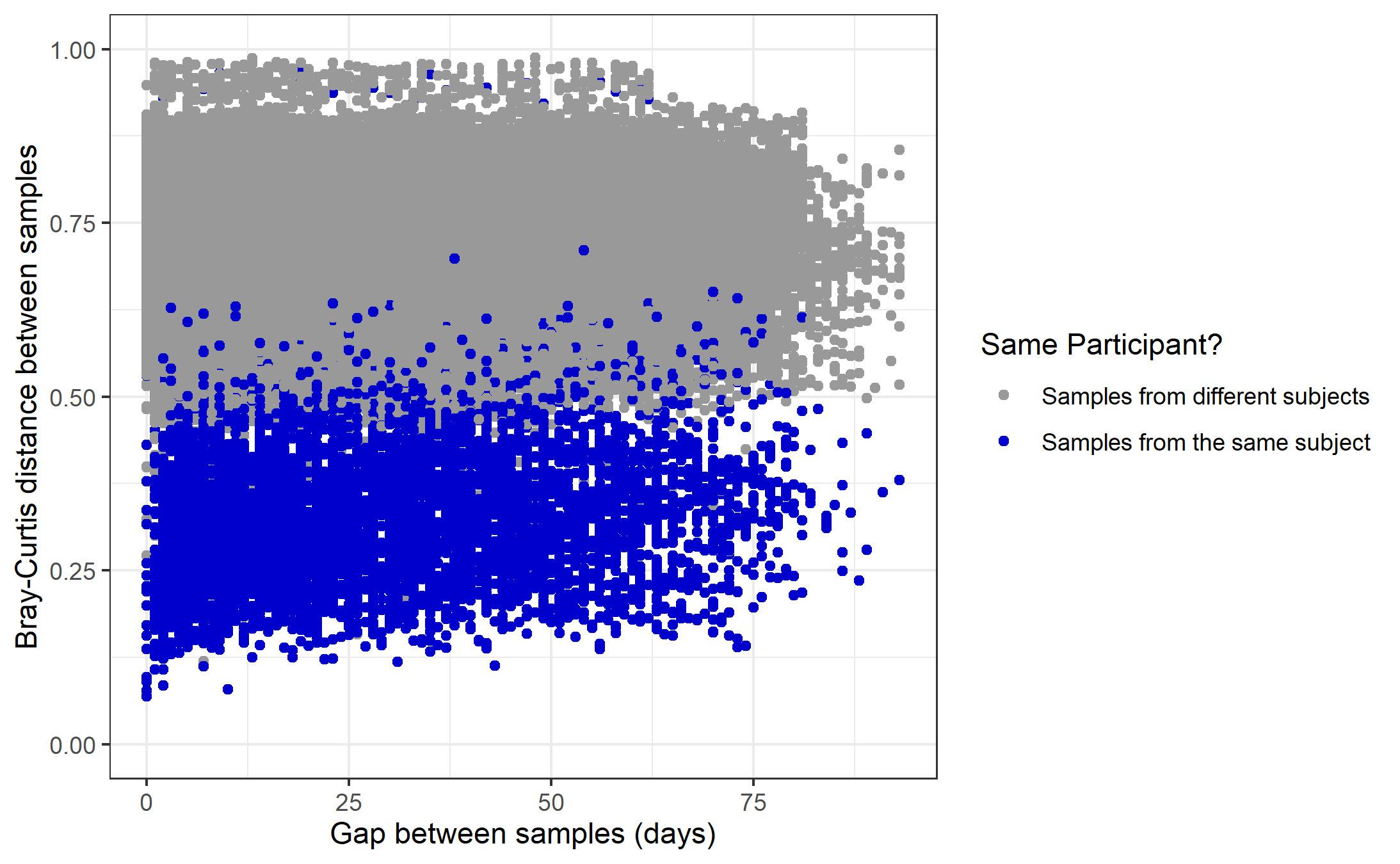

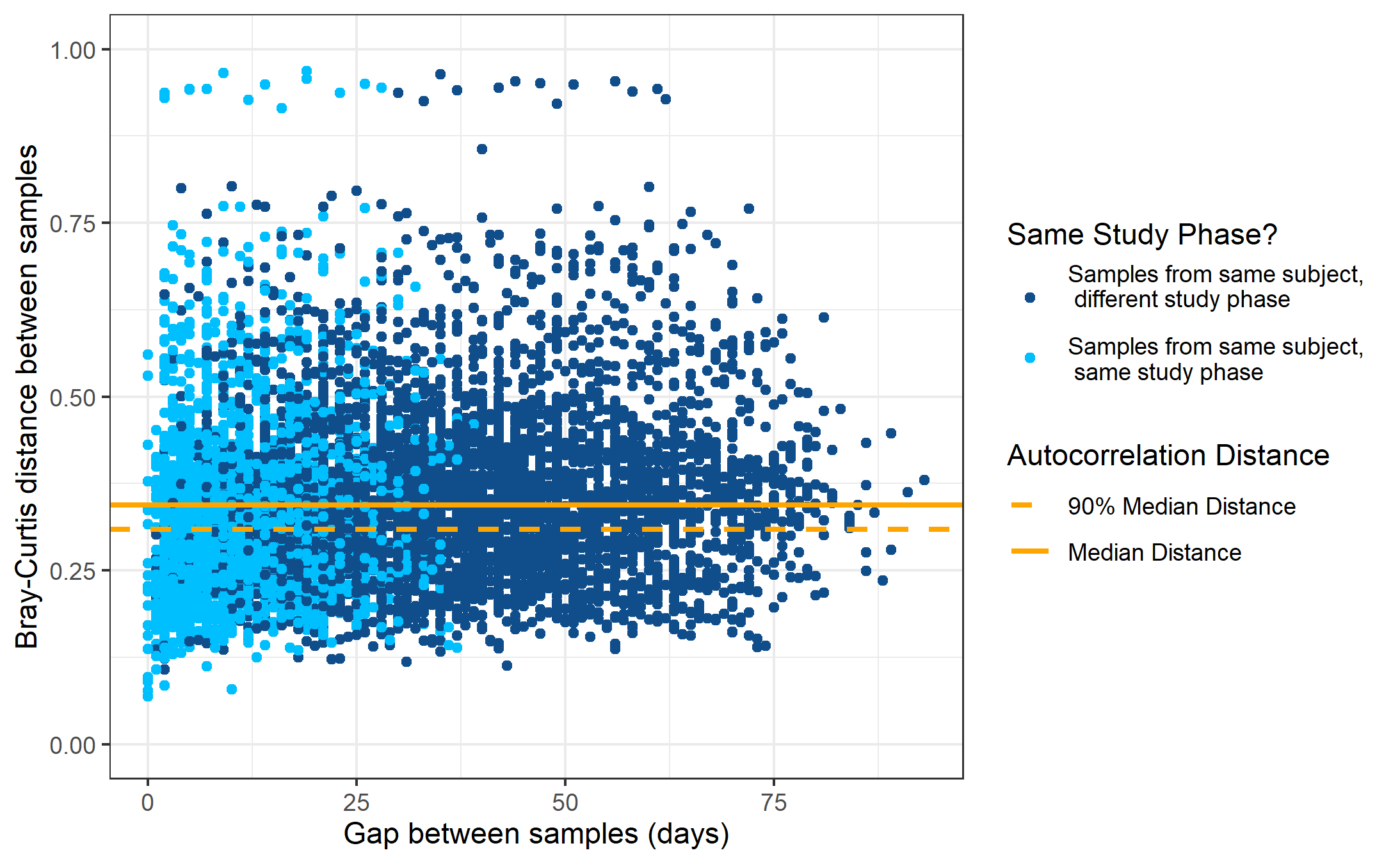

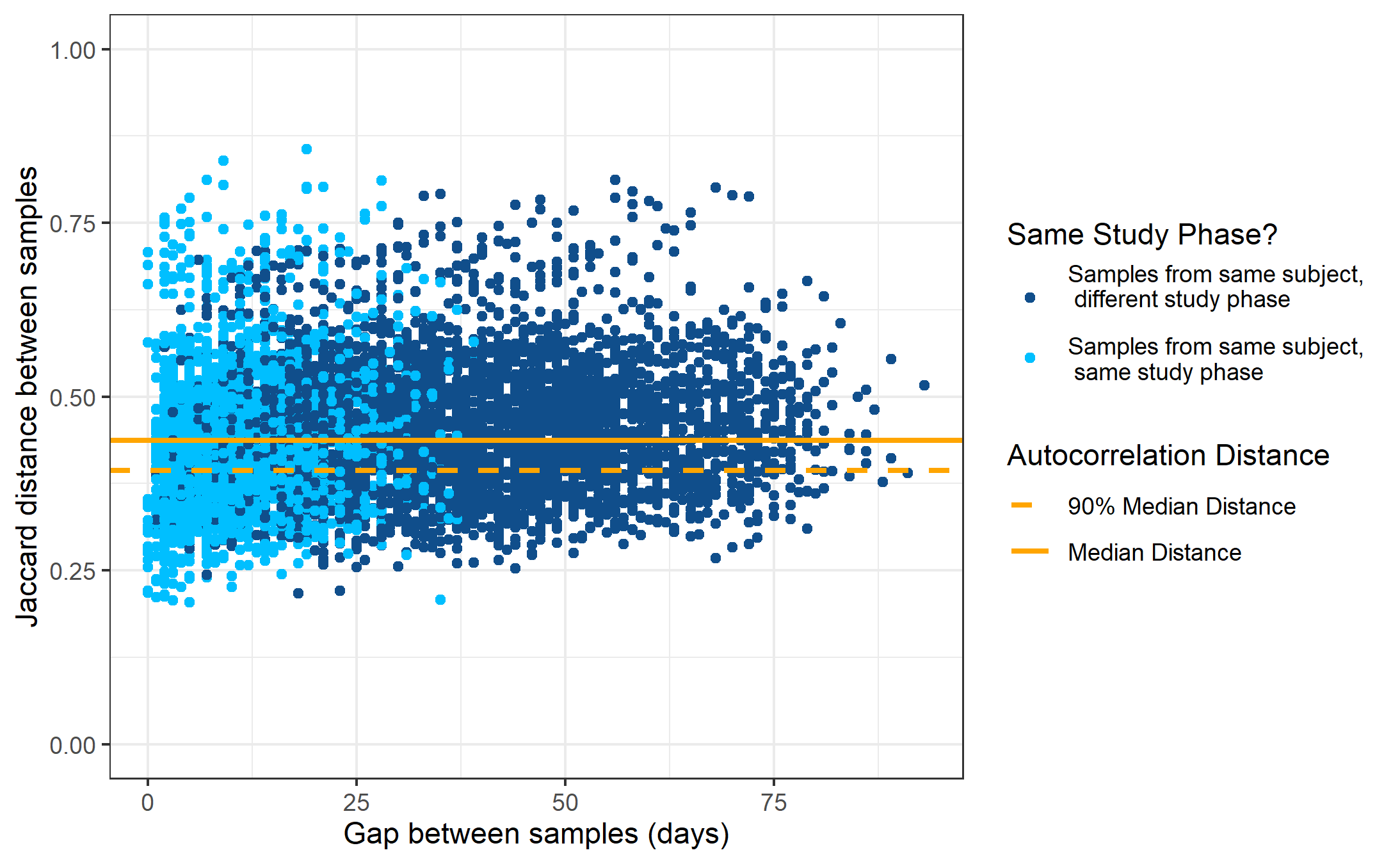

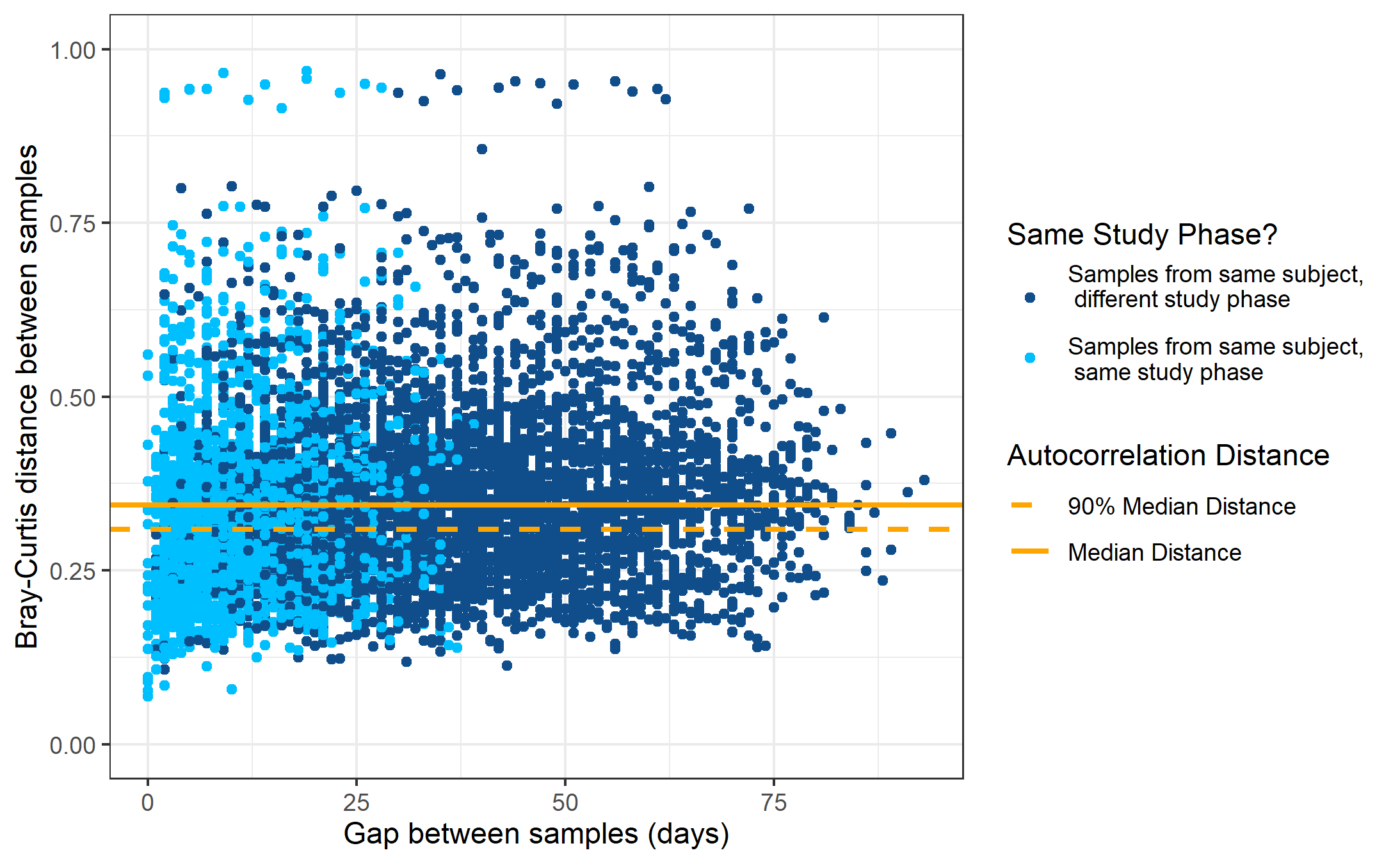
**Supplementary Figure 9.** (a) Bray-Curtis distance between every combination of samples for pairs from the same individual, colored by whether they are from the same study phase or not. (b) Jaccard distance between every combination of samples for pairs from the same individual, colored by whether they are from the same study phase or not. The value at 90% of the median distance between any two randomly chosen samples for each subject is shown along with the median distance at 100%. (c) Bray-Curtis distance between every combination of sample pairs versus the days between when the samples were collected, colored by whether samples in the pair are from the same subject or not. Panel a shows only the subset of samples shown in panel c where both samples are from the same subject. (d) Jaccard distance between every combination of sample pairs versus the days between when the samples were collected, colored by whether samples in the pair are from the same subject or not. Panels a and b show only the subset of samples shown in panels c and d where both samples are from the same subject. Panels a and b show that samples quickly reach the 90% of the median distance; there doesn't seem to be a clear major increase in distance between samples over the course of the study for samples greater in distance than the autocorrelation time. This is surprising because samples in the same experimental phase are closer in time than randomly chosen pairs of samples; thus, one should expect samples within the same phase to be closer in their microbiota structure as the null model, even regardless of the varying regimes of dairy product inclusion in the diet. The fact that temporal clustering ranged in most subjects from limited to none suggests surprisingly high resistance of the microbiota structure to the experimental interventions. Notably, this does not rule out the possibility that some change in microbiota structure occurred in response to the intervention, but rather suggests that these changes are small compared to other sources of variation among samples of the same individual.

(c)

(d)

(b)

(a)

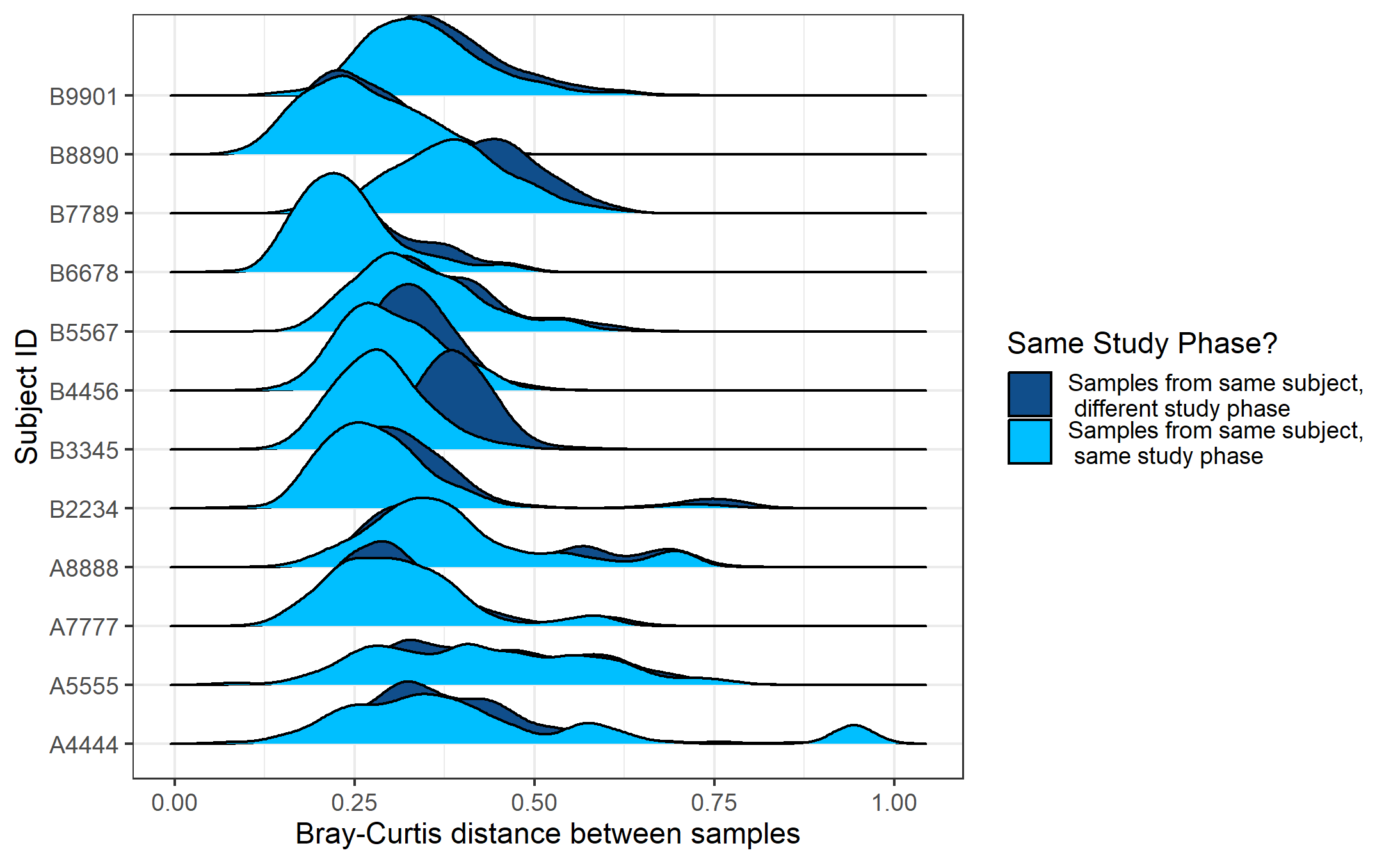

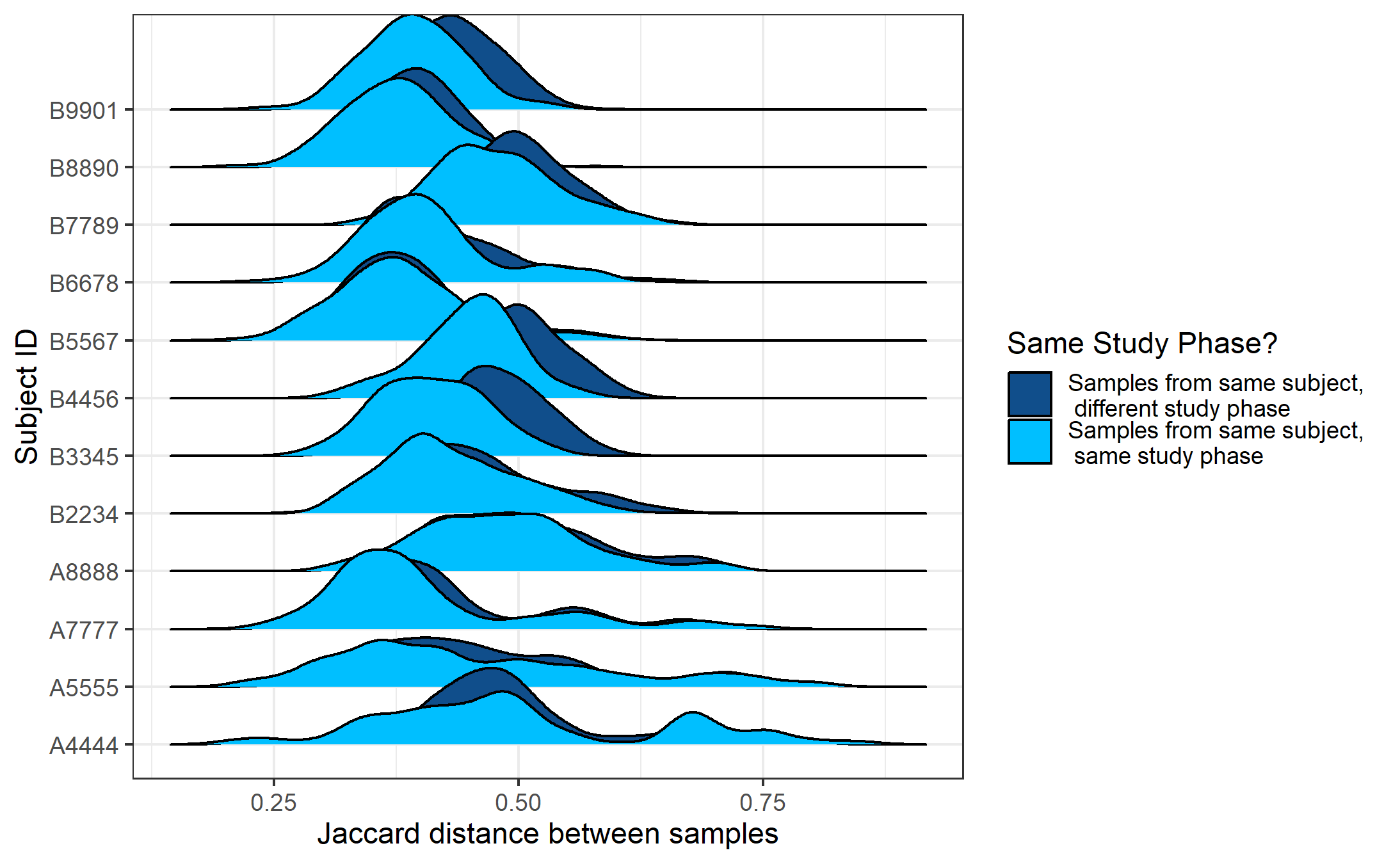

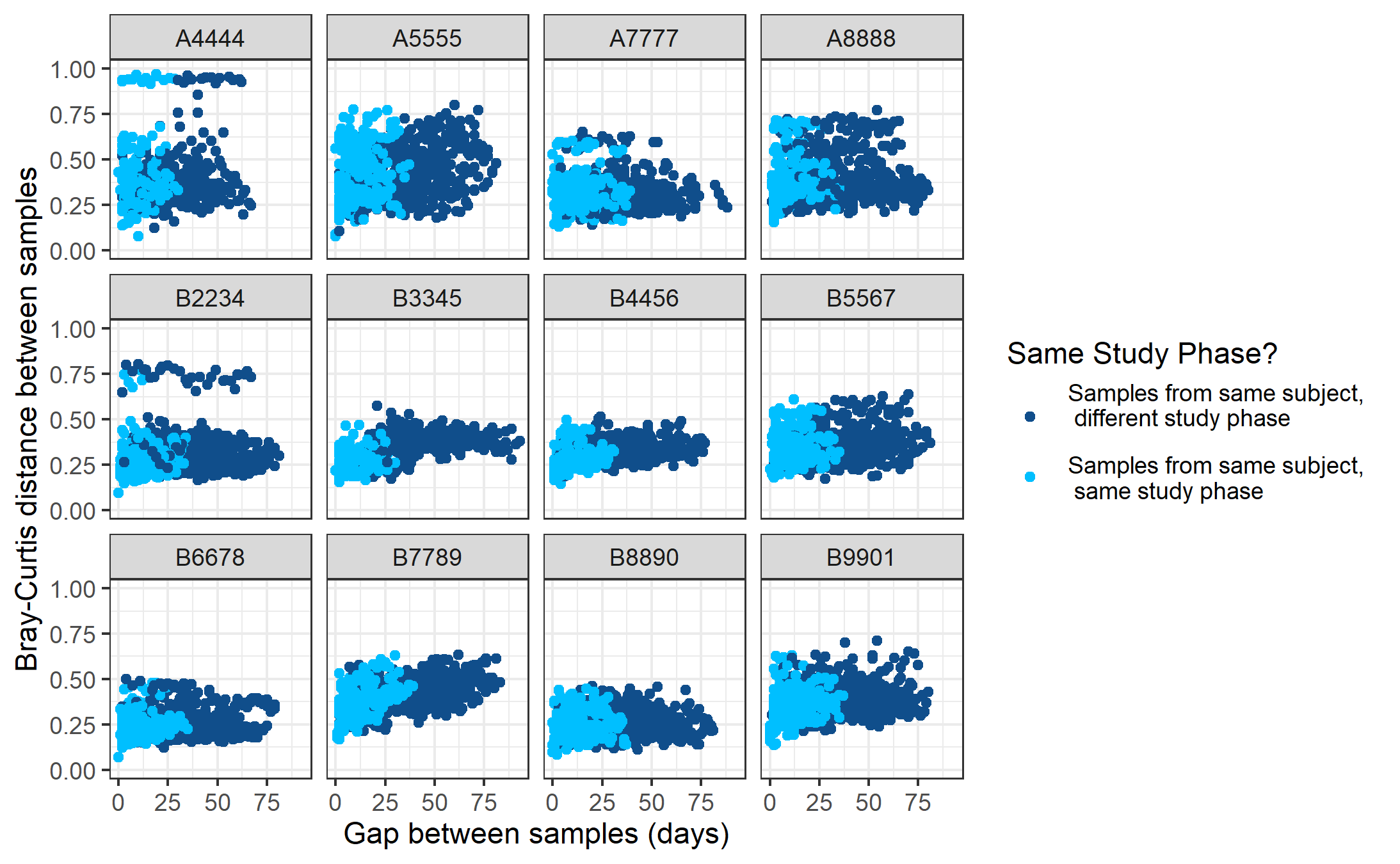

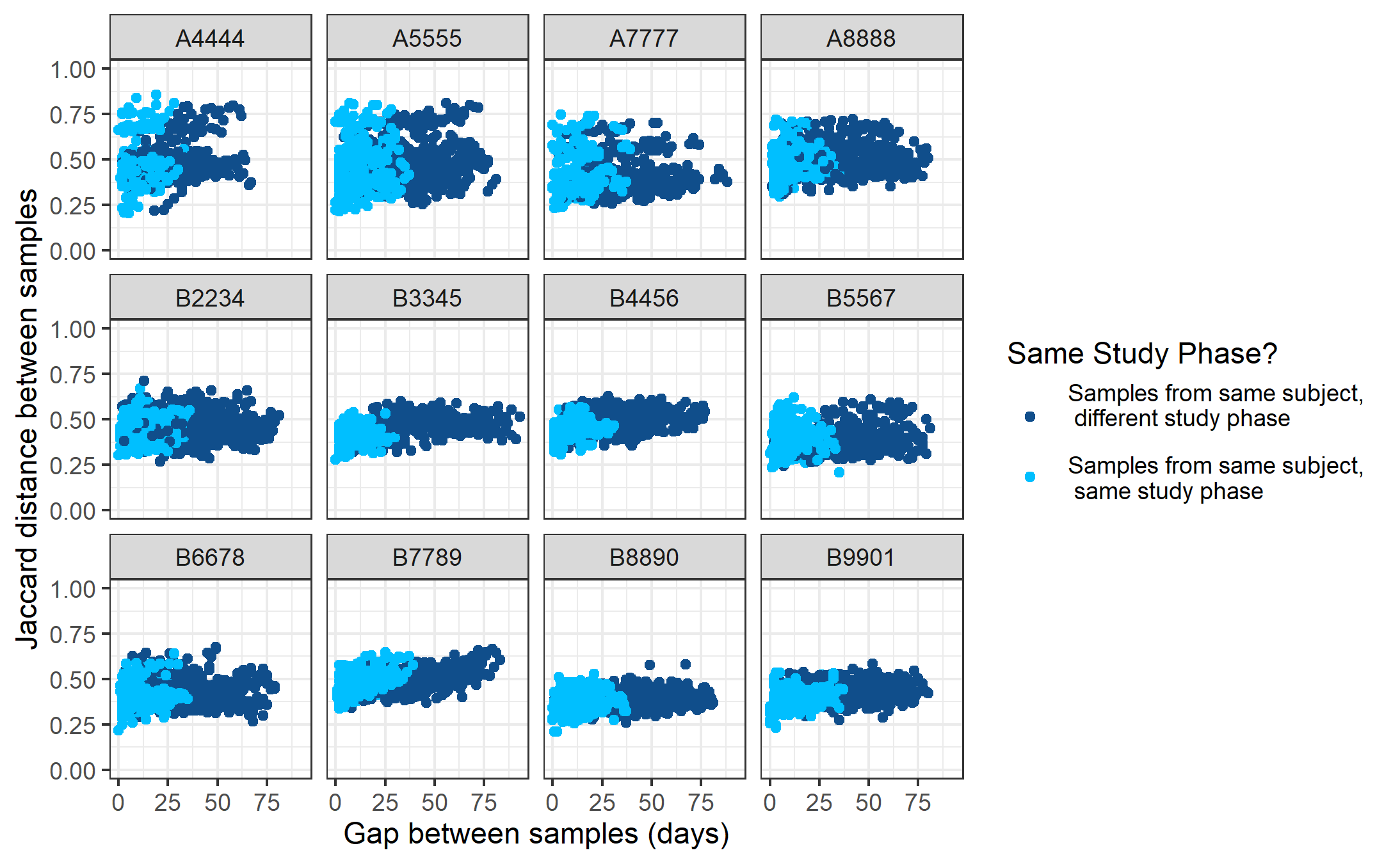

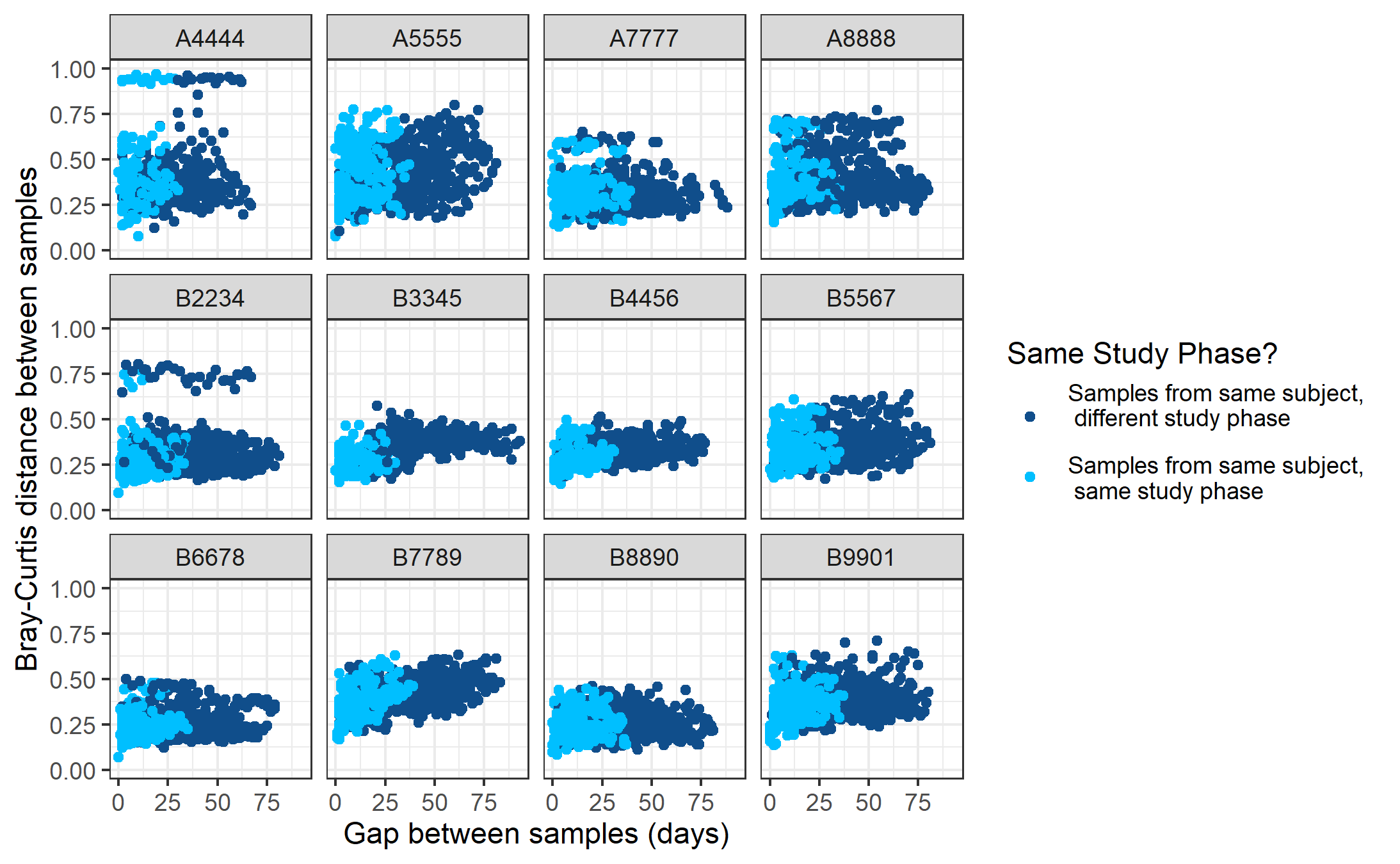
**Supplementary Figure 10.** (a,c) Bray-Curtis and (b,d) Jaccard index distance between every combination of sample pairs where both pairs are from the same individual versus (a,b) the days between when the samples were collected and (c,d) as ridgeline distribution plots. Colored by whether the samples in a given pair are from the same study phase or not. The trends for each subject overall follow relatively similar trends in distribution of distances between pairs of samples at different time gaps between samples through the study.

(a)

(b)

(d)

(c)

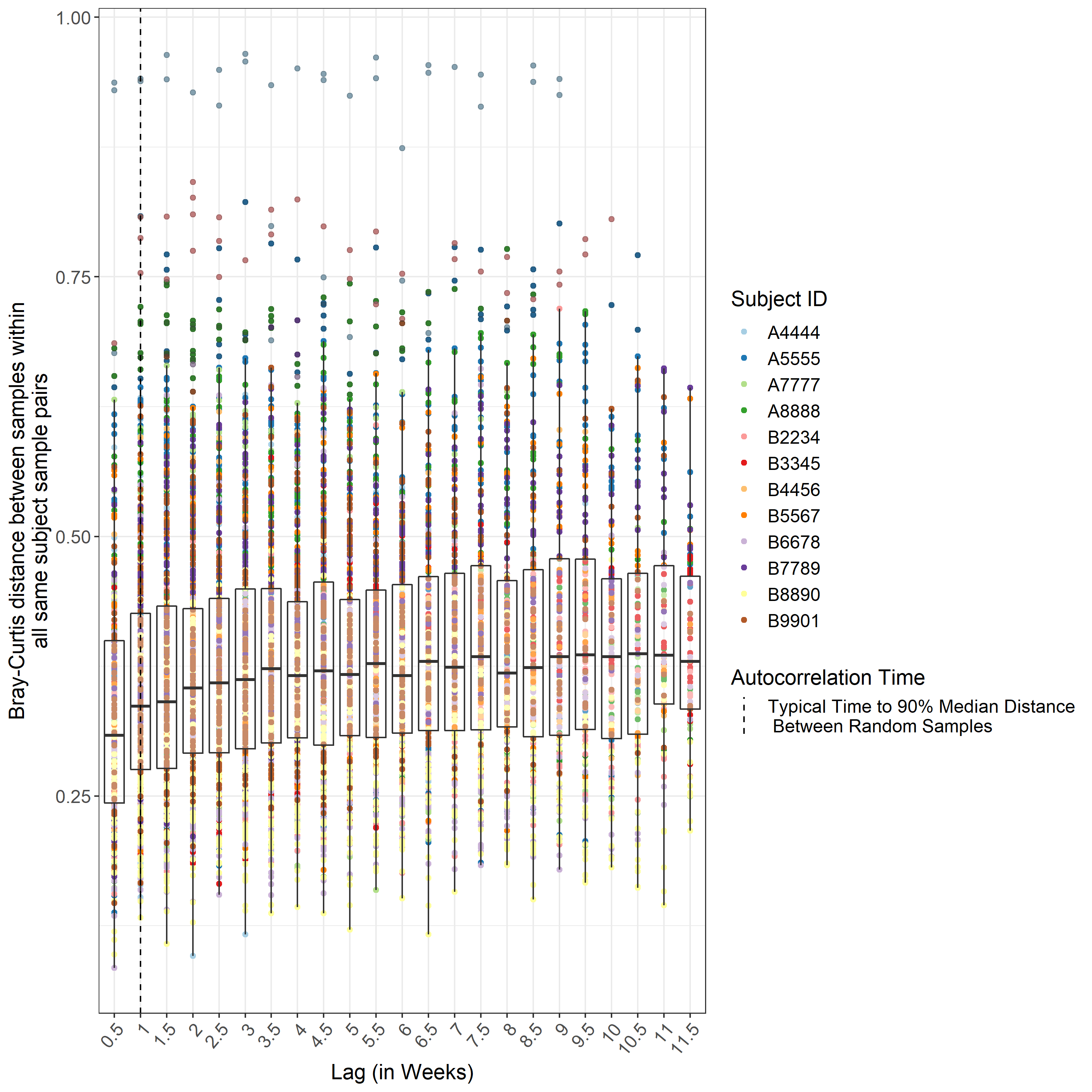

**Supplementary Figure 11.** Bray-Curtis index distance between every sample pair from the same subject throughout the study binning the lag into half week temporal bins. The autocorrelation time using the Bray-Curtis index distance, defined as 90% of the median distance between any two randomly chosen samples where both samples are from the same subject, is 7.00 days.

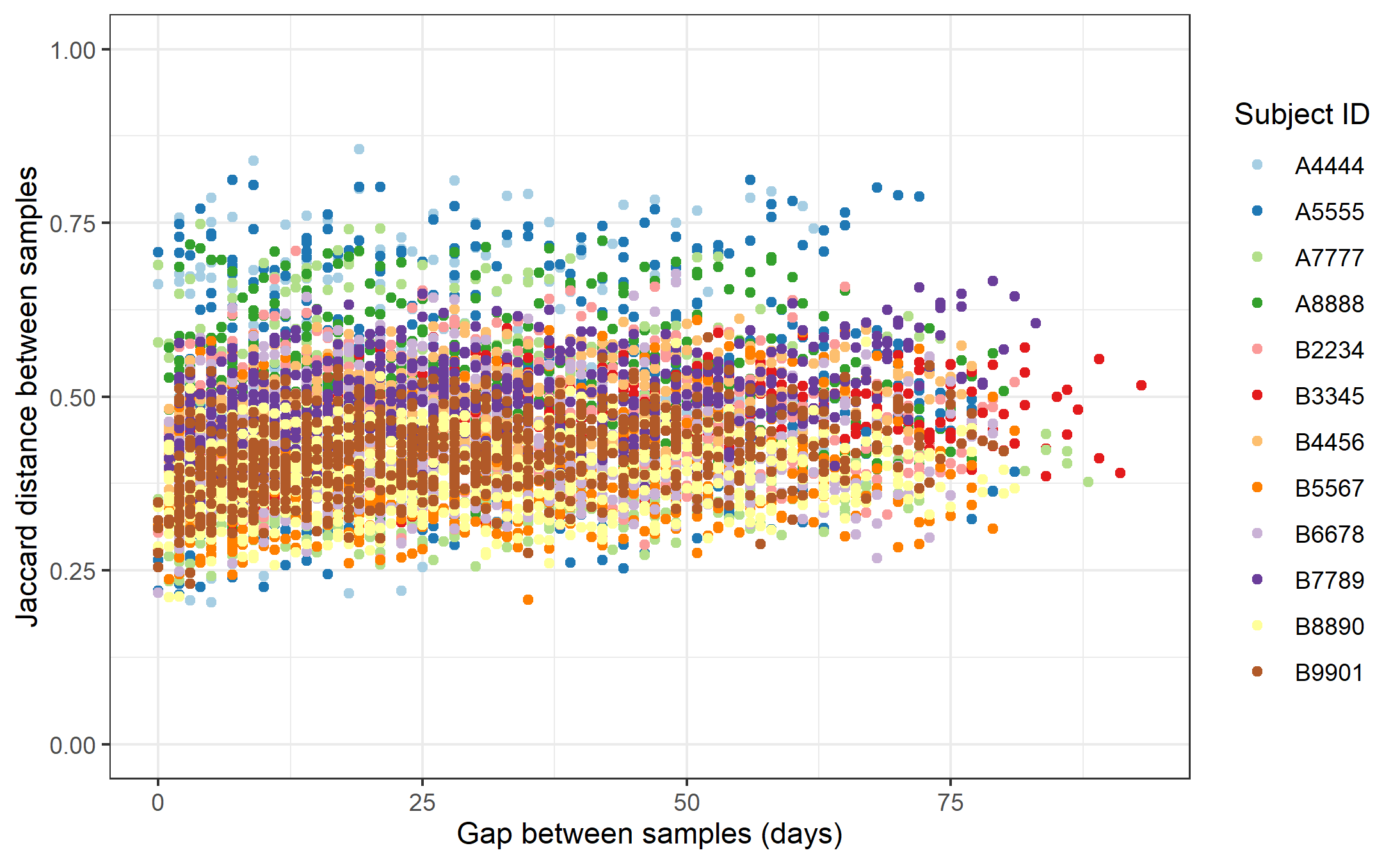

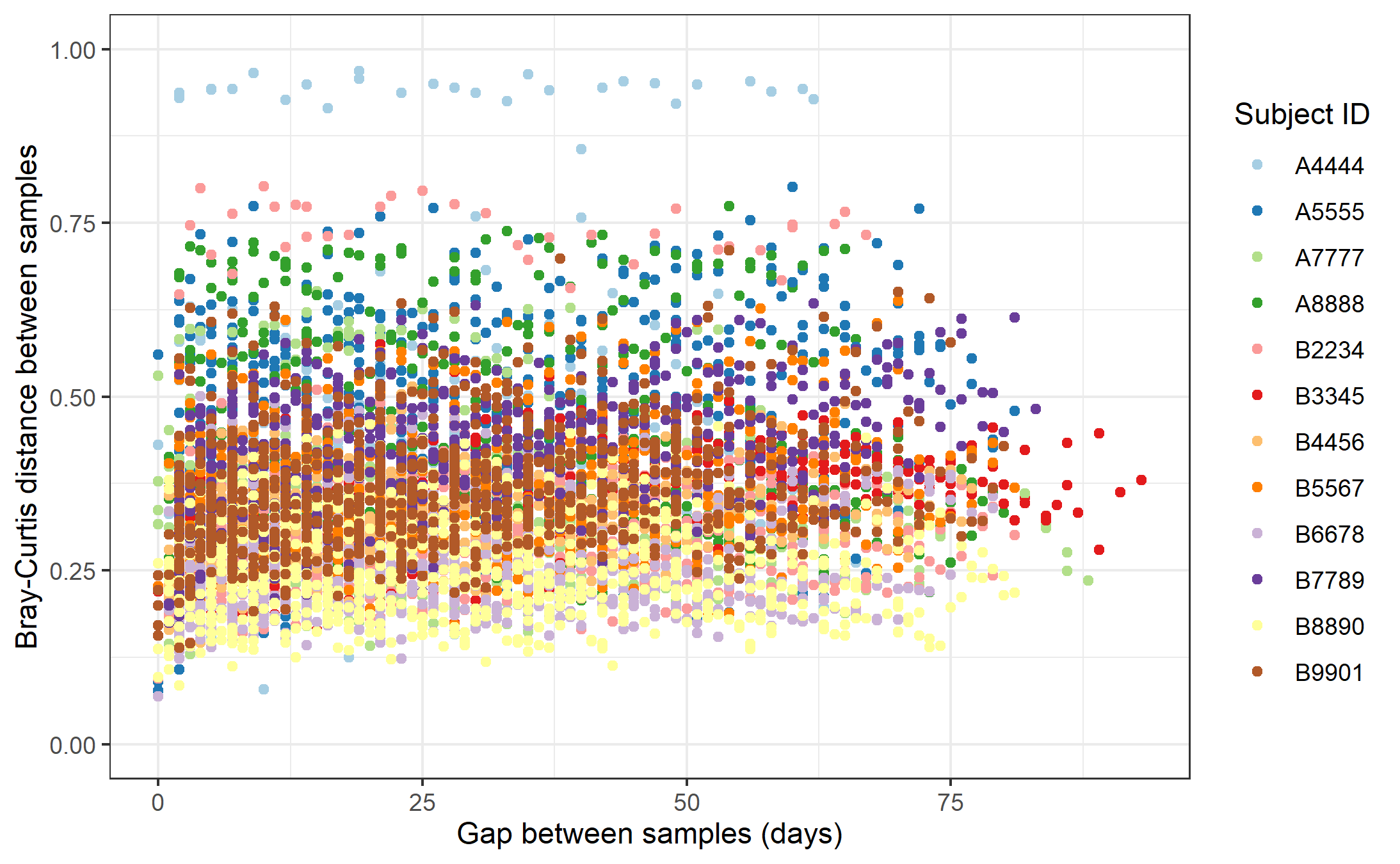
**Supplementary Figure 12.** (a) Bray-Curtis and (b) Jaccard index distance between every combination of sample pairs where both pairs are from the same individual versus the days between when the samples were collected. Colored by which individual the samples are from. Distance between samples where both samples are from the same subject remains relatively consistent across subjects throughout the study.

(b)

(a)

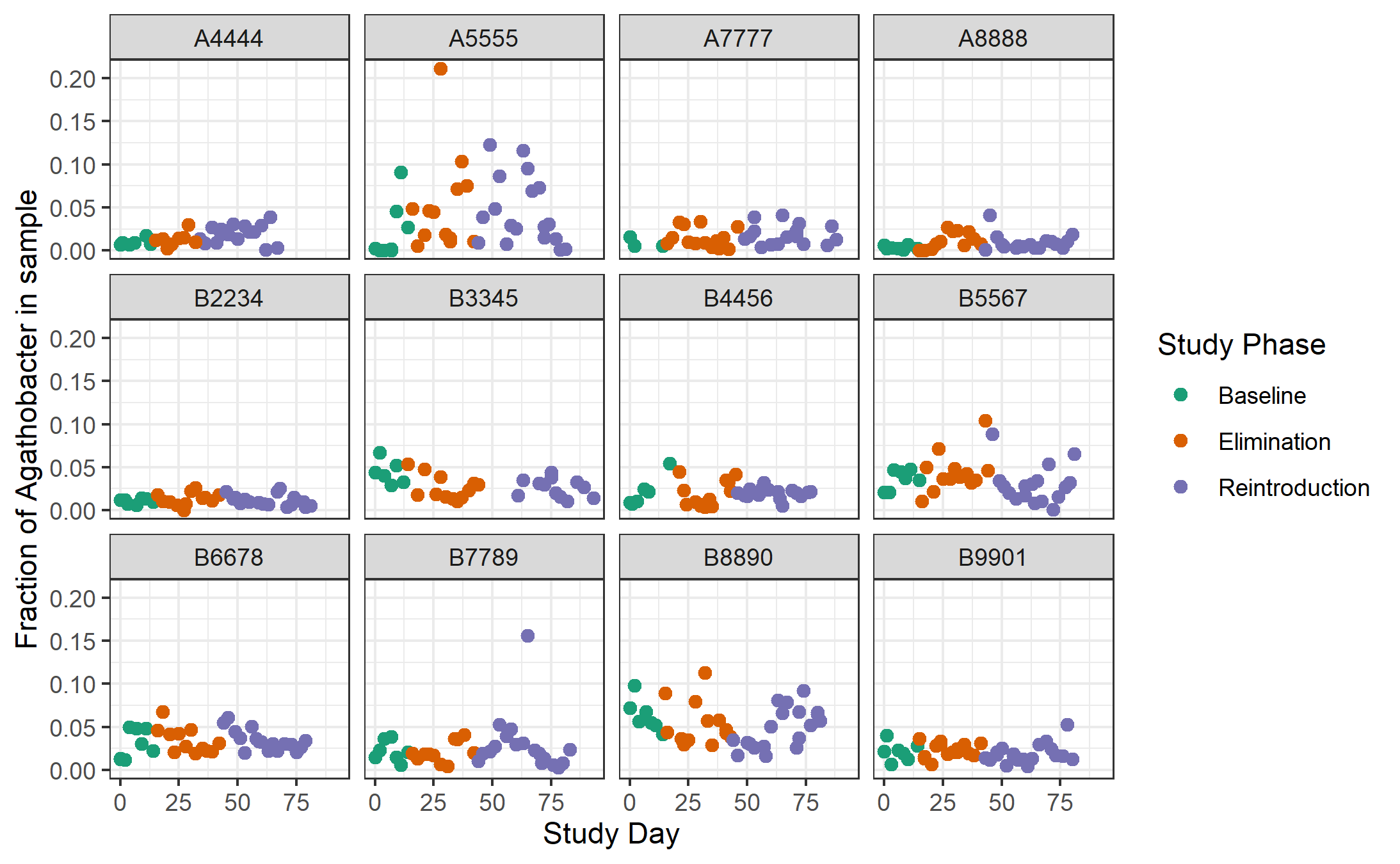

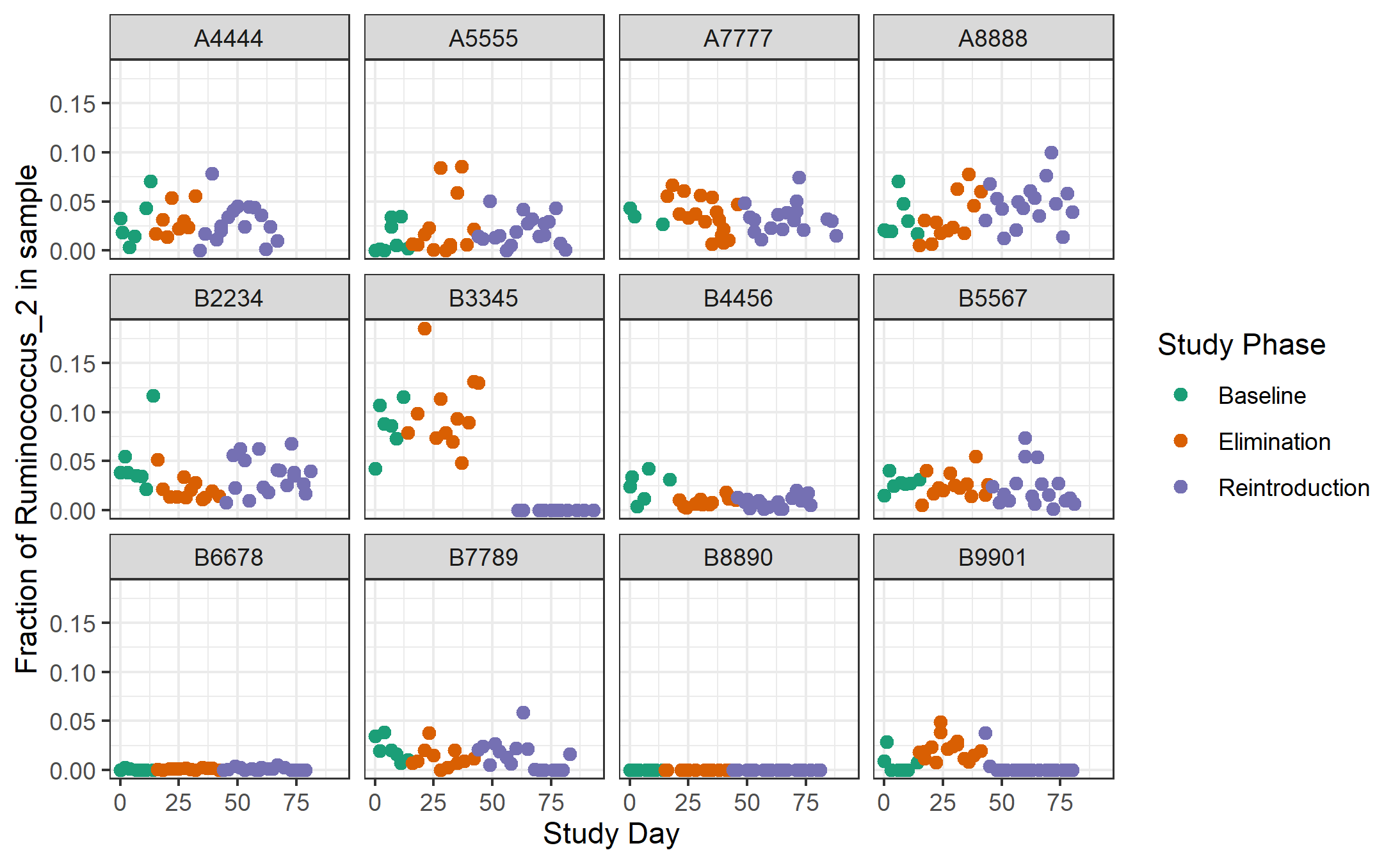
**Supplementary Figure 13.** Abundance of (a) *Ruminococcus_2* genus and (b) *Agathobacter* genus in each sample (normalized based on the total bacterial abundance in that sample) for all subjects over the course of the study, colored by study phase. These genera, along with *Bifidobacterium* encompass many of the ASVs prioritized by treeDA. There was not an obvious consistent shift in the abundance of *Ruminococcus_2 or Agathobacter* in most subjects during either the elimination or reintroduction phase as might be expected. This may indicate that the prioritization from treeDA was a result of overfitting and not anything biological.

(a)

(b)

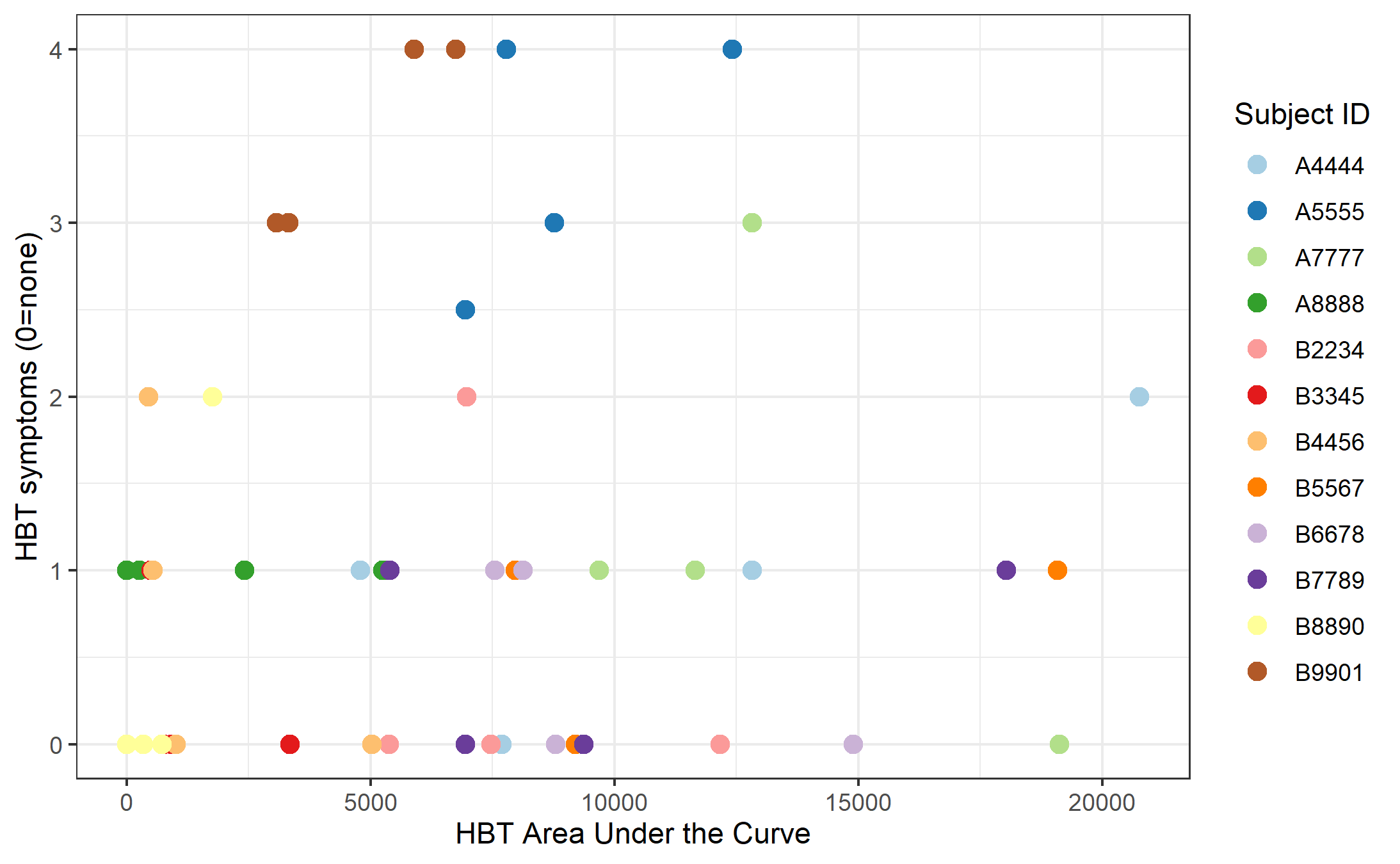
**Supplementary Figure 14.** Rating of self-reported symptoms by subjects (0=no symptoms, 4=severe symptoms such as diarrhea) during each HBT versus the HBT area under the curve of combined concentration for that HBT for each test for all subjects. No significant relationship was found overall (r(44)=0.0915, p=0.541). In addition, we did not observe a clear increase in symptom severity following the elimination phase (Wilcoxon signed rank test p-value=0.85), nor a statistically significant decrease in symptoms across subjects following the reintroduction phase to baseline (Wilcoxon signed rank test p-value=0.42) as might have been expected from the increase in HBT area under the curve observed in **Figure 2**. This may be attributable to the diverse range of possible symptoms and the challenge of quantifying them in a consistent and comparable manner across subjects, coupled with the relatively small number of subjects in the study. In addition, reports of symptom intensity can be subjective and can be influenced by subject perception, such as a placebo of expecting more symptoms the first HBT after not having had two cups of milk in a long time. In fact, the change in reported symptoms from the 1^st^ to 2^nd^ HBT was significant (Wilcoxon signed rank test p-value = 0.036) and could be reflecting this placebo effect.

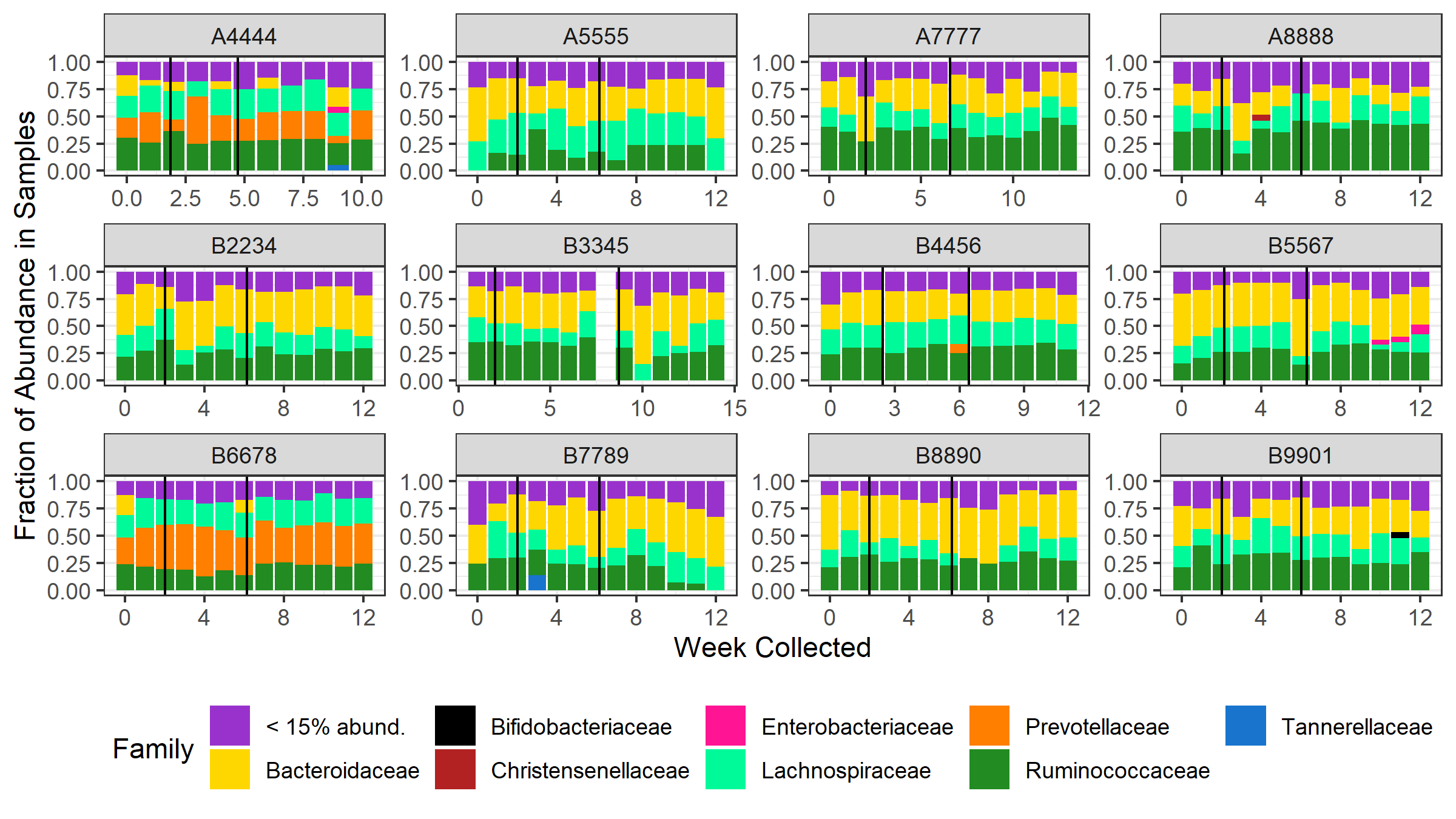

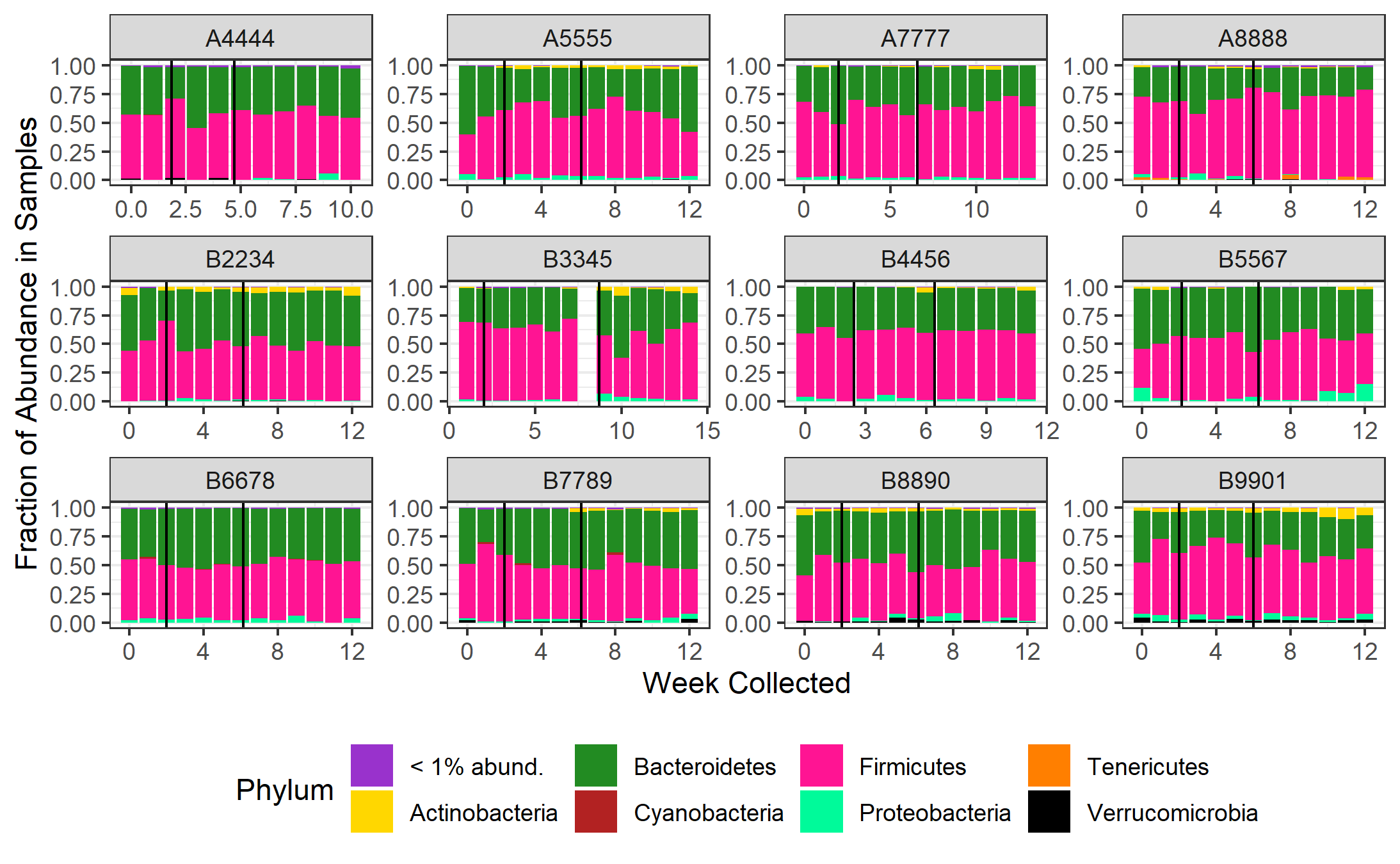

(b)

(a)

**Supplementary Figure 15.** Average abundance of top (a) Family and (b) Phyla for each subject for each week. Black vertical lines represent change in study phase from baseline to elimination and then elimination to reintroduction. Most subjects did not experience a dramatic and consistent shift in the distribution of their top taxa throughout the study and no consistent trend was observed across subjects.

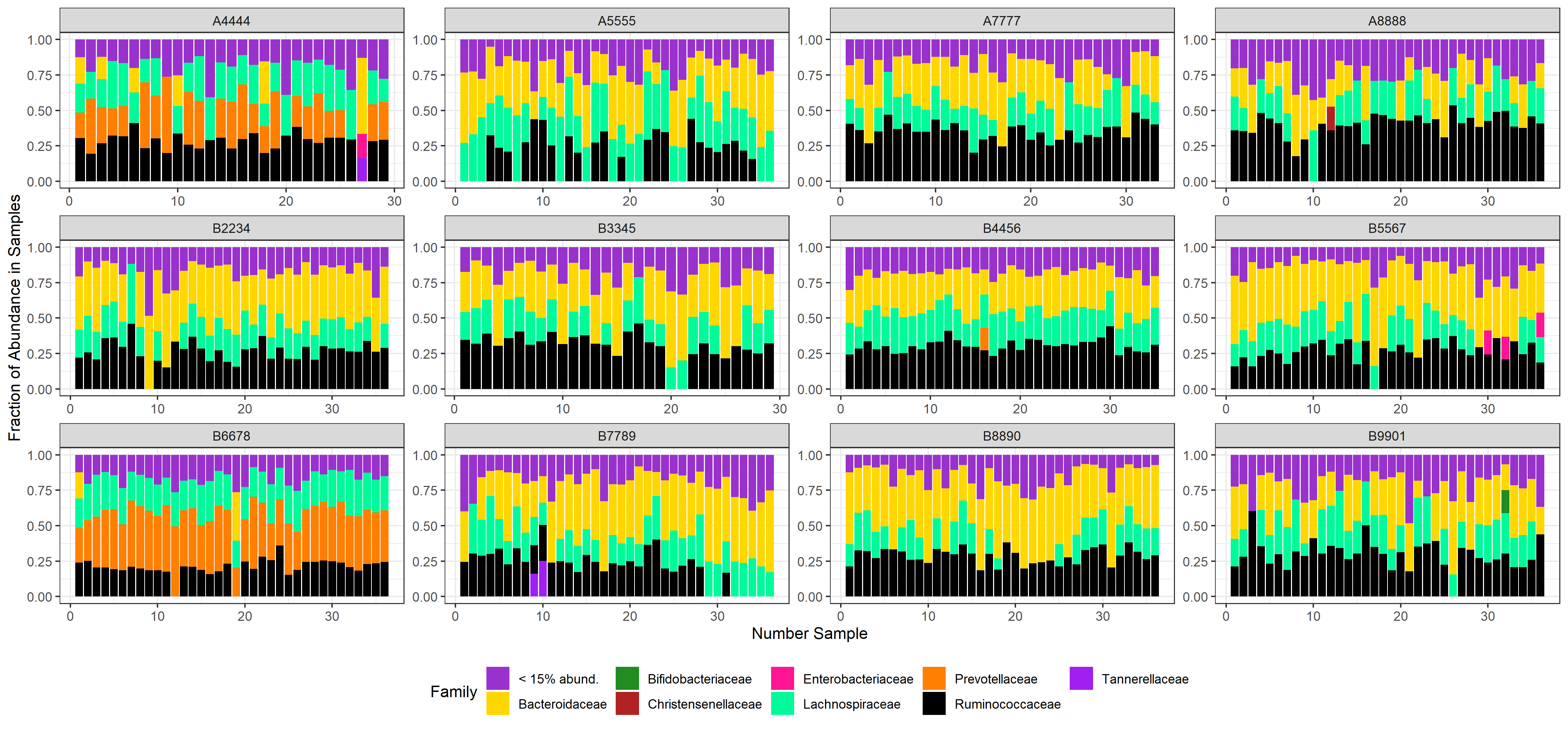

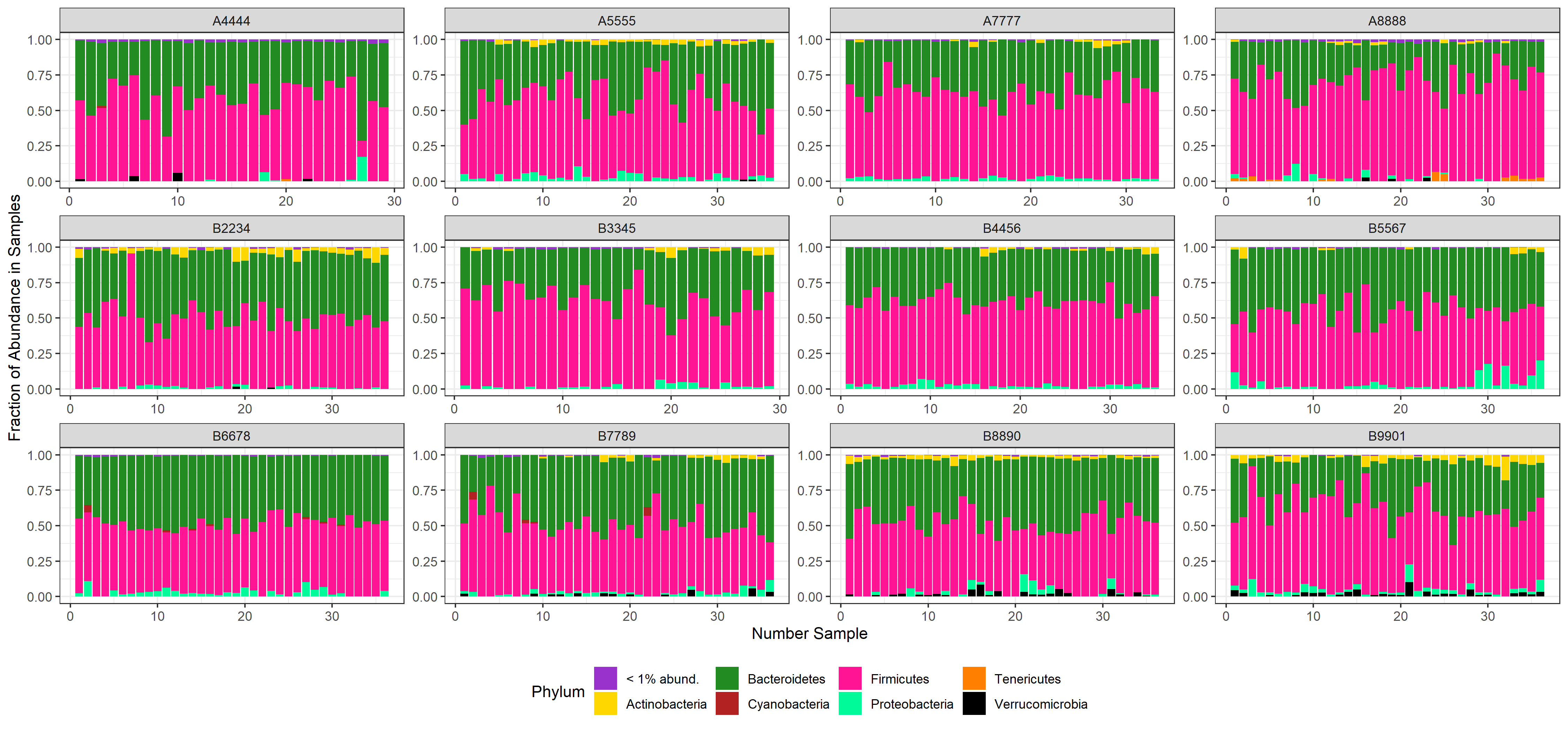
**Supplementary Figure 16.** Average abundance of top (a) Family and (b) Phyla for each subject for each consecutive sample throughout the study. Most subjects did not experience a dramatic and consistent shift in the distribution of their top taxa throughout the study and no consistent trend was observed across subjects.

(a)

(b)

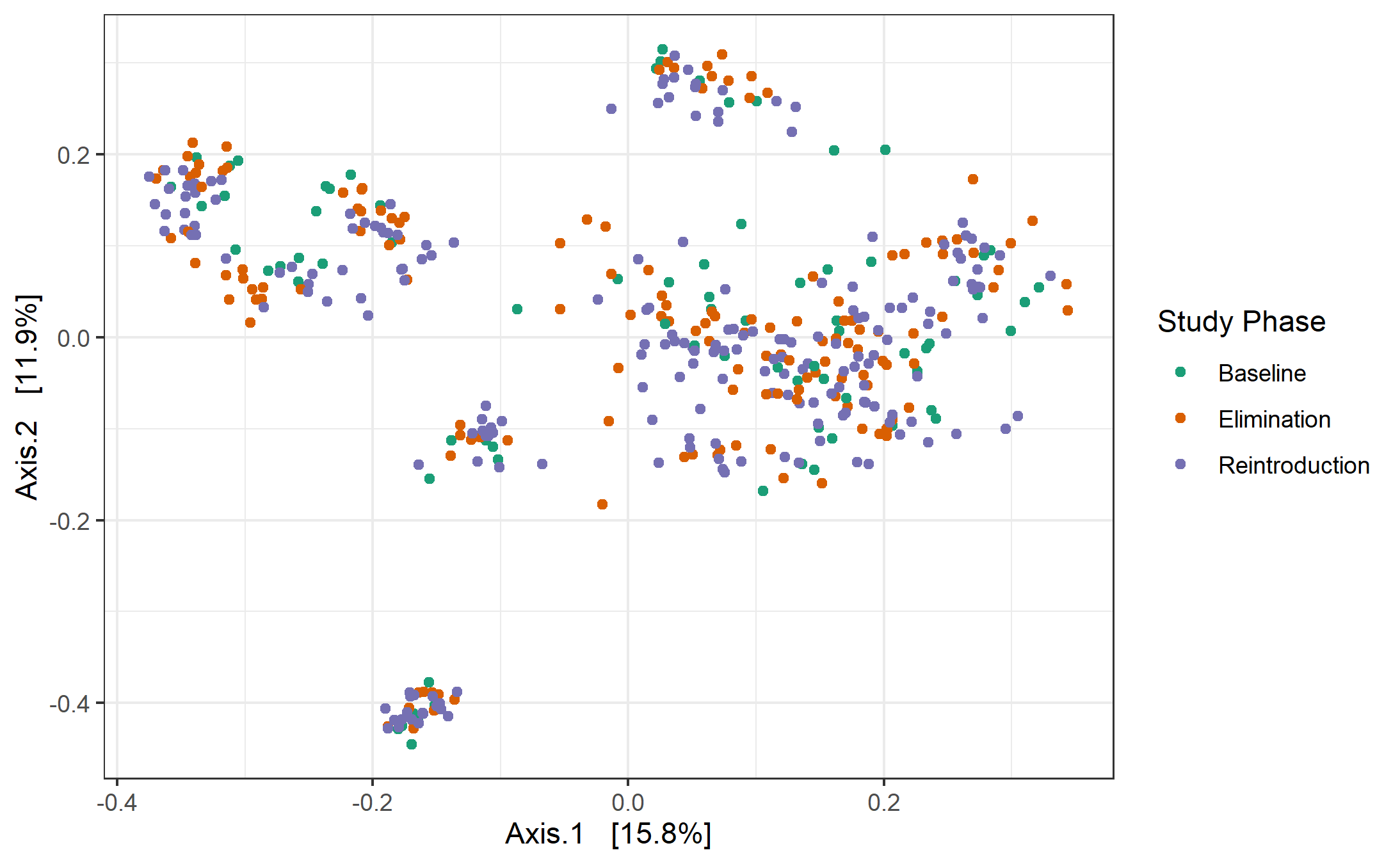

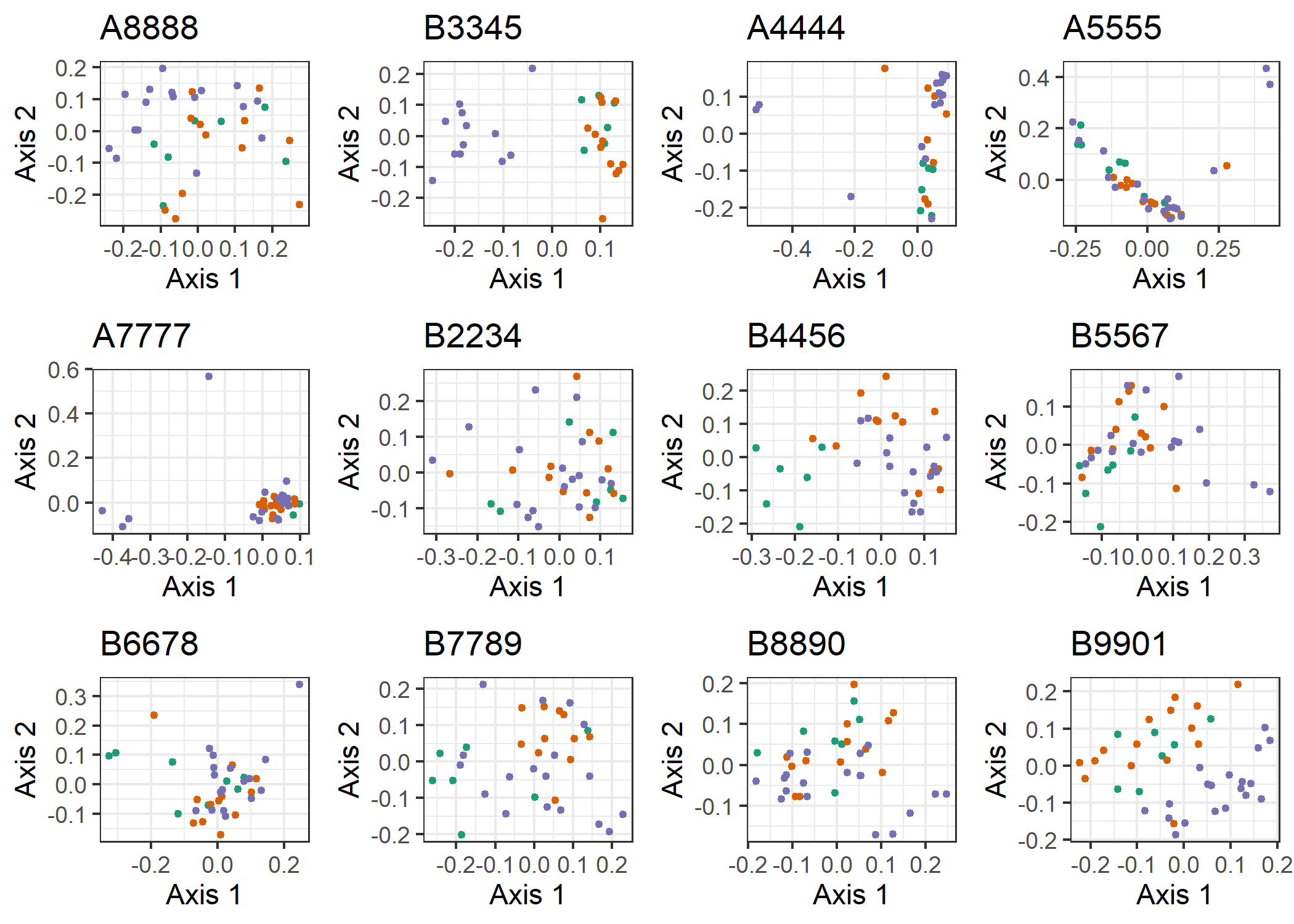

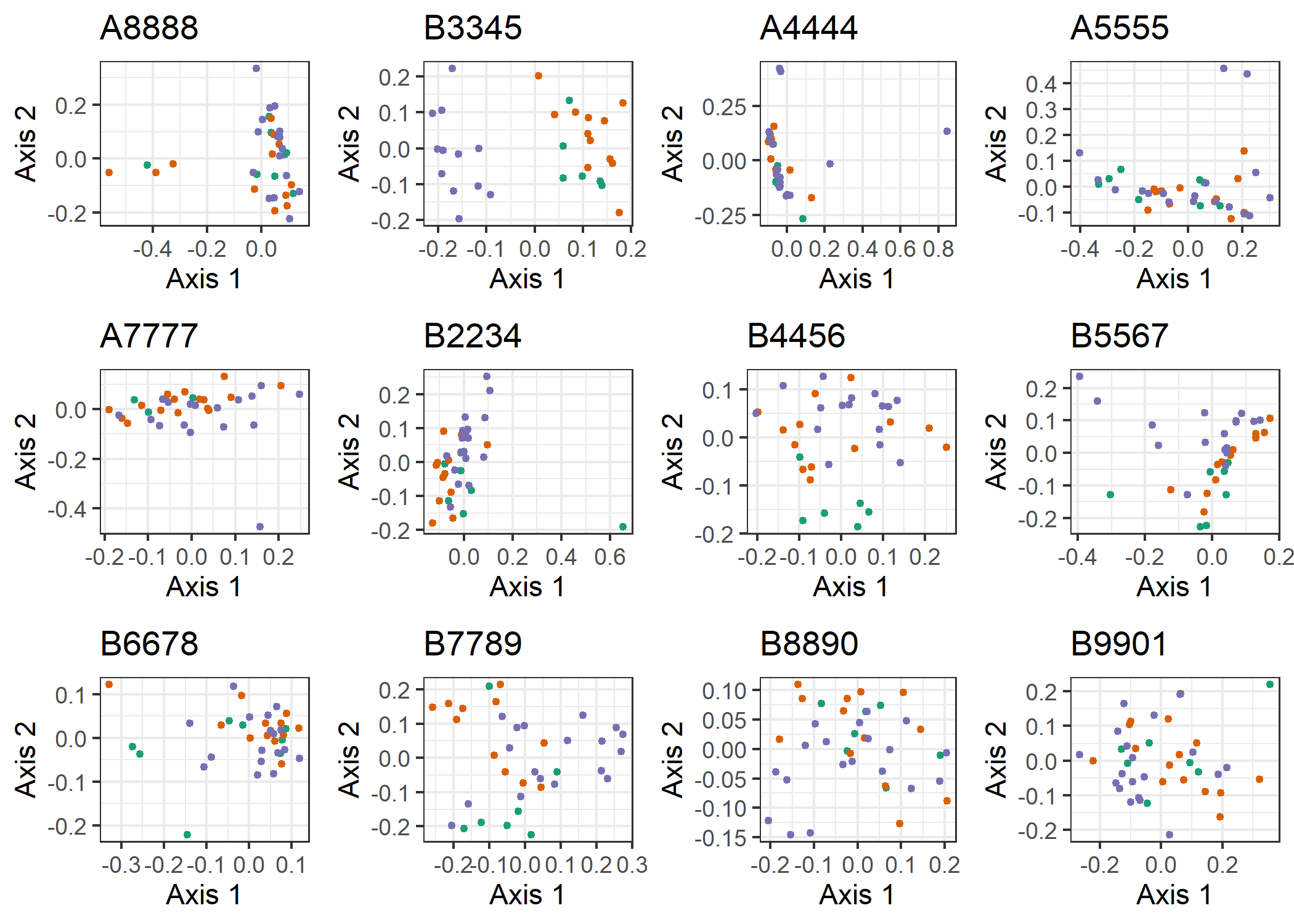
**Supplementary Figure 17.** Visualization of the first two principal coordinates from PCoA across all samples for each subject individually using (a) Bray-Curtis and (b) Jaccard index distance metrics, colored by study phase. Most samples for many subjects showed minimal signs of clustering by phase, if at all.

(a)

(b)
